# Amygdala Persistence in Aging: Replication and Extension of Puccetti et al. (2021)

**DOI:** 10.64898/2026.10.14.759196

**Authors:** Adam Turnbull, Meishan Ai, James J. Gross, Feng Vankee-Lin

## Abstract

Age-related differences in affect are well established, but their neural mechanisms remain unclear. In a preregistered replication of Puccetti et al. (2021) and extension, we examined age, affect, left-amygdala negative persistence, and amygdala–ventromedial prefrontal cortex (vmPFC) task-based connectivity using MIDUS 3 daily diary and fMRI data. Older age predicted higher positive and lower negative affect in the daily diary sample (*N* = 1,168; *ps* < .001). In the primary diary + fMRI sample (*N* = 81; ages 48–95, 61.7% female, 82.7% White), greater persistence predicted higher raw negative affect (β = 0.278, 95% CI [0.012, 0.544], *p* = .020); associations with positive and log-transformed negative affect were nonsignificant (*ps* ≥ .063). Older age predicted lower persistence in the final fMRI sample (*N* = 127; β = −0.00260, 95% CI [−0.00504, −0.00016], *p* = .018), but this association was weaker in the primary diary + fMRI sample (*p* = .200), with no evidence of mediation. Greater anterior vmPFC connectivity predicted lower raw negative affect (β = −0.106, 95% CI [−0.223, 0.012], *p* = .039) and log-transformed negative affect (*p* = .014). In MIDUS Refresher 1, affect estimates were directionally similar but nonsignificant (*N* = 48; *ps* ≥ .081), whereas the age–persistence estimate reversed direction. These findings provide partial support for amygdala persistence and task-based connectivity as neural correlates of daily affect, while highlighting variability in the magnitude and replicability of fMRI–behavior associations.

**Public Significance Statement:** Among middle-aged and older adults, lingering brain responses to negative images and communication between emotion-related brain regions were associated with negative feelings in daily life. However, the findings did not establish why emotional experience differs with age, and evidence from a second sample was inconclusive. Larger studies are needed to determine how reliably these brain measures relate to everyday emotional well-being.

## Introduction

Older adults consistently report more positive affect and less negative affect than younger adults (Mather, 2012). Despite this, there are significant individual differences in affective experience in older adults. Positive affective experience has been highlighted as a key resilience factor for promoting healthy aging (Anthony et al., 2024) while a shift toward negative affect is associated with the emergence of late-life psychiatric diagnoses (Alexopoulos, 2019) as well as increased risk for dementia (Hu et al., 2020). There is a need to better characterize the neurocognitive circuit-level mechanisms of affective experience in healthy older adults to better understand these individual differences.

Task-based fMRI is a promising tool for understanding the circuit-level mechanisms underlying affective experience in healthy older adults. Recent perspectives suggest that task-based fMRI can better probe at-risk processes in older adults by highlighting key differences that only emerge when these processes are engaged (Mooraj et al., 2025). Previous research has highlighted that both the activity of the amygdala and vmPFC-amygdala connectivity in response to negative affective experiences are important for adaptive emotional responding. Initially, this connectivity was conceptualized as a form of model-free or implicit emotion regulation (Etkin et al., 2015), although more recent models frame this in terms of adjusting or modulating affective valuations encoded in the amygdala (Turnbull et al., 2026; Zhang et al., 2025), with anterior vmPFC hypothesized to preferentially adjust towards safety while posterior vmPFC adjusts towards threat (Tashjian et al., 2021).

To test whether vmPFC-amygdala circuitry was associated with age-related differences in daily life affect we leveraged the recent release of the Midlife in the US (MIDUS) (https://midus.wisc.edu/) 3 Neuroscience project (MIDUS3; 2017-2022). Specifically, we performed a pre-registered replication and extension of a study linking increased persistence in the voxel-wise pattern of left amygdala responses to negative images to daily life affect in the MIDUS 2 dataset (Puccetti et al., 2021). Pre-registration and replication are important strategies to reduce type 1 error, particularly in neuroimaging analyses in which there are increased researcher degrees of freedom (Botvinik-Nezer et al., 2020; Carp, 2012) and known inflation of effect sizes (Marek et al., 2022).

We tested a set of pre-registered hypotheses (**Table 1**). First, we aimed to replicate the established findings that older adults report improved daily life affect (Mather, 2012) and greater amygdala persistence following negative images relates to worse daily life affect (Puccetti et al., 2021). We then tested two confirmatory extensions: that amygdala persistence is reduced in older adults and that this reduced persistence statistically mediates age-related improvements in affect. We then tested the pre-registered exploratory hypothesis that the engagement of vmPFC-amygdala task-based connectivity predicts amygdala persistence and daily life affect (see **Figure 1**). Finally, we explored whether these relationships are attenuated by habitual use of reappraisal or suppression (reflecting compensation via model-based emotion regulation).

**Figure 1.**
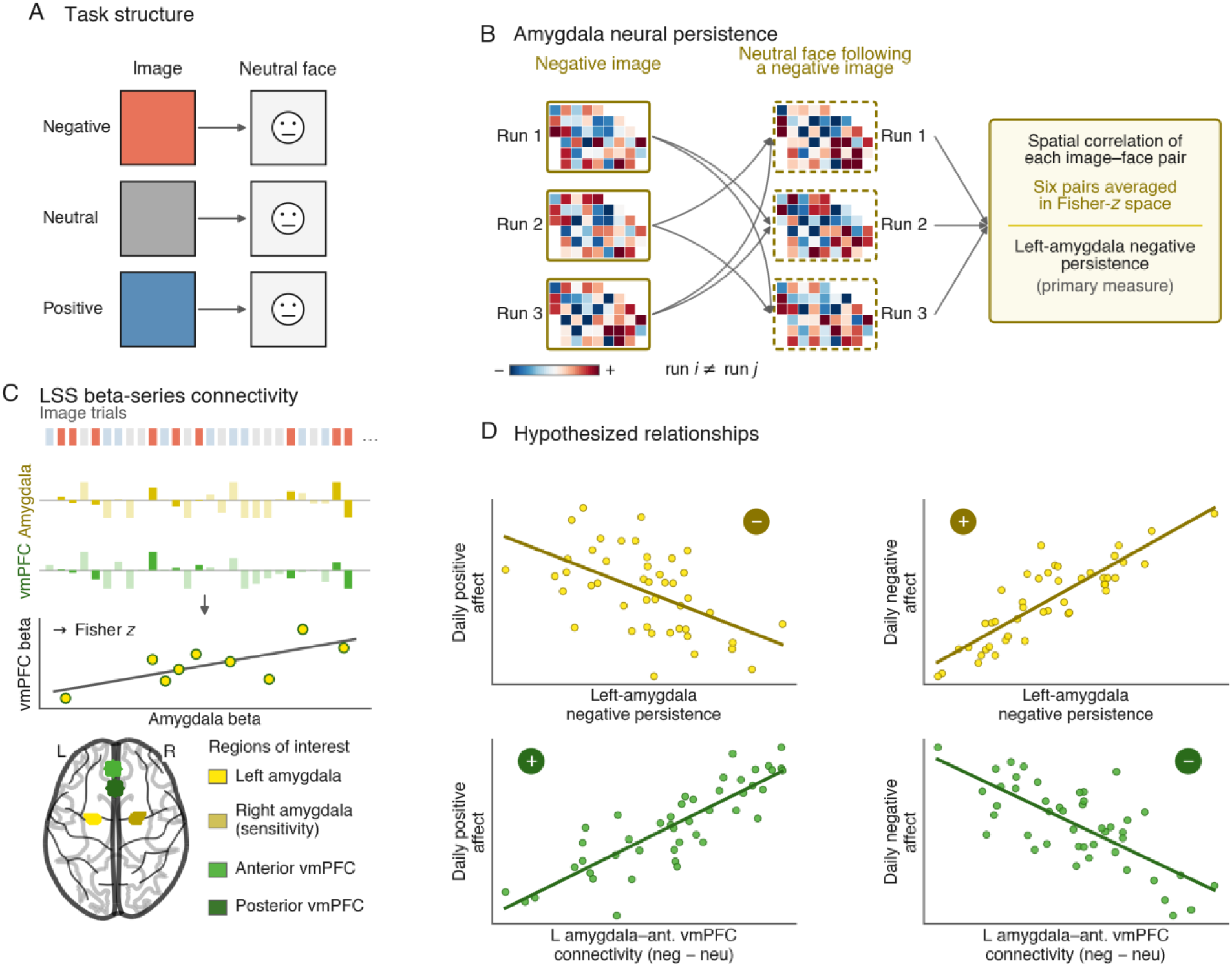
Task design, neural measures, and hypothesized relationships. (A) Negative, neutral, and positive images were each followed by a neutral face. (*B*) Left-amygdala negative persistence was the mean Fisher-*z*-transformed spatial correlation across six directional cross-run negative-image/following-neutral-face pattern pairs. (C) Least-squares-separate beta-series connectivity correlated trial-level amygdala and vmPFC responses within run and valence; negative-minus-neutral connectivity was the primary contrast. The inset shows bilateral amygdala and anterior/posterior vmPFC regions of interest. (D) Hypothesized directions relating persistence and left-amygdala–anterior vmPFC connectivity to daily positive and negative affect. Schematic values are illustrative.

**Figure 2.**
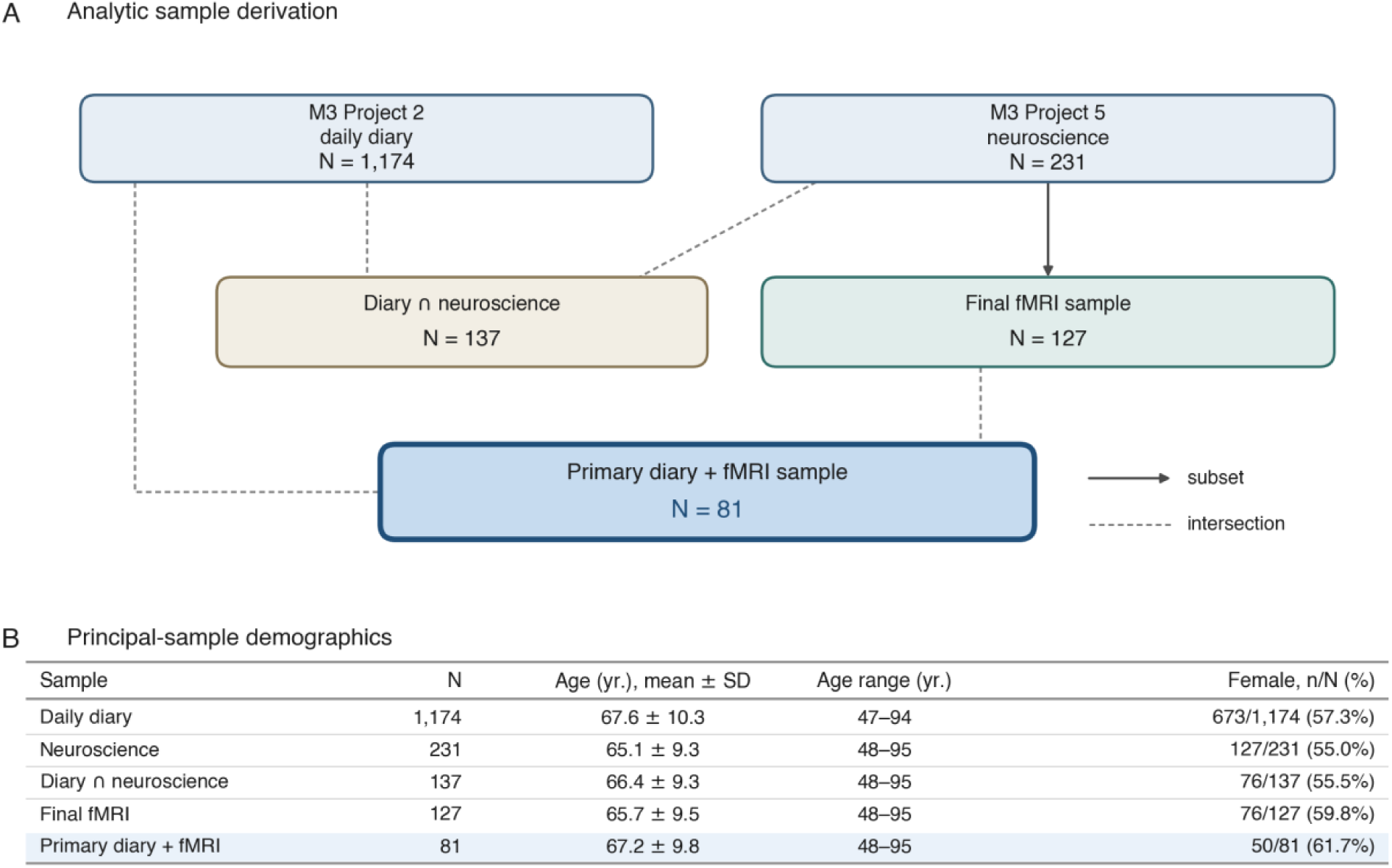
Analytic sample derivation and demographics. (A) MIDUS 3 Project 2 diary and Project 5 neuroscience samples and their intersections. Solid arrows indicate subsets; dashed lines indicate intersections. (*B*) Demographic characteristics of the principal analytic samples.

**Figure 3.**
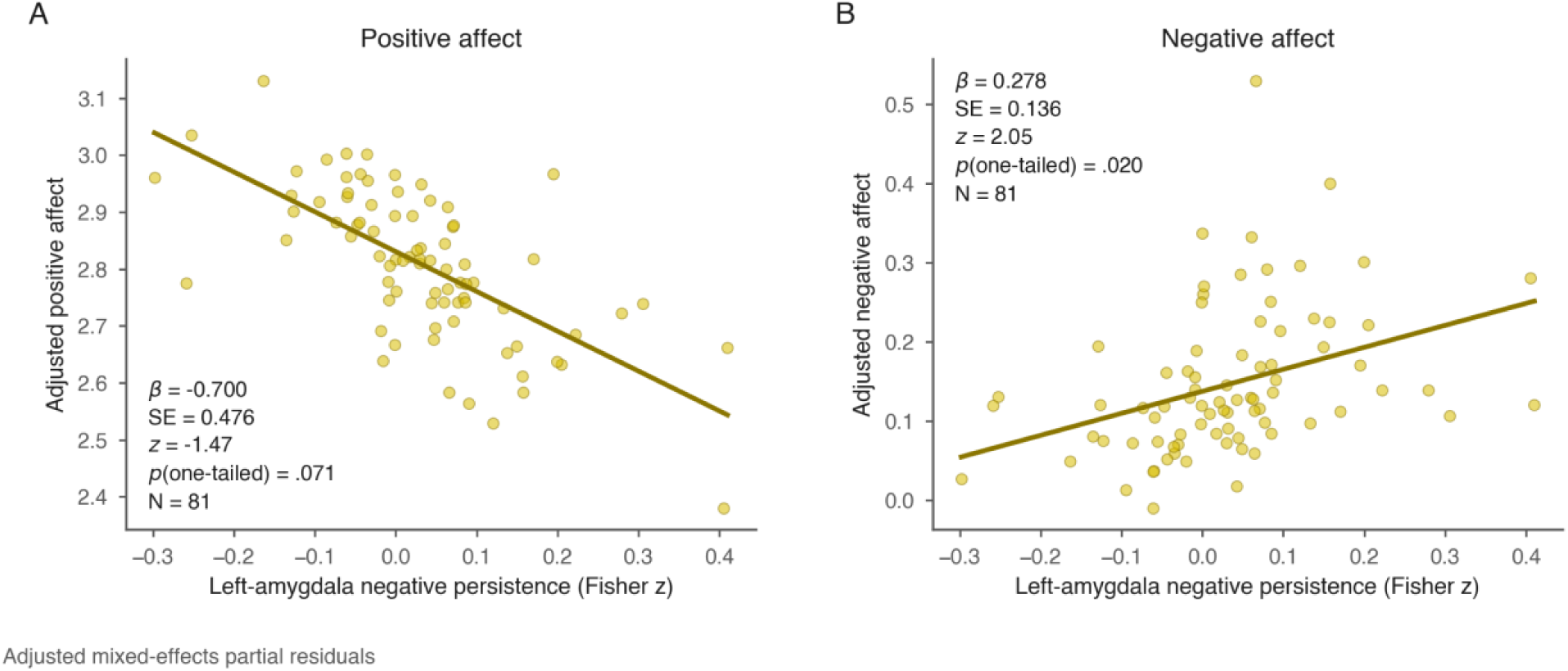
Left-amygdala negative persistence and daily affect. Mixed-effects partial residuals show adjusted associations with (A) positive affect and (*B*) raw negative affect in the primary diary + fMRI sample (*N* = 81). Lines show fitted fixed-effect associations.

**Figure 4.**
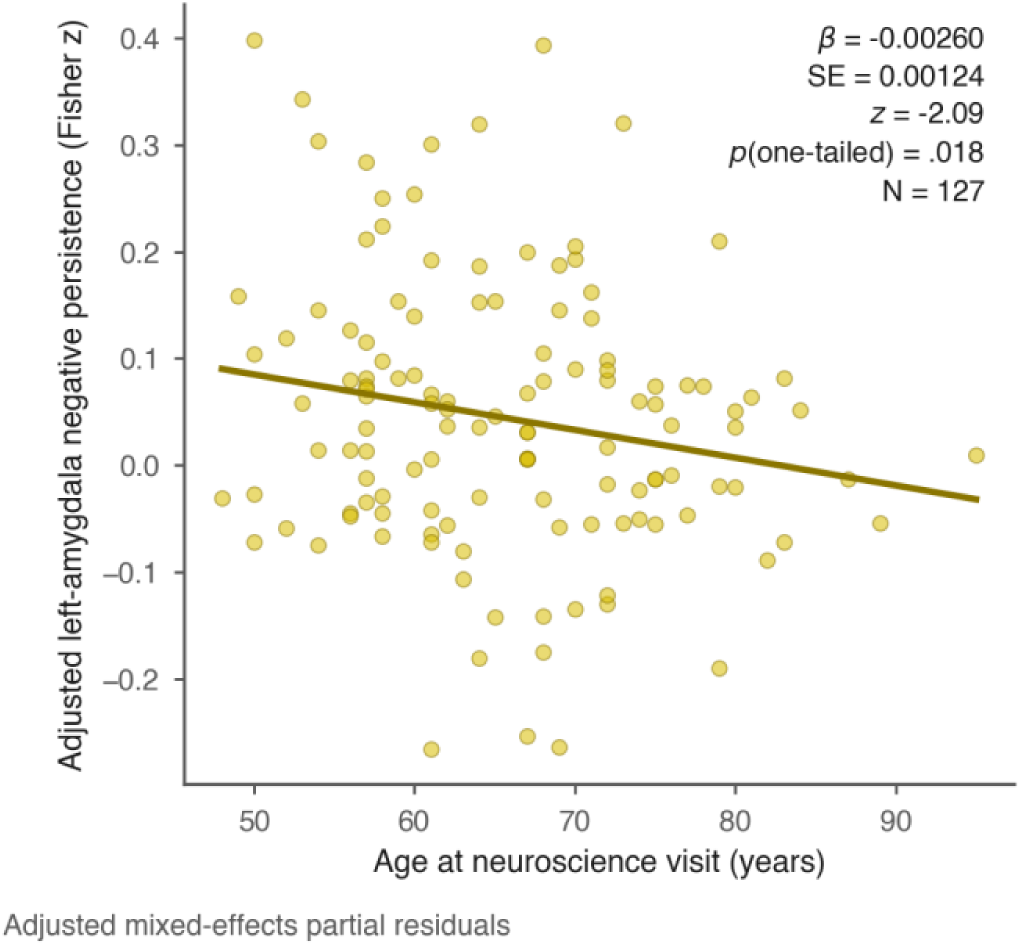
Age and left-amygdala negative persistence. Mixed-effects partial residuals show the adjusted association between age at the neuroscience visit and persistence in the final fMRI sample (*N* = 127); the line shows the fitted fixed-effect association.

**Figure 5.**
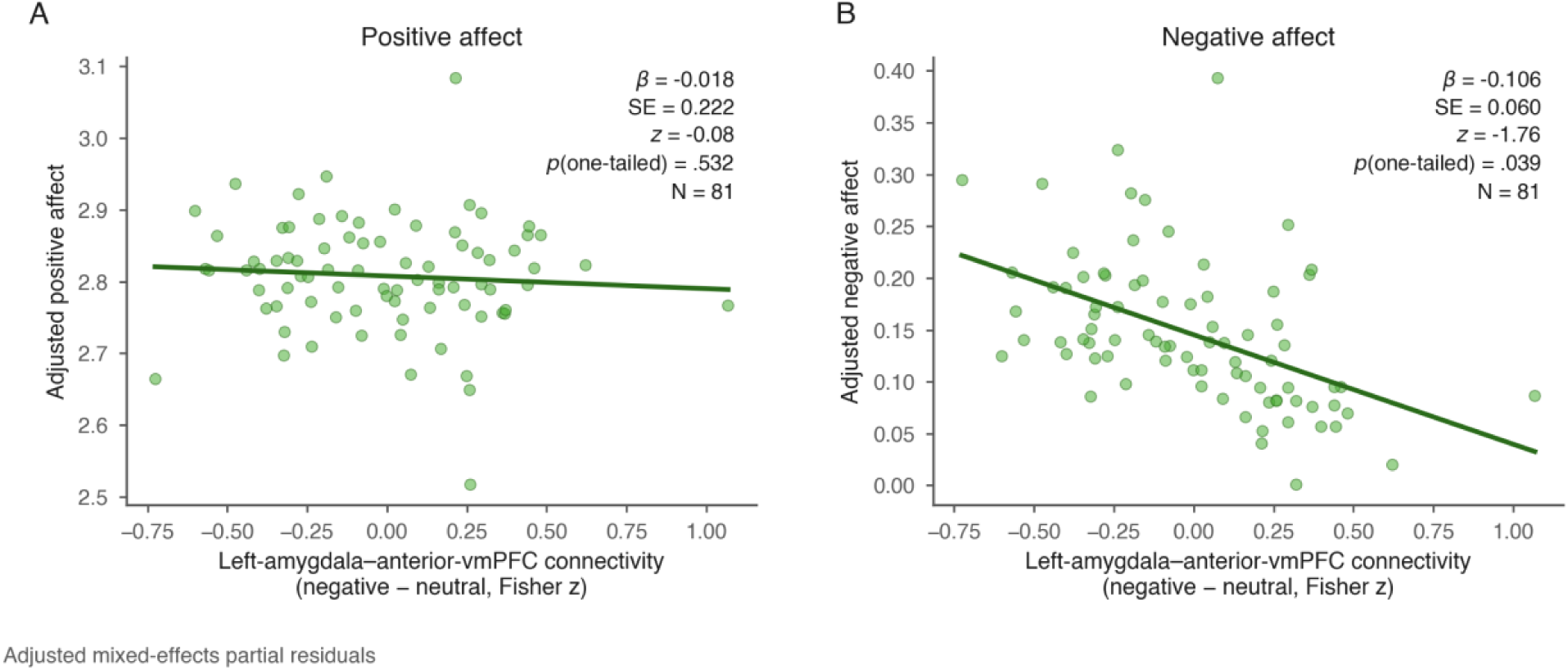
Left-amygdala–anterior vmPFC connectivity and daily affect. Mixed-effects partial residuals show adjusted associations for negative-minus-neutral connectivity with (A) positive affect and (*B*) raw negative affect in the primary diary + fMRI sample (*N* = 81). Lines show fitted fixed-effect associations.

**Table 1.**
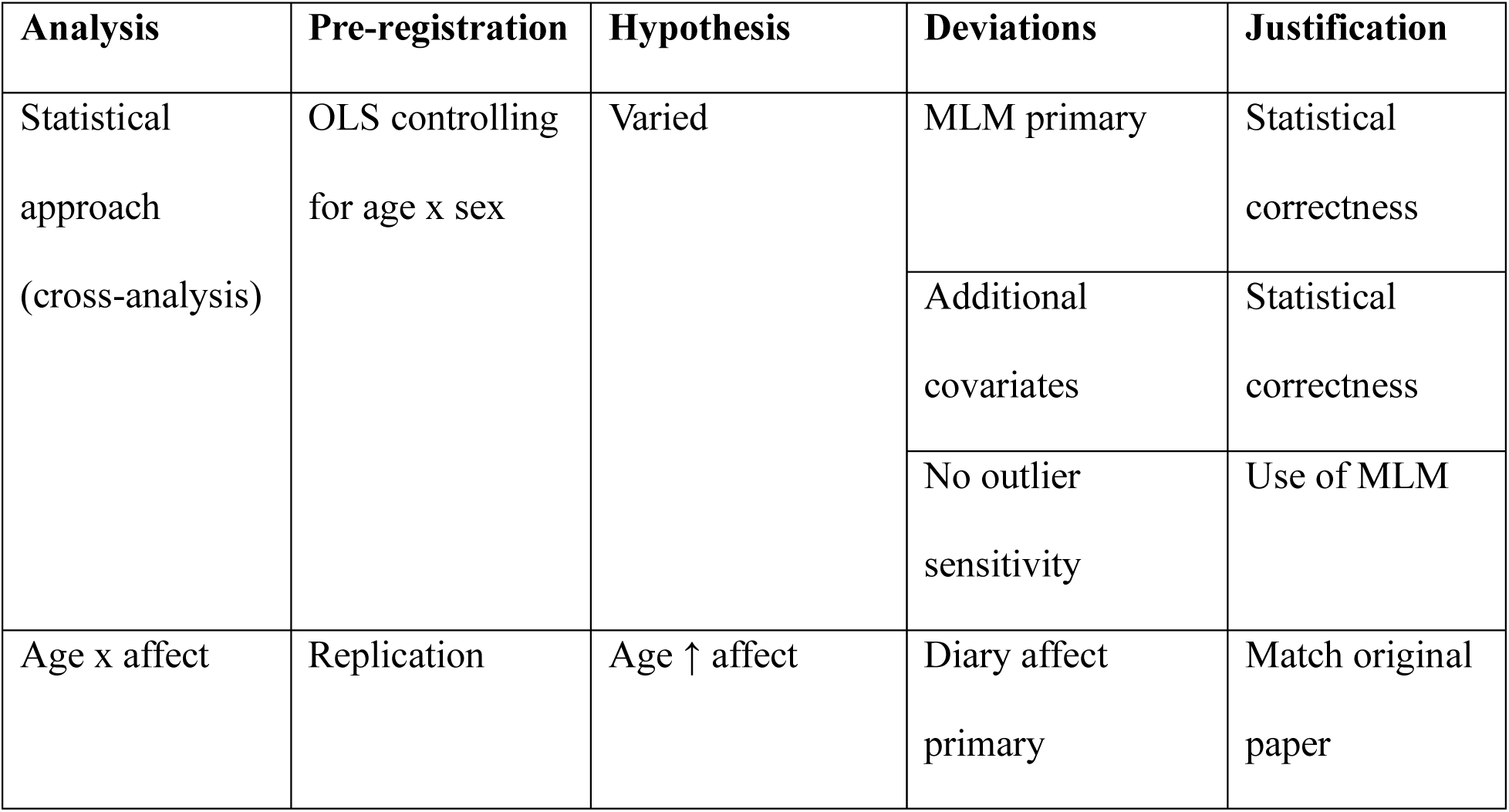
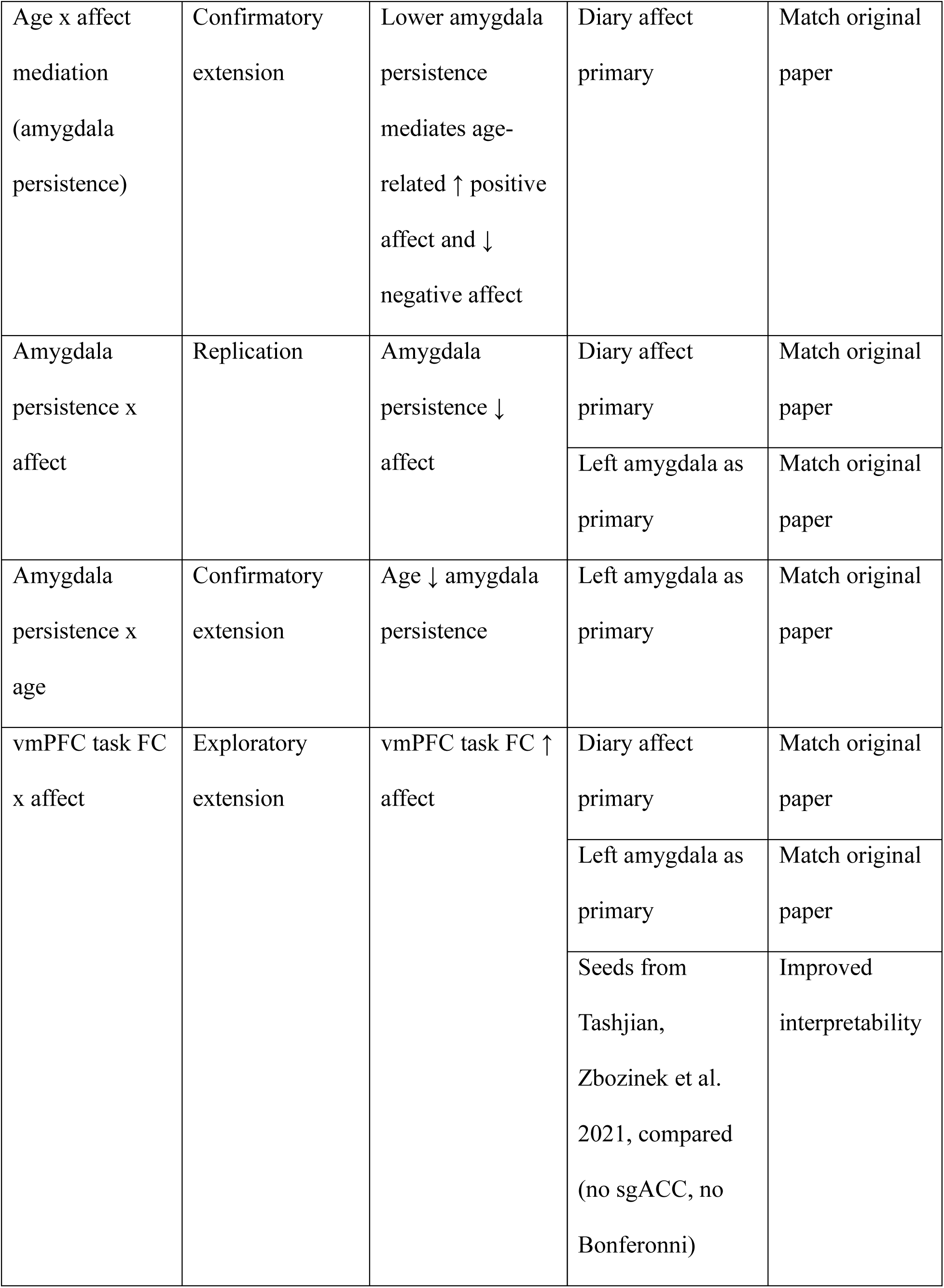
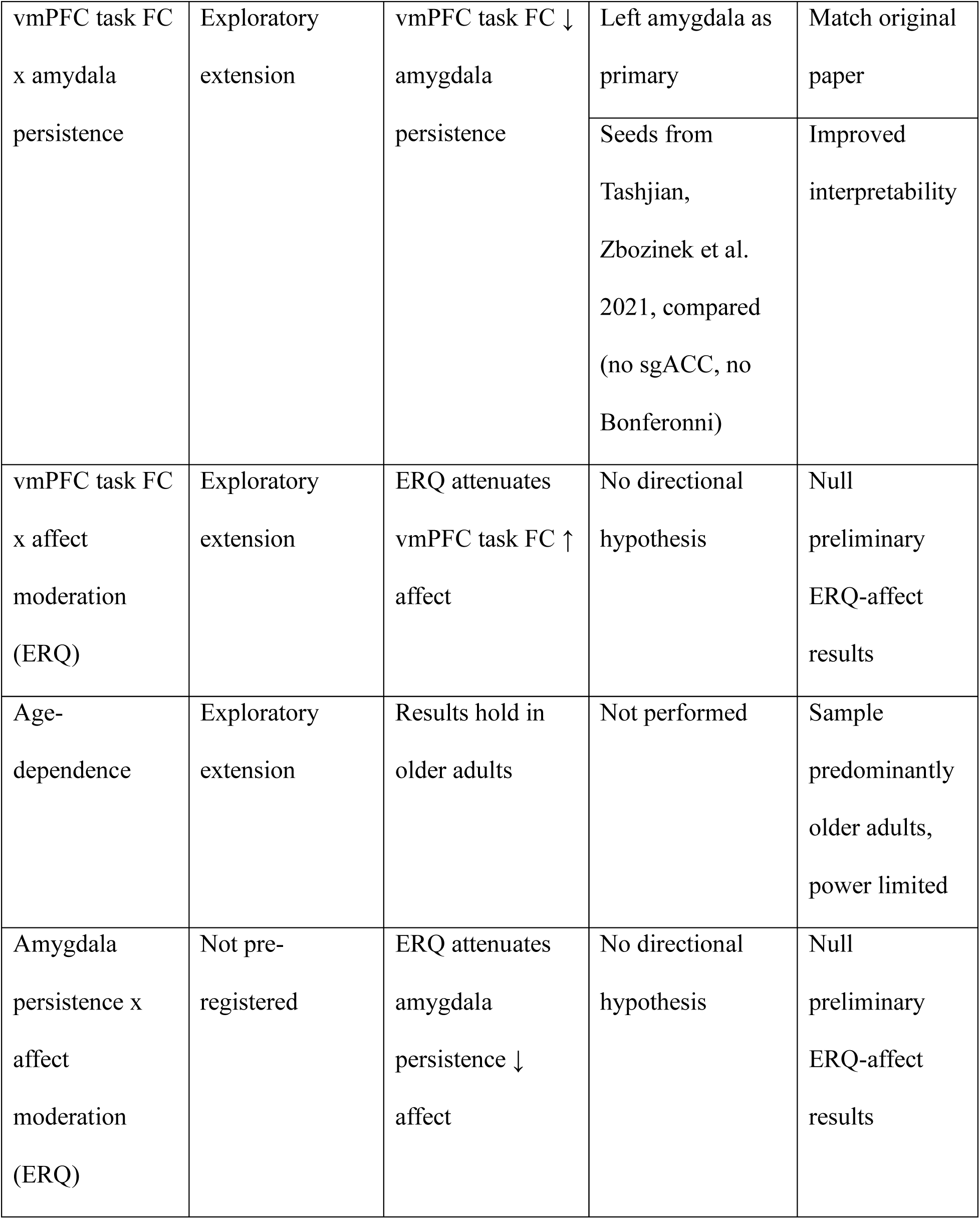
Description of pre-registered analyses, including hypotheses, deviations from pre-registration, and justifications.

## Methods

### Transparency and Openness

This study used existing MIDUS data. Sample-size determination is described in Participants. We report how we determined our sample size, all data exclusions (if any), all manipulations, and all measures in the study. The M3 cross-sectional design, hypotheses, and analytic plan were preregistered before analysis (https://doi.org/10.17605/OSF.IO/85H2D), deviations are reported in Table 1, and the subsequent MR1 evaluation followed the finalized M3 analysis plan. The authors do not redistribute participant-level MIDUS data, including derived imaging measures and participant-level quality-control summaries. Access to these data is governed by the applicable MIDUS data-use agreements. Public-use data and documentation are available through NACDA/ICPSR (https://www.icpsr.umich.edu/web/ICPSR/series/203), and restricted neuroimaging data may be requested through the MIDUS Neuroscience Repository (https://midus.wisc.edu/midus-neuroscience-repository/). Analysis code and disclosure-reviewed aggregate results are available at https://github.com/adamgeorgeturnbull/MIDUS_AMYGDALA_PERSISTENCE. Study documentation and materials supplied by MIDUS are available through the MIDUS (https://midus.wisc.edu/) and ICPSR (https://www.icpsr.umich.edu/web/ICPSR/series/203) project pages, subject to their applicable access conditions.

Imaging preprocessing used FSL 6.0.7.10 (Jenkinson et al., 2012), fMRIPrep 24.1.0 (Esteban et al., 2019), and FreeSurfer 7.4.1 (Fischl, 2012). Python analyses used Nilearn 0.12.1 (Nilearn contributors et al., 2025), NumPy 2.0.2 (Harris et al., 2020), pandas 2.3.3 (McKinney, 2010), SciPy 1.13.1 (Virtanen et al., 2020), statsmodels 0.14.6 (Seabold & Perktold, 2010), and scikit-learn 1.6.1 (Pedregosa et al., 2011). Figures used Matplotlib 3.9.4 (Hunter, 2007) and seaborn 0.13.2 (Waskom, 2021). The code repository includes a requirements file specifying the remaining Python dependencies.

Generative artificial intelligence tools, Claude (Anthropic, 2026) and Codex (OpenAI, 2026), were used through iterative prompting to assist with analysis-code development and review, visualization-code development, and drafting and editing portions of the manuscript. The tools received only code, synthetic data, aggregate statistical outputs, and manuscript text; no MIDUS participant-level data were shared. All AI-assisted outputs were reviewed, verified, and finalized by the authors, who take responsibility for the content.

### Design

This study used a cross-sectional observational design to test preregistered hypotheses replicating associations between amygdala neural persistence and daily-life affect and extending these hypotheses to vmPFC–amygdala circuitry. The MIDUS 3 (M3) hypotheses and analysis plan were preregistered on the Open Science Framework (https://doi.org/10.17605/OSF.IO/85H2D) before analysis. Deviations from the preregistered plan are summarized in Table 1. A subsequent, analysis-plan-constrained evaluation in MIDUS Refresher 1 (MR1) examined the corresponding persistence, age, and functional-connectivity associations in an independent cohort.

### Participants

Both MIDUS cohorts were assessed in the United States. Sample sizes were determined by the availability of the required diary, demographic, and neuroimaging variables and by the quality-control and complete-case criteria described below. The preregistration included a detectable-effect calculation for the anticipated neuroscience sample; it did not determine recruitment for these secondary analyses.

The M3 primary diary + fMRI sample was drawn from the M3 daily diary (2017–2019; Ryff & Almeida, 2022) and neuroscience (2017–2022; Ryff & Davidson, 2025a) projects. The daily diary sample comprised 1,174 participants (mean age = 67.6 years, *SD* = 10.3, range = 47–94; 57.3% female). Race was available for 1,168 diary participants: 82.4% White, 10.6% Black, 1.4% Native American, 0.3% Asian, and 5.3% other race. Of these, 137 also had age data from the neuroscience visit (the diary–neuroscience sample). The final fMRI sample comprised 127 participants who met all imaging quality-control criteria (mean age = 65.7 years, *SD* = 9.5, range = 48–95; 59.8% female). This sample was 75.6% White, 18.9% Black, 0.8% Native American, 1.6% Asian, and 3.1% other race. The primary diary + fMRI sample comprised 81 participants (mean age = 67.2 years, *SD* = 9.8, range = 48–95; 61.7% female; 82.7% White, 14.8% Black, 1.2% Native American, and 1.2% Asian), of whom 80 had complete emotion-regulation questionnaire data for moderation analyses. PANAS sensitivity analyses used the final fMRI sample (*N* = 127), irrespective of diary participation.

MR1 daily diary data were collected in 2012–2014 (Ryff & Almeida, 2020), and neuroscience data in 2012–2016 (Ryff & Davidson, 2025b). The MR1 imaging cohort contained 123 participants with reproduced imaging measures. Of these, 115 met the MR1 quality-control criteria (the MR1 final fMRI sample) (mean age = 48.4 years, *SD* = 11.9, range = 26–76; 53.9% female; 64.3% White, 28.7% Black, 1.7% Native American, 0.9% Asian, and 4.3% other race), and 48 had both daily diary data and imaging data meeting QC criteria (the MR1 diary + fMRI sample) (mean age = 46.9 years, *SD* = 9.5, range = 29–70; 54.2% female; 93.8% White and 6.2% other race).

MIDUS neuroscience procedures were approved by the University of Wisconsin–Madison Institutional Review Board, and daily diary procedures were approved by the Pennsylvania State University Institutional Review Board. Participants provided informed consent for participation in MIDUS.

### Behavioral Preprocessing

#### Daily diary affect

Daily affect was assessed for approximately eight consecutive days. The 27 affect items were rated on a five-point scale from 0 (none of the time) to 4 (all of the time). Positive affect (PA) was calculated as the mean of 13 positive-mood items, and negative affect (NA) as the mean of 14 negative-mood items. Days with missing responses on any affect item were excluded, and participant-level PA and NA scores were calculated by averaging across complete diary days. Because NA was positively skewed, a secondary log-transformed score was calculated as log(NA + c), where c was half the minimum nonzero NA value. Raw NA was the primary negative-affect outcome; log-transformed NA was treated as a robustness outcome. Corresponding MR1 diary items were scored using the same procedure. Repeated MR1 participant-day records were counted once after verifying agreement on all affect items and diary dates, so each complete day contributed equally.

### PANAS

Positive and negative affect assessed with the PANAS (Watson et al., 1988) at the neuroscience visit were examined as convergent-validity sensitivity outcomes. PANAS negative affect was natural-log transformed without an added constant because all valid scores were positive.

#### Emotion regulation

Reappraisal and suppression were assessed with the Emotion Regulation Questionnaire (Gross & John, 2003) at the neuroscience visit.

#### Demographic variables

Age at the diary wave was used for diary-only age analyses, whereas age at the neuroscience visit was used in analyses involving neuroscience measures. Age at diary assessment was estimated from diary year and birth year for the main MIDUS cohort, and from interview age and elapsed time for the Milwaukee cohort. Biological sex retained the original MIDUS coding (1 = male, 2 = female). Race was represented by indicator variables with White as the reference category; indicators with no variation within the model sample were omitted. The interval between the diary and neuroscience visits was calculated in months. In M3, twin pairs were identified using twin-sample membership and a shared family identifier. Participants without an observed co-twin were assigned individual grouping identifiers. The available MR1 analytic inputs contained no usable family, household, sibling, or twin-pair grouping identifier.

#### fMRI Task and Acquisition

Participants completed three runs of an emotional image-viewing task. Each run included 10 negative, 10 neutral, and 10 positive images, followed by neutral face probes. Images were presented for 4 s, followed by a 2-s fixation interval and a 0.5-s face probe. Participants categorized the gender of each face. Negative and positive images were matched on normative arousal, and images within each valence category were matched on luminosity, visual complexity, and social content.

M3 data were acquired on a 3T GE MR750 scanner using a 32-channel head coil. Each of the three functional runs contained 231 volumes (TR = 2,000 ms; TE = 20 ms; flip angle = 60°; field of view = 220 mm; matrix = 96 × 64; 40 interleaved sagittal slices, 3-mm thickness with a 1-mm gap). A T1-weighted BRAVO anatomical image was also acquired (TR = 8.2 ms; TE = 3.2 ms; flip angle = 12°; field of view = 256 mm; matrix = 256 × 256; 160 axial slices; inversion time = 450 ms).

#### fMRI Preprocessing and Quality Control

First, because the M3 raw images had a nonstandard acquisition direction, images were reoriented and the axes temporarily exchanged before slice-timing correction. Slice-timing correction was performed in FSL 6.0.7.10 using the 1-based interleaved acquisition order derived from the imaging metadata; the axes and original geometry were then restored. Structural and functional data were then processed with fMRIPrep 24.1.0, with fMRIPrep slice-timing correction disabled so that slice-timing correction was applied exactly once. Preprocessing included anatomical reconstruction, motion correction, functional-to-anatomical registration, and normalization to MNI152NLin2009cAsym space. Framewise displacement (FD) was derived from the resulting fMRIPrep confound time series.

The M3 final fMRI sample required all three functional runs to pass visual quality control, mean FD across task runs to be less than 0.5 mm, and all six directional cross-run pairs needed for the primary left-amygdala persistence measure to be available. Visual ratings were evaluated separately from quantitative FD. MR1 imaging data were reproduced using fMRIPrep 24.1.0 with standard BIDS slice-timing handling due to standard acquisition direction. Visual quality-control was similarly performed and FD was recalculated from the reproduced confound time series. In both final fMRI samples, all participants had three task runs containing 231 volumes each before removal of the four initial dummy volumes.

## fMRI Analysis

### First-level general linear model

A first-level general linear model was estimated separately for each run using nilearn (Nilearn contributors et al., 2025). Six task regressors modeled negative, neutral, and positive images and the neutral face probes that followed images in each of those three valence conditions. Thus, the face regressors indexed the valence of the preceding image rather than the valence of the face itself. Each regressor was convolved with the Glover hemodynamic response function (Glover, 1999). Models included a 24-parameter motion model, cosine drift terms with a 1/128-Hz high-pass cutoff, and an AR(1) noise model. The first four volumes and corresponding confound rows were discarded.

### Amygdala neural persistence

Left and right amygdala regions were defined from the Harvard– Oxford atlas (FMRIB Software Library, n.d.) at a 50% probability threshold. For each hemisphere, persistence was calculated as the spatial Pearson correlation between the voxelwise beta pattern for negative images in run i and the pattern for neutral faces following negative images in run j, where i ≠ j. Correlations were transformed to Fisher *z* values and averaged across the six directional cross-run pairs. Left-amygdala negative persistence was the primary measure; right-amygdala, positive-image, and vmPFC persistence were examined in sensitivity analyses. Anterior and posterior vmPFC persistence used the same six directional image-to-following-face cross-run correlations within the 10-mm spherical regions defined below.

### Task-based functional connectivity

Amygdala–vmPFC functional connectivity was estimated with a least-squares-separate beta-series procedure (Mumford et al., 2012; Rissman et al., 2004). For each image trial, the trial of interest was modeled separately, other trials of the same valence were combined in a second regressor, and trials from each remaining condition were combined into condition-specific regressors. Each image event was modeled for 6s, encompassing the 4s image and subsequent 2s fixation interval. Face-probe events were not modeled with separate regressors. Trial-specific beta estimates were extracted from left and right amygdala and from 10-mm-radius anterior (MNI coordinates: −2, 46, −10) and posterior (0, 26, −12) vmPFC spheres. Beta-series correlations were calculated separately for each connection, condition, and run, transformed to Fisher *z*, and averaged across runs. Negative-minus-neutral connectivity was the primary contrast. Analyses focused on left-amygdala connections, with right-amygdala, positive-minus-neutral, and negative-condition-only measures examined as sensitivity analyses.

#### Statistical Analyses

For M3, linear mixed-effects models with a random intercept for family were the primary adjusted analyses. Paired twins shared a family grouping identifier; participants without an observed co-twin were assigned individual identifiers. Models were estimated with restricted maximum likelihood. Supplementary analyses included zero-order Pearson correlations and ordinary least-squares (OLS) regression; OLS models included twin-pair indicators instead of the family random intercept. Supplementary Tables S5 and S7 report M3 OLS and zero-order correlation comparisons for main and sensitivity analyses, respectively; Tables S6 and S8 provide the corresponding MR1 comparisons.

Adjusted models included age at the neuroscience visit, biological sex, and race dummy variables. Models with diary outcomes additionally included the interval between the diary and neuroscience visits and the number of complete diary days. Age was omitted from the covariate set when it was the predictor. Prespecified directional hypotheses were evaluated with one-tailed tests, calculated from the corresponding two-tailed probability according to the observed direction. All reported regression coefficients (β) are unstandardized. Confidence intervals are two-sided 95% intervals: normal-Wald intervals for mixed-effects coefficients, Student-*t* intervals for OLS coefficients, condition means, and paired mean differences, and Fisher-*z* intervals for Pearson correlations. A significant prespecified one-tailed test can therefore accompany a two-sided interval that includes zero. Supplementary Tables S1–S11 provide effect estimates and intervals for analyses summarized in the text. Raw NA was the primary negative-affect outcome, with log-transformed NA used to assess robustness in the main and diary-affect sensitivity analyses in both cohorts. Right-hemisphere and other specificity analyses used two-tailed tests unless an explicit directional prediction had been specified. Convergence correlations, task-condition tests, and exploratory moderation tests were two-tailed. Predictor and moderator variables were mean centered before interaction terms were calculated. Sensitivity and exploratory analyses were reported with nominal *p* values without multiplicity adjustment and were not used to redefine the primary hypotheses.

The prespecified statistical indirect association between age and daily affect through left-amygdala negative persistence was evaluated in the primary diary + fMRI sample. Separate mixed-effects models estimated the age–persistence path (a), the persistence–affect path adjusted for age (*b*), the direct age–affect path (c′), and the total age–affect path (c) using a common complete-case sample and the covariates described above. Tests of mediation path coefficients were one-tailed in the hypothesized directions. Indirect effects (a × *b*) were evaluated in the predicted direction (positive for PA and negative for NA) using one-sided 95% percentile bounds from 2,000 family-level cluster-bootstrap resamples; two-sided 95% confidence intervals were retained for estimation. Age at the diary assessment and log-transformed negative affect were examined as sensitivity analyses. Because assessment order varied across participants and the study did not establish the temporal sequence required for causal mediation, these models were interpreted as statistical indirect associations rather than causal mediation.

For MR1, corresponding participant-level OLS models were used because the available analytic inputs contained no usable family, household, sibling, or twin-pair grouping identifier. Models included the same available covariates and directional tests as the corresponding M3 analyses. The analysis of the MR1 final fMRI sample was primary, and models without QC restriction were retained as secondary analyses. Direct results were classified as supported when significant in the predicted direction, directionally consistent when nonsignificant but in the predicted direction, and not supported when the estimate was in the opposite direction. MR1 sensitivity analyses were interpreted as robustness or specificity checks and did not alter the direct-replication classification. MR1 sensitivity results are reported in the Supplementary Materials.

## Results

### Convergence of Affect Measures

Daily diary and PANAS affect scores were positively correlated in the diary–PANAS sample of 136 participants with both measures: diary PA with PANAS positive affect (*r*(134) = .52, 95% CI [.39, .63]), diary NA with PANAS negative affect (*r*(134) = .42, 95% CI [.28, .55]), and log-transformed diary NA with log-transformed PANAS negative affect (*r*(134) = .39, 95% CI [.23, .52]; all *p*s < .001). Adjusted mixed-effects associations were also positive (all *p*s < .001; Supplementary Table S9). These correlations support convergence while showing substantial unshared variance between the measures.

### Task-Related Activation and Functional Connectivity

In the M3 final fMRI sample, left-amygdala activation was greater than zero for negative images (*M* = 0.0492, 95% CI [0.0319, 0.0665], *t*(126) = 5.63, *p* < .001) and positive images (*M* = 0.0401, 95% CI [0.0242, 0.0560], *t*(126) = 4.99, *p* < .001), but not for neutral images (*M* = 0.0148, 95% CI [−0.0013, 0.0308], *t*(126) = 1.82, *p* = .070). Right-amygdala activation was positive in all three conditions (all *t*s ≥ 5.30, all *p*s < .001). Both amygdalae responded more strongly to negative than neutral images (left: mean difference = 0.0344, 95% CI [0.0164, 0.0524], *t*(126) = 3.77, *p* < .001; right: mean difference = 0.0252, 95% CI [0.0095, 0.0410], *t*(126) = 3.17, *p* = .002), whereas negative and positive responses did not differ significantly (both *p*s ≥ .074).

Anterior vmPFC showed significant deactivation during negative images (*M* = −0.0397, 95% CI [−0.0618, −0.0175], *t*(126) = −3.54, *p* < .001) and neutral images (*M* = −0.0289, 95% CI [−0.0480, −0.0098], *t*(126) = −2.99, *p* = .003), but not positive images (*M* = −0.0124, 95% CI [−0.0316, 0.0068], *t*(126) = −1.28, *p* = .204). Deactivation was greater during negative than positive images (negative-minus-positive mean difference = −0.0273, 95% CI [−0.0454, −0.0092], *t*(126) = −2.98, *p* = .003). Posterior vmPFC activation did not differ from zero or between conditions (smallest *p* = .114).

All 12 amygdala–vmPFC connection-by-condition estimates were positive (all *p*s ≤ .003), indicating reliable task-related coupling. However, no connection differed significantly between negative and neutral or between negative and positive conditions (all *p*s ≥ .097; condition means, mean differences, and their 95% CIs appear in Supplementary Table S10). Anterior and posterior vmPFC estimates showed moderate-to-strong correspondence across conditions for activation (*r*(125) = .62–.73), persistence (*r*(125) = .47–.48), and beta-series connectivity (*r*(125) = .52–.64; all *p*s < .001), while retaining substantial unshared variance.

### Age and Affect

In the daily diary sample, older age was associated with higher PA (β = 0.00987, 95% CI [0.00593, 0.01381], *z* = 4.91, *p* < .001), lower raw NA (β = −0.00285, 95% CI [−0.00422, −0.00147], *z* = −4.07, *p* < .001), and lower log-transformed NA (β = −0.0222, 95% CI [−0.0304, −0.0139], *z* = −5.28, *p* < .001) in the primary adjusted models (*N* = 1,168 after covariate-complete-case restriction). In the diary–neuroscience sample (*N* = 137), age was not significantly associated with PA (β = 0.00829, 95% CI [−0.00495, 0.02152], *z* = 1.23, *p* = .110) or raw NA (β = −0.00215, 95% CI [−0.00728, 0.00298], *z* = −0.82, *p* = .205), but was associated with lower log-transformed NA (β = −0.0319, 95% CI [−0.0601, −0.0038], *z* = −2.22, *p* = .013). In the PANAS sample (*N* = 230), age was not significantly associated with positive affect (β = 0.00117, 95% CI [−0.00854, 0.01088], *z* = 0.24, *p* = .406) or raw negative affect (β = −0.00503, 95% CI [−0.01116, 0.00110], *z* = −1.61, *p* = .054), but was associated with lower log-transformed negative affect (β = −0.00395, 95% CI [−0.00769, −0.00022], *z* = −2.08, *p* = .019).

### Amygdala Persistence and Affect

In the primary diary + fMRI sample (*N* = 81), greater left-amygdala negative persistence was associated with higher raw NA in the primary mixed-effects model (β = 0.278, 95% CI [0.012, 0.544], *SE* = 0.136, *z* = 2.05, *p* = .020). Associations with PA (β = −0.700, 95% CI [−1.633, 0.233], *SE* = 0.476, *z* = −1.47, *p* = .071) and log-transformed NA (β = 1.95, 95% CI [−0.55, 4.45], *SE* = 1.27, *z* = 1.53, *p* = .063) were in the predicted directions but were not significant. The original MIDUS II analytic roster was unavailable. We therefore used an overlap proxy, excluding participants with recorded MIDUS II diary entries and indicated neuroimaging completion. In the 55-person overlap-proxy-excluded subsample, the raw NA association was directionally consistent with the primary result and somewhat smaller in magnitude (β = 0.223 versus 0.278; subsample 95% CI [−.043, .489], *p* = .0504). This provides tentative directional support, but the estimate remains imprecise and does not meet the *p* < .05 criterion. Associations with PA and log-transformed NA were also in the predicted directions but nonsignificant (*p*s = .112 and .084, respectively).

In the MR1 diary + fMRI sample (*N* = 48), the raw NA estimate was in the same direction and numerically similar in magnitude to M3, but was not significant (β = 0.311 versus 0.278; MR1 95% CI [−0.129, 0.752], *SE* = 0.218, *t*(41) = 1.43, *p* = .081). The PA and log-transformed NA estimates were also in the same directions as M3 but smaller in magnitude and nonsignificant (PA: β = −0.195 versus −0.700; MR1 95% CI [−1.367, 0.976], *SE* = 0.580, *t*(41) = −0.34, *p* = .369; log-transformed NA: β = 1.42 versus 1.95; MR1 95% CI [−1.11, 3.95], *t*(41) = 1.14, *p* = .131). These descriptive comparisons suggest directional consistency, but the MR1 estimates remain imprecise and do not establish replication.

Right-amygdala negative persistence did not significantly predict diary PA, raw NA, or log-transformed NA in the primary M3 sensitivity models (*p*s ≥ .084). Left-amygdala persistence also did not significantly predict PANAS affect in the final fMRI sample (*p*s ≥ .088). Positive persistence was not significantly associated with diary affect (all ps > .222). Regional activation measures were also not significantly associated with diary affect (all *p*s > .083). Anterior and posterior vmPFC persistence were also not significantly associated with diary affect (*p*s ≥ .121; Supplementary Table S3).

### Age and Amygdala Persistence

Older age was associated with lower left-amygdala negative persistence in the primary mixed-effects model (β = −0.00260, 95% CI [−0.00504, −0.00016], *SE* = 0.00124, *z* = −2.09, *p* = .018; *N* = 127). Associations between age and right-amygdala negative persistence or left/right positive persistence were not significant in sensitivity analyses (all *p*s ≥ .159). Age was also not significantly associated with anterior or posterior vmPFC persistence (*p*s = .510 and .902, respectively; Supplementary Table S3).

In MR1, age was positively rather than negatively associated with left-amygdala persistence in the MR1 final fMRI sample (*N* = 115; β = 0.00134, 95% CI [−0.00157, 0.00425], *SE* = 0.00147, *t*(108) = 0.91, *p* = .818). This estimate was nonsignificant and opposite to the association observed in M3, although the age of this sample was younger than in M3.

### Persistence mediation of age and affect

We next tested the prespecified statistical indirect association between age and daily affect through left-amygdala negative persistence. Although age was associated with lower persistence in the final fMRI sample, the age–persistence path was weaker in the primary diary + fMRI sample, β = −0.00132, 95% CI [−0.00441, 0.00176], *SE* = 0.00157, *z* = −0.84, *p* = .200, *N* = 81. There was no evidence of an indirect association with positive affect, ab = 0.000927, 95% bootstrap CI [−0.001341, 0.005164], negative affect, ab = −0.000368, 95% bootstrap CI [−0.001092, 0.000366], or log-transformed negative affect, ab = −0.00258, 95% bootstrap CI [−0.01041, 0.00237]. Within the primary diary + fMRI sample (*N* = 81), older age was associated with lower raw and log-transformed NA before and after adjustment for persistence. Adjusted associations were β = −0.00434, 95% CI [−0.00807, −0.00062], *z* = −2.29, *p* = .011 for raw NA and β = −0.0576, 95% CI [−0.0906, −0.0246], *z* = −3.42, *p* < .001 for log-transformed NA. The conclusion was unchanged when age at the diary assessment was used instead: the age– persistence path was β = −0.00115, 95% CI [−0.00423, 0.00194], *SE* = 0.00157, *z* = −0.73, *p* = .233, and none of the indirect associations met the one-tailed significance criterion.

### Amygdala–vmPFC Functional Connectivity and Affect

In the primary diary + fMRI sample (*N* = 81), greater left-amygdala–anterior vmPFC connectivity for negative relative to neutral images was associated with lower raw NA (β = −0.106, 95% CI [−0.223, 0.012], *SE* = 0.0601, *z* = −1.76, *p* = .039) and lower log-transformed NA (β = −1.06, 95% CI [−2.01, −0.11], *SE* = 0.485, *z* = −2.19, *p* = .014) in the primary mixed-effects models. It was not associated with PA (β = −0.0178, 95% CI [−0.4527, 0.4172], *z* = −0.08, *p* = .532).

Posterior vmPFC connectivity did not significantly predict any outcome in the prespecified direction. Its mixed-effects association with PA was negative (β = −0.353, 95% CI [−0.649, −0.057], *z* = −2.33, *p* = .990). Posterior connectivity was not significantly associated with raw NA (β = −0.0590, 95% CI [−0.1676, 0.0495], *z* = −1.07, *p* = .143) or log-transformed NA (β = −0.562, 95% CI [−1.546, 0.422], *z* = −1.12, *p* = .132).

In MR1, left-amygdala–anterior vmPFC connectivity was negatively but non-significantly associated with raw NA (*N* = 48; β = −0.0801, 95% CI [−0.2832, 0.1229], *SE* = 0.101, *t*(41) = −0.80, *p* = .215). The MR1 point estimate was in the same direction but smaller in magnitude than M3 (β = −0.0801 versus −0.106); its confidence interval was wider and included zero. The log-transformed NA estimate was also in the same direction but smaller in magnitude than M3 and nonsignificant (β = −0.340 versus −1.06; MR1 95% CI [−1.500, 0.820], *SE* = 0.574, *t*(41) = −0.59, *p* = .279).

In PANAS sensitivity analyses (*N* = 127), both anterior and posterior left-amygdala–vmPFC connectivity were associated with lower PANAS negative affect in mixed-effects models (raw: anterior β = −0.251, 95% CI [−0.467, −0.035], and posterior β = −0.249, 95% CI [−0.463, −0.036], both *p*s = .011; log transformed: anterior β = −0.139, 95% CI [−0.264, −0.014], *p* = .015, and posterior β = −0.128, 95% CI [−0.253, −0.003], *p* = .023). Neither connection predicted PANAS positive affect (*p*s ≥ .329). Right-amygdala and positive-minus-neutral connectivity did not significantly predict diary PA, raw NA, or log-transformed NA (*p*s ≥ .205). Negative-condition-only connectivity was also not significantly associated with diary affect (all *p*s > .074). In an exploratory analysis of relative anterior versus posterior connectivity, the left anterior-minus-posterior index predicted higher PA in the mixed-effects model (β = 0.471, 95% CI [0.103, 0.839], *z* = 2.51, *p* = .006). The right-amygdala anterior-minus-posterior index was not significantly associated with diary affect (all *p*s > .057), and neither hemisphere’s index was significantly associated with PANAS affect (all *p*s > .422).

### Functional Connectivity and Amygdala Persistence

Neither left anterior nor left posterior amygdala–vmPFC connectivity for negative relative to neutral images predicted left-amygdala negative persistence in the primary mixed-effects models (anterior: β = −0.0180, 95% CI [−0.0849, 0.0489], *z* = −0.53, *p* = .299; posterior: β = 0.0109, 95% CI [−0.0558, 0.0777], *z* = 0.32, *p* = .626; *N* = 127). These findings provided no evidence that individual differences in the connectivity contrast were associated with amygdala persistence.

### Assessment of Head Motion

Neither mean FD across runs nor mean FD during negative-condition trials was associated with left-amygdala persistence or left-amygdala–vmPFC connectivity in mixed-effects models (all *p*s ≥ .129). In motion-adjusted follow-ups, all significant primary mixed-effects findings remained significant after controlling separately for each motion measure: persistence with raw NA (*p*s = .038 and .026), age with persistence (*p*s = .019 and .019), and anterior connectivity with raw NA (*p*s = .048 and .040) and log-transformed NA (*p*s = .018 and .015). Motion-adjusted estimates and 95% CIs are reported in Supplementary Table S3. These results reduce concern that those tested associations were attributable to head motion.

### Emotion Regulation and Affect

Reappraisal and suppression were not significantly correlated, *r*(227) = −.03, *p* = .638. Neither strategy significantly predicted diary PA, raw NA, or log-transformed NA in the primary adjusted models (all *p*s ≥ .065). The reappraisal estimates for raw and log-transformed diary NA were positive, opposite to the expected association with lower negative affect, although neither was significant. In contrast, reappraisal predicted higher PANAS positive affect (β = 0.178, 95% CI [0.094, 0.262], *z* = 4.16, *p* < .001), whereas suppression predicted lower PANAS positive affect (β = −0.102, 95% CI [−0.177, −0.026], *z* = −2.63, *p* = .009) and higher PANAS negative affect (raw: β = 0.0593, 95% CI [0.0129, 0.1056], *z* = 2.51, *p* = .012; log transformed: β = 0.0360, 95% CI [0.0075, 0.0646], *z* = 2.47, *p* = .013). Age was not significantly associated with either strategy (both *p*s ≥ .204).

### Exploratory Moderation by Emotion Regulation

All moderation analyses were exploratory and used nominal tests. In the primary mixed-effects models (*N* = 80), reappraisal moderated the association between left-amygdala negative persistence and PA (βinteraction = 1.00, 95% CI [0.03, 1.97], *SE* = 0.496, *p* = .044), and suppression moderated the same persistence–PA association (βinteraction = 0.586, 95% CI [0.290, 0.883], *SE* = 0.151, *p* < .001). The negative persistence–PA association weakened at higher strategy scores (Supplementary Figure S1). This was directionally consistent with attenuation, although higher suppression was associated with lower PA at average persistence. No persistence interaction significantly predicted either raw or log-transformed diary NA (all *p*s ≥ .120). Neither persistence–PA interaction was significant when PANAS positive affect was the outcome (reappraisal: *p* = .992; suppression: *p* = .321). Interpretation of the diary interactions is complicated by the nonsignificant associations of persistence, reappraisal, and suppression with diary PA in models without interaction terms.

For connectivity, suppression moderated the association between left-amygdala–posterior vmPFC connectivity and raw diary NA in the primary mixed-effects model (βinteraction = 0.0975, 95% CI [0.0012, 0.1938], *SE* = 0.0491, *p* = .047). No other diary FC interaction was significant (all *p*s ≥ .151). In the PANAS sensitivity analysis, suppression moderated the association between posterior connectivity and negative affect in the mixed-effects models (raw: βinteraction = −0.246, 95% CI [−0.442, −0.049], *p* = .014; log transformed: βinteraction = −0.128, 95% CI [−0.243, −0.013], *p* = .029). The diary connectivity–NA association also weakened at higher suppression (Supplementary Figure S1), whereas the PANAS interactions had the opposite sign. For all three primary diary interactions, the corresponding brain–affect associations in models without interactions were nonsignificant. These exploratory findings are model-dependent, do not establish beneficial compensation, and require replication given the multiple unadjusted tests.

## Discussion

The present study tested a set of pre-registered hypotheses linking amygdala persistence, amygdala–vmPFC task-based connectivity, aging, and daily-life affect in MIDUS 3, with an analysis-plan-constrained validation in MIDUS Refresher 1. Older age was associated with higher positive and lower negative affect in the daily diary sample (all *p*s < .001), although these associations were attenuated in the diary–neuroscience and PANAS samples. We partially replicated the original persistence–affect finding (Puccetti et al., 2021): greater left-amygdala negative persistence was associated with higher raw negative affect (*p* = .020), whereas its associations with positive affect and log-transformed negative affect were in the predicted directions but did not reach significance (*p*s ≥ .063). Older age was also associated with lower left-amygdala negative persistence (*p* = .018) in the final fMRI sample. However, the age– persistence association was attenuated in the primary diary + fMRI sample (*p* = .200), and none of the indirect age–affect associations through persistence met the one-tailed significance criterion. Greater left-amygdala–anterior vmPFC connectivity during negative relative to neutral image viewing was associated with lower raw and log-transformed negative affect (*p*s = .014– .039), but neither anterior nor posterior connectivity predicted persistence (*p*s ≥ .299), showing associations of both persistence and connectivity with affect in separate models. In the smaller independent MIDUS Refresher 1 cohort, the corresponding affect estimates were directionally similar but did not reach significance (*p*s ≥ .081), whereas the age–persistence estimate reversed direction. Together, these findings provide partial support for the pre-registered persistence and connectivity hypotheses while underscoring the need for replication in larger independent samples and further emphasizing the fact that fMRI-behavior associations often have small and unreliable effect sizes.

The association between older age and lower left-amygdala persistence extends the literature on age-related differences in emotional experience (Mather, 2012) by identifying a feature of amygdala response patterns, rather than mean regional activation, that varies with age. It is possible that less persistent amygdala representations of negative information contribute to the tendency for older adults to experience less negative affect, although cross-sectional mediation analyses in this manuscript were non-significant (Mather, 2012). The age–persistence association was observed in the final fMRI sample but was weaker in the primary diary + fMRI sample, the statistical indirect effects included zero, and the order of diary and neuroimaging assessments varied across participants. The association reversed direction in the younger MIDUS Refresher 1 cohort, however, this could suggest this difference is specific to the age range in M3. Lifespan and longitudinal studies are needed to establish whether amygdala persistence changes within individuals with age and whether such change prospectively relates to affect.

These findings have implications for our understanding of the neural circuit-level mechanisms of affect in older adults. First, greater amygdala persistence was associated with higher raw daily negative affect (*p* = .020). This is important given the lack of reliable brain-behavior relationships between amygdala activation (Sicorello et al., 2025) and affect in the literature and further suggests that analyses looking at patterns of activation in amygdala provide information missed by traditional activation analyses. Contrary to our hypotheses, vmPFC–amygdala beta-series connectivity for negative relative to neutral images did not predict amygdala persistence (*p*s ≥ .299), although anterior connectivity was associated with lower daily negative affect (*p*s = .014–.039). This provides preliminary evidence that this circuit, might be relevant to daily negative affect, although its relationship to amygdala persistence remains unclear.

This may have implications for older adults at-risk for Alzheimer’s Disease (AD): a recent perspective suggests that vmPFC-amygdala circuitry may be vulnerable to early AD pathology (Turnbull et al., 2026) and that model-based emotion regulation strategies may be able to compensate for these changes. The present study found associations between vmPFC-amygdala task connectivity and daily negative affect (*p*s = .014–.039), but did not provide evidence that reappraisal moderated these associations (*p*s ≥ .151). Reappraisal did attenuate the non-significant persistence daily positive affect relationship but should be interpreted cautiously given the exploratory nature of these analyses and the lack of significant main effects. Future studies should test these relationships in samples including AD biomarkers to assess whether there is any impact of amyloid or ptau in the vmPFC. Additionally, although our exploratory analysis and primary findings provided some evidence that the anterior vmPFC (hypothesized to be more related to “safety” signaling) may be preferentially related to affect, this was not consistent across analyses and methods, and there were relatively strong correlations between anterior and posterior vmPFC-amygdala connectivity. Overall, this suggests that the hypothesized differential roles for anterior and posterior vmPFC in safety and threat signaling (Tashjian et al., 2021), respectively, warrants further investigation.

Our results also have important methodological considerations for affective science more broadly. Although trait affect and daily life affect showed reasonable associations with each other, they showed very different relationships to both neuroscience and behavioral measures. In particular, daily life affect was associated with amygdala persistence, while trait affect was not. These differences were also apparent for amygdala-vmPFC task connectivity, but less so, with relationships being more consistent across measures. For emotion regulation, only trait affect measured via a questionnaire showed significant relationships, potentially highlighting the consistency between measurement approaches (Podsakoff et al., 2003). Overall, these findings add to the literature that suggest that ecological and questionnaire measures of affect may be capturing distinct processes (Kahneman & Deaton, 2010), with intensive sampling potentially providing greater sensitivity for identifying links to biological processes (Conner & Barrett, 2012). Additionally, findings were sometimes but not always consistent across negative and positive affect, which is also in line with the literature showing some shared and some distinct mechanisms (Lindquist et al., 2012; Zhang et al., 2025).

This study has several limitations. Firstly, participants may overlap with those in the original paper, although the assessments occurred roughly 10 years later. The original analytic roster was unavailable, and excluding potential overlap using diary participation and neuroimaging completion records yielded only tentative directional support in the overlap-proxy-excluded subsample. Second, there were several deviations from pre-registration (**Table 1**). Several of these reflected well-justified methodological choices from the original paper that were overlooked by the authors, highlighting the need for care when pre-registering methodological details. Other choices reflected issues that arose when examining the data, either in data cleaning, preprocessing, or preliminary analyses. This is a well-known issue with pre-registration, where there is a need to balance pre-registering early in the process prior to looking at the data (to reduce opportunities for researcher degrees of freedom) with making the correct choices for statistical correctness. This is particularly challenging for neuroimaging analyses, where there are increased preprocessing and analysis choices (Carp, 2012), and sample size can be significantly reduced due to quality control procedures. We aimed to make choices that best balanced pre-registration and statistical correctness and we hope that researchers aiming to perform pre-registered neuroimaging analyses can learn from our approach, potentially generating multi-stage registrations that account for the need to understand the data before pre-registering hypotheses and analysis/preprocessing procedures. However, these deviations should still be a consideration when interpreting these findings. Finally, the task used here were not best suited for examination of emotion regulation: connectivity did not differ significantly between the negative and other conditions (all *p*s ≥ .097). Other tasks that are designed directly to probe emotion regulation, particularly more implicit/model-free aspects (e.g., fear conditioning and the emotional Stroop), along with tasks aimed at examining model-based strategy engagement and success, are needed to better understand these mechanisms.

### Constraints on Generality

These findings are most directly applicable to U.S. midlife and older adults who remained engaged in MIDUS and completed intensive daily-diary and neuroscience protocols. Participation in these projects and imaging quality control produced selectively retained samples. The M3 primary diary + fMRI sample was small (*N* = 81), ranged from 48 to 95 years, and was 82.7% White; the MR1 diary + fMRI sample was smaller (*N* = 48), younger (range = 29–70), and 93.8% White. The current analytic summaries characterize age, recorded binary sex, and race, but not gender identity, ethnicity, nativity or immigration history, socioeconomic status, disability, sexual orientation, or clinical diagnoses. Generalizability to younger people, people with other demographic or clinical characteristics, non-U.S. contexts, and people less able or willing to complete intensive longitudinal and imaging protocols is therefore uncertain. The findings may also depend on the passive image-viewing task, the negative, neutral, and positive stimuli and neutral-face probes, the diary affect measures, scanner acquisition and preprocessing choices, region-of-interest definitions, and the cross-run persistence and beta-series connectivity calculations. The general associations could be tested with other stimuli, scanners, and intensive affect measures, but direct replications should preserve the task timing, image-to-face sequence, region definitions, persistence pairing scheme, connectivity contrast, and quality-control criteria. Larger lifespan, longitudinal, and more demographically and culturally diverse samples are needed to establish which results generalize beyond these cohorts and methods.

## Funding and Acknowledgements

The MIDUS study has been funded by the John D. and Catherine T. MacArthur Foundation Research Network and the National Institute on Aging (P01 AG020166 and U19 AG051426). MIDUS 3 neuroscience data collection was additionally supported by the Eunice Kennedy Shriver National Institute of Child Health and Human Development (P50 HD105353). This work was supported by the National Institute on Aging (K01 AG086665 to Adam Turnbull). The content is solely the responsibility of the authors and does not necessarily represent the official views of the National Institutes of Health.

