## Supplementary files for "Amygdala Persistence in Aging: Replication and Extension of Puccetti et al. (2021)"

M3 primary adjusted analyses use mixed-effects models (MLM); MR1 uses OLS. Main tables include untransformed primary outcomes and explicitly labeled exploratory moderation. Log-NA robustness, alternative samples/measures, and motion checks appear in sensitivity tables. Statistical indirect effects and their age sensitivity remain together in Table S11.

All coefficients are unstandardized. Confidence intervals are two-sided 95%: normal-Wald for MLM, Student-*t* for OLS and condition means/differences, Fisher-*z* for Pearson correlations, and family-level percentile bootstrap for indirect effects. *p* (analysis) preserves the saved directional or two-sided test; *p (2-sided) is also shown except for the directional mediation analyses in Table S11.* Each method retains its own complete-case *N*. Comparisons are not independent replications or a basis for selecting the primary method. MR1 OLS estimates are repeated in comparison tables for convenience.

PANAS negative affect uses the natural log without an added constant; diary negative affect retains its separate offset log transformation. Exploratory and sensitivity *p* values are nominal, without multiplicity adjustment. Unavailable CIs indicate unavailable inference, not nonsignificance. Correlations have no counterpart for interaction coefficients or motion-adjusted regressions.

Contents: S1–S2 main results; S3–S4 sensitivity results; S5–S6 main-method comparisons; S7–S8 sensitivity-method comparisons; S9 contextual associations; S10 task-condition results; S11 statistical indirect effects. Supplementary Figure S1 follows the tables.

### Table S1. M3 main results: mixed-effects models

Moderation rows are exploratory. Main versus sensitivity organization does not change preregistration status.

| Analysis / specification | Predictor / interaction | Outcome | *N* | β | 95% CI | *p* (analysis) | *p* (2-sided) |
|---|---|---|---|---|---|---|---|
| Age and affect | Age (diary) | Diary PA | 1168 | 0.00987 | [0.00593, 0.01381] | < .001 | < .001 |
| Age and affect | Age (diary) | Diary NA | 1168 | −0.00285 | [−0.00422, −0.00147] | < .001 | < .001 |
| Persistence and affect | Left negative persistence | Diary PA | 81 | −0.69957 | [−1.63263, 0.23349] | .0708 | .1417 |
| Persistence and affect | Left negative persistence | Diary NA | 81 | 0.27793 | [0.01191, 0.54394] | .0203 | .0406 |
| Age and persistence | Age (neuroscience) | Left negative persistence | 127 | −0.00260 | [−0.00504, −0.00016] | .0183 | .0367 |
| Connectivity and affect | Left amygdala-anterior vmPFC (negative − neutral) | Diary PA | 81 | −0.01776 | [−0.45273, 0.41722] | .5319 | .9362 |
| Connectivity and affect | Left amygdala-anterior vmPFC (negative − neutral) | Diary NA | 81 | −0.10554 | [−0.22329, 0.01221] | .0395 | .0790 |
| Connectivity and affect | Left amygdala-posterior vmPFC (negative − neutral) | Diary PA | 81 | −0.35253 | [−0.64851, −0.05655] | .9902 | .0196 |
| Connectivity and affect | Left amygdala-posterior vmPFC (negative − neutral) | Diary NA | 81 | −0.05903 | [−0.16756, 0.04950] | .1432 | .2864 |
| Connectivity and persistence | Left amygdala-anterior vmPFC (negative − neutral) | Left negative persistence | 127 | −0.01802 | [−0.08493, 0.04890] | .2989 | .5977 |
| Connectivity and persistence | Left amygdala-posterior vmPFC (negative − neutral) | Left negative persistence | 127 | 0.01094 | [−0.05584, 0.07772] | .6259 | .7481 |
| Exploratory persistence moderation / reappraisal | Left negative persistence × Reappraisal | Diary PA | 80 | 0.99955 | [0.02769, 1.97140] | .0438 | .0438 |
| Exploratory persistence moderation / reappraisal | Left negative persistence × Reappraisal | Diary NA | 80 | −0.20428 | [−0.46145, 0.05290] | .1195 | .1195 |
| Exploratory persistence moderation / suppression | Left negative persistence × Suppression | Diary PA | 80 | 0.58628 | [0.28959, 0.88297] | < .001 | < .001 |
| Exploratory persistence moderation / suppression | Left negative persistence × Suppression | Diary NA | 80 | 0.09257 | [−0.15418, 0.33933] | .4622 | .4622 |
| Exploratory connectivity moderation / reappraisal | Left amygdala-anterior vmPFC (negative − neutral) × Reappraisal | Diary PA | 80 | −0.35083 | [−0.82992, 0.12827] | .1512 | .1512 |
| Exploratory connectivity moderation / reappraisal | Left amygdala-anterior vmPFC (negative − neutral) × Reappraisal | Diary NA | 80 | 0.02431 | [−0.11945, 0.16806] | .7404 | .7404 |

| **Analysis / specification** | **Predictor / interaction** | **Outcome** | ***N*** | **β** | **95% CI** | ***p* (analysis)** | ***p* (2-sided)** |
|---|---|---|---|---|---|---|---|
| Exploratory connectivity moderation / reappraisal | Left amygdala-posterior vmPFC (negative − neutral) × Reappraisal | Diary PA | 80 | −0.04084 | [−0.47517, 0.39349] | .8538 | .8538 |
| Exploratory connectivity moderation / reappraisal | Left amygdala-posterior vmPFC (negative − neutral) × Reappraisal | Diary NA | 80 | −0.00005 | [−0.11830, 0.11820] | .9994 | .9994 |
| Exploratory connectivity moderation / suppression | Left amygdala-anterior vmPFC (negative − neutral) × Suppression | Diary PA | 80 | 0.11799 | [−0.24419, 0.48017] | .5231 | .5231 |
| Exploratory connectivity moderation / suppression | Left amygdala-anterior vmPFC (negative − neutral) × Suppression | Diary NA | 80 | −0.01365 | [−0.11695, 0.08964] | .7956 | .7956 |
| Exploratory connectivity moderation / suppression | Left amygdala-posterior vmPFC (negative − neutral) × Suppression | Diary PA | 80 | −0.02157 | [−0.27275, 0.22961] | .8663 | .8663 |
| Exploratory connectivity moderation / suppression | Left amygdala-posterior vmPFC (negative − neutral) × Suppression | Diary NA | 80 | 0.09747 | [0.00117, 0.19378] | .0473 | .0473 |

### Table S2. MR1 main results: OLS models

MR1 is a targeted replication; moderation and mediation were not fitted.

| Analysis / specification | Predictor / interaction | Outcome | *N* | β | 95% CI | *p* (analysis) | *p* (2-sided) |
|---|---|---|---|---|---|---|---|
| Persistence and affect | Left negative persistence | Diary PA | 48 | −0.19538 | [−1.36712, 0.97635] | .3690 | .7380 |
| Persistence and affect | Left negative persistence | Diary NA | 48 | 0.31133 | [−0.12922, 0.75188] | .0806 | .1611 |
| Age and persistence | Age (neuroscience) | Left negative persistence | 115 | 0.00134 | [−0.00157, 0.00425] | .8177 | .3646 |
| Connectivity and affect | Left amygdala-anterior vmPFC (negative − neutral) | Diary PA | 48 | 0.05918 | [−0.47246, 0.59083] | .4116 | .8232 |
| Connectivity and affect | Left amygdala-anterior vmPFC (negative − neutral) | Diary NA | 48 | −0.08013 | [−0.28321, 0.12294] | .2150 | .4301 |
| Connectivity and affect | Left amygdala-posterior vmPFC (negative − neutral) | Diary PA | 48 | −0.14926 | [−0.75131, 0.45279] | .6904 | .6193 |
| Connectivity and affect | Left amygdala-posterior vmPFC (negative − neutral) | Diary NA | 48 | 0.02214 | [−0.21006, 0.25433] | .5759 | .8483 |
| Connectivity and persistence | Left amygdala-anterior vmPFC (negative − neutral) | Left negative persistence | 115 | 0.03129 | [−0.06716, 0.12974] | .7350 | .5300 |
| Connectivity and persistence | Left amygdala-posterior vmPFC (negative − neutral) | Left negative persistence | 115 | −0.01751 | [−0.11901, 0.08399] | .3665 | .7330 |

### Table S3. M3 sensitivity results: mixed-effects models

The negative-condition anterior FC sensitivity required a targeted increase to 2,000 optimizer iterations with unchanged model/sample. The persistence × reappraisal raw-PANAS-NA MLM has invalid fitted covariance: affected inference is unavailable despite alternative-optimizer checks. Coefficients are retained for transparency.

| Analysis / specification | Predictor / interaction | Outcome | *N* | β | 95% CI | *p* (analysis) | *p* (2-sided) |
|---|---|---|---|---|---|---|---|
| Age and affect / log-NA robustness | Age (diary) | Diary log NA | 1168 | −0.02218 | [−0.03040, −0.01395] | < .001 | < .001 |
| Age and affect / Neuroscience subsample | Age (neuroscience) | Diary PA | 137 | 0.00829 | [−0.00495, 0.02152] | .1098 | .2197 |
| Age and affect / Neuroscience subsample | Age (neuroscience) | Diary NA | 137 | −0.00215 | [−0.00728, 0.00298] | .2054 | .4107 |
| Age and affect / Neuroscience subsample / log-NA robustness | Age (neuroscience) | Diary log NA | 137 | −0.03191 | [−0.06005, −0.00377] | .0131 | .0263 |
| Age and affect / PANAS | Age (neuroscience) | PANAS PA | 230 | 0.00117 | [−0.00854, 0.01088] | .4065 | .8130 |
| Age and affect / PANAS | Age (neuroscience) | PANAS NA | 230 | −0.00503 | [−0.01116, 0.00110] | .0540 | .1080 |
| Age and affect / PANAS / log-NA robustness | Age (neuroscience) | PANAS log NA | 230 | −0.00395 | [−0.00769, −0.00022] | .0190 | .0379 |
| Persistence and affect / log-NA robustness | Left negative persistence | Diary log NA | 81 | 1.95009 | [−0.54517, 4.44534] | .0628 | .1256 |
| Persistence and affect / Positive persistence | Left positive persistence | Diary PA | 81 | −0.36717 | [−1.24859, 0.51425] | .4142 | .4142 |
| Persistence and affect / Positive persistence | Left positive persistence | Diary NA | 81 | −0.06017 | [−0.34768, 0.22733] | .6817 | .6817 |
| Persistence and affect / Positive persistence / log-NA robustness | Left positive persistence | Diary log NA | 81 | −1.76772 | [−4.60874, 1.07329] | .2226 | .2226 |
| Persistence and affect / PANAS | Left negative persistence | PANAS PA | 127 | −0.59143 | [−1.44776, 0.26490] | .0879 | .1758 |
| Persistence and affect / PANAS | Left negative persistence | PANAS NA | 127 | −0.05243 | [−0.76512, 0.66025] | .5573 | .8853 |
| Persistence and affect / PANAS / log-NA robustness | Left negative persistence | PANAS log NA | 127 | 0.00058 | [−0.35460, 0.35576] | .4987 | .9974 |
| Persistence and affect / Right hemisphere | Right negative persistence | Diary PA | 81 | −0.74982 | [−1.59900, 0.09936] | .0835 | .0835 |
| Persistence and affect / Right hemisphere | Right negative persistence | Diary NA | 81 | −0.03381 | [−0.33986, 0.27224] | .8286 | .8286 |
| Persistence and affect / Right hemisphere / log-NA robustness | Right negative persistence | Diary log NA | 81 | 0.68558 | [−2.05227, 3.42344] | .6236 | .6236 |

| Analysis / specification | Predictor / interaction | Outcome | *N* | β | 95% CI | *p* (analysis) | *p* (2-sided) |
|---|---|---|---|---|---|---|---|
| Persistence and affect / Regional activation | Left amygdala neg image | Diary PA | 81 | 0.93873 | [−0.12523, 2.00269] | .0838 | .0838 |
| Persistence and affect / Regional activation | Left amygdala neg image | Diary NA | 81 | −0.09807 | [−0.44747, 0.25134] | .5823 | .5823 |
| Persistence and affect / Regional activation / log-NA robustness | Left amygdala neg image | Diary log NA | 81 | −0.60774 | [−3.78167, 2.56618] | .7074 | .7074 |
| Persistence and affect / Regional activation | anterior vmPFC neg image | Diary PA | 81 | 0.23470 | [−0.57856, 1.04795] | .5716 | .5716 |
| Persistence and affect / Regional activation | anterior vmPFC neg image | Diary NA | 81 | −0.00780 | [−0.28636, 0.27076] | .9562 | .9562 |
| Persistence and affect / Regional activation / log-NA robustness | anterior vmPFC neg image | Diary log NA | 81 | 0.67452 | [−1.72004, 3.06907] | .5809 | .5809 |
| Persistence and affect / Regional activation | posterior vmPFC neg image | Diary PA | 81 | 0.61858 | [−0.35445, 1.59160] | .2128 | .2128 |
| Persistence and affect / Regional activation | posterior vmPFC neg image | Diary NA | 81 | −0.17524 | [−0.46787, 0.11738] | .2405 | .2405 |
| Persistence and affect / Regional activation / log-NA robustness | posterior vmPFC neg image | Diary log NA | 81 | 0.34106 | [−2.28370, 2.96583] | .7990 | .7990 |
| Persistence and affect / vmPFC persistence | ant vmPFC neg image-to-face persistence (Fisher *z*) | Diary PA | 81 | 0.13460 | [−0.44473, 0.71394] | .6488 | .6488 |
| Persistence and affect / vmPFC persistence | ant vmPFC neg image-to-face persistence (Fisher *z*) | Diary NA | 81 | −0.11027 | [−0.31377, 0.09324] | .2882 | .2882 |
| Persistence and affect / vmPFC persistence / log-NA robustness | ant vmPFC neg image-to-face persistence (Fisher *z*) | Diary log NA | 81 | −0.84991 | [−2.66417, 0.96435] | .3585 | .3585 |
| Persistence and affect / vmPFC persistence | post vmPFC neg image-to-face persistence (Fisher *z*) | Diary PA | 81 | −0.49097 | [−1.11188, 0.12993] | .1212 | .1212 |
| Persistence and affect / vmPFC persistence | post vmPFC neg image-to-face persistence (Fisher *z*) | Diary NA | 81 | 0.03910 | [−0.20669, 0.28490] | .7552 | .7552 |
| Persistence and affect / vmPFC persistence / log-NA robustness | post vmPFC neg image-to-face persistence (Fisher *z*) | Diary log NA | 81 | 1.44246 | [−0.77593, 3.66084] | .2025 | .2025 |
| Age and persistence / Positive persistence | Age (neuroscience) | Left positive persistence | 127 | −0.00183 | [−0.00439, 0.00072] | .1590 | .1590 |
| Age and persistence / Positive persistence | Age (neuroscience) | Right positive persistence | 127 | −0.00003 | [−0.00235, 0.00229] | .9786 | .9786 |
| Age and persistence / Right hemisphere | Age (neuroscience) | Right negative persistence | 127 | 0.00043 | [−0.00194, 0.00279] | .7235 | .7235 |

| **Analysis / specification** | **Predictor / interaction** | **Outcome** | ***N*** | **β** | **95% CI** | ***p* (analysis)** | ***p* (2-sided)** |
|---|---|---|---|---|---|---|---|
| Age and persistence / vmPFC persistence | Age (neuroscience) | ant vmPFC neg image-to-face persistence (Fisher *z*) | 127 | −0.00106 | [−0.00421, 0.00209] | .5096 | .5096 |
| Age and persistence / vmPFC persistence | Age (neuroscience) | post vmPFC neg image-to-face persistence (Fisher *z*) | 127 | 0.00016 | [−0.00245, 0.00278] | .9020 | .9020 |
| Connectivity and affect / log-NA robustness | Left amygdala-anterior vmPFC (negative − neutral) | Diary log NA | 81 | −1.06334 | [−2.01370, −0.11298] | .0142 | .0283 |
| Connectivity and affect / log-NA robustness | Left amygdala-posterior vmPFC (negative − neutral) | Diary log NA | 81 | −0.56183 | [−1.54616, 0.42250] | .1316 | .2633 |
| Connectivity and affect / Negative condition | Left amygdala-anterior vmPFC neg | Diary PA | 81 | 0.13912 | [−0.28623, 0.56446] | .2607 | .5215 |
| Connectivity and affect / Negative condition | Left amygdala-anterior vmPFC neg | Diary NA | 81 | −0.06067 | [−0.19598, 0.07463] | .1897 | .3795 |
| Connectivity and affect / Negative condition / log-NA robustness | Left amygdala-anterior vmPFC neg | Diary log NA | 81 | −0.81772 | [−1.92842, 0.29298] | .0745 | .1490 |
| Connectivity and affect / Negative condition | Left amygdala-posterior vmPFC neg | Diary PA | 81 | −0.11265 | [−0.59341, 0.36812] | .6770 | .6461 |
| Connectivity and affect / Negative condition | Left amygdala-posterior vmPFC neg | Diary NA | 81 | −0.03720 | [−0.18158, 0.10718] | .3068 | .6136 |
| Connectivity and affect / Negative condition / log-NA robustness | Left amygdala-posterior vmPFC neg | Diary log NA | 81 | −0.65313 | [−1.92492, 0.61867] | .1571 | .3142 |
| Connectivity and affect / PANAS | Left amygdala-anterior vmPFC (negative − neutral) | PANAS PA | 127 | −0.15668 | [−0.46852, 0.15515] | .8376 | .3247 |
| Connectivity and affect / PANAS | Left amygdala-anterior vmPFC (negative − neutral) | PANAS NA | 127 | −0.25090 | [−0.46676, −0.03504] | .0114 | .0227 |
| Connectivity and affect / PANAS / log-NA robustness | Left amygdala-anterior vmPFC (negative − neutral) | PANAS log NA | 127 | −0.13900 | [−0.26397, −0.01404] | .0146 | .0292 |
| Connectivity and affect / PANAS | Left amygdala-posterior vmPFC (negative − neutral) | PANAS PA | 127 | 0.07033 | [−0.24024, 0.38089] | .3286 | .6572 |
| Connectivity and affect / PANAS | Left amygdala-posterior vmPFC (negative − neutral) | PANAS NA | 127 | −0.24937 | [−0.46305, −0.03569] | .0111 | .0222 |

| Analysis / specification | Predictor / interaction | Outcome | *N* | β | 95% CI | *p* (analysis) | *p* (2-sided) |
|---|---|---|---|---|---|---|---|
| Connectivity and affect / PANAS / log-NA robustness | Left amygdala-posterior vmPFC (negative − neutral) | PANAS log NA | 127 | −0.12779 | [−0.25278, −0.00281] | .0225 | .0451 |
| Connectivity and affect / Positive minus neutral | Left amygdala-anterior vmPFC (positive − neutral) | Diary PA | 81 | −0.07700 | [−0.40660, 0.25260] | .6470 | .6470 |
| Connectivity and affect / Positive minus neutral | Left amygdala-anterior vmPFC (positive − neutral) | Diary NA | 81 | −0.01756 | [−0.11318, 0.07807] | .7189 | .7189 |
| Connectivity and affect / Positive minus neutral / log-NA robustness | Left amygdala-anterior vmPFC (positive − neutral) | Diary log NA | 81 | −0.40321 | [−1.24579, 0.43937] | .3483 | .3483 |
| Connectivity and affect / Positive minus neutral | Left amygdala-posterior vmPFC (positive − neutral) | Diary PA | 81 | −0.00587 | [−0.36823, 0.35648] | .9747 | .9747 |
| Connectivity and affect / Positive minus neutral | Left amygdala-posterior vmPFC (positive − neutral) | Diary NA | 81 | −0.02144 | [−0.13503, 0.09214] | .7114 | .7114 |
| Connectivity and affect / Positive minus neutral / log-NA robustness | Left amygdala-posterior vmPFC (positive − neutral) | Diary log NA | 81 | −0.12354 | [−1.13950, 0.89242] | .8116 | .8116 |
| Connectivity and affect / Right amygdala | Right amygdala-anterior vmPFC (negative − neutral) | Diary PA | 81 | 0.17369 | [−0.25073, 0.59811] | .4225 | .4225 |
| Connectivity and affect / Right amygdala | Right amygdala-anterior vmPFC (negative − neutral) | Diary NA | 81 | 0.05226 | [−0.06955, 0.17406] | .4004 | .4004 |
| Connectivity and affect / Right amygdala / log-NA robustness | Right amygdala-anterior vmPFC (negative − neutral) | Diary log NA | 81 | 0.69201 | [−0.37886, 1.76289] | .2053 | .2053 |
| Connectivity and affect / Right amygdala | Right amygdala-posterior vmPFC (negative − neutral) | Diary PA | 81 | −0.24155 | [−0.75139, 0.26829] | .3531 | .3531 |
| Connectivity and affect / Right amygdala | Right amygdala-posterior vmPFC (negative − neutral) | Diary NA | 81 | 0.04267 | [−0.08623, 0.17158] | .5165 | .5165 |
| Connectivity and affect / Right amygdala / log-NA robustness | Right amygdala-posterior vmPFC (negative − neutral) | Diary log NA | 81 | 0.67874 | [−0.43439, 1.79188] | .2320 | .2320 |
| Anterior/posterior contrast / diary | Left amygdala-ant minus posterior vmPFC (negative − neutral) | Diary PA | 81 | 0.47102 | [0.10314, 0.83890] | .0060 | .0121 |
| Anterior/posterior contrast / diary | Left amygdala-ant minus posterior vmPFC (negative − neutral) | Diary NA | 81 | −0.03859 | [−0.15778, 0.08060] | .2628 | .5257 |

| Analysis / specification | Predictor / interaction | Outcome | *N* | β | 95% CI | *p* (analysis) | *p* (2-sided) |
|---|---|---|---|---|---|---|---|
| Anterior/posterior contrast / diary / log-NA robustness | Left amygdala-ant minus posterior vmPFC (negative − neutral) | Diary log NA | 81 | −0.58900 | [−1.66077, 0.48277] | .1407 | .2814 |
| Anterior/posterior contrast / diary | Right amygdala-ant minus posterior vmPFC (negative − neutral) | Diary PA | 81 | 0.29776 | [−0.07216, 0.66769] | .0573 | .1147 |
| Anterior/posterior contrast / diary | Right amygdala-ant minus posterior vmPFC (negative − neutral) | Diary NA | 81 | 0.01373 | [−0.10662, 0.13408] | .5885 | .8231 |
| Anterior/posterior contrast / diary / log-NA robustness | Right amygdala-ant minus posterior vmPFC (negative − neutral) | Diary log NA | 81 | 0.07193 | [−1.00376, 1.14763] | .5521 | .8957 |
| Anterior/posterior contrast / panas | Left amygdala-ant minus posterior vmPFC (negative − neutral) | PANAS PA | 127 | −0.26703 | [−0.60453, 0.07047] | .9395 | .1210 |
| Anterior/posterior contrast / panas | Left amygdala-ant minus posterior vmPFC (negative − neutral) | PANAS NA | 127 | 0.00344 | [−0.22774, 0.23462] | .5116 | .9767 |
| Anterior/posterior contrast / panas / log-NA robustness | Left amygdala-ant minus posterior vmPFC (negative − neutral) | PANAS log NA | 127 | −0.01004 | [−0.14480, 0.12473] | .4420 | .8839 |
| Anterior/posterior contrast / panas | Right amygdala-ant minus posterior vmPFC (negative − neutral) | PANAS PA | 127 | −0.14694 | [−0.49568, 0.20179] | .7956 | .4089 |
| Anterior/posterior contrast / panas | Right amygdala-ant minus posterior vmPFC (negative − neutral) | PANAS NA | 127 | −0.01643 | [−0.31553, 0.28268] | .4571 | .9143 |
| Anterior/posterior contrast / panas / log-NA robustness | Right amygdala-ant minus posterior vmPFC (negative − neutral) | PANAS log NA | 127 | −0.01461 | [−0.16166, 0.13245] | .4228 | .8456 |
| Connectivity and persistence / Negative condition | Left amygdala-anterior vmPFC neg | Left negative persistence | 127 | 0.00099 | [−0.07876, 0.08074] | .5097 | .9806 |
| Connectivity and persistence / Negative condition | Left amygdala-posterior vmPFC neg | Left negative persistence | 127 | 0.01618 | [−0.07139, 0.10375] | .6414 | .7172 |
| Connectivity and persistence / Positive persistence | Left amygdala-anterior vmPFC (negative − neutral) | Left positive persistence | 127 | −0.01010 | [−0.07770, 0.05750] | .7696 | .7696 |

| **Analysis / specification** | **Predictor / interaction** | **Outcome** | ***N*** | **β** | **95% CI** | ***p* (analysis)** | ***p* (2-sided)** |
|---|---|---|---|---|---|---|---|
| Connectivity and persistence / Positive persistence | Left amygdala-posterior vmPFC (negative − neutral) | Left positive persistence | 127 | −0.02150 | [−0.08815, 0.04514] | .5271 | .5271 |
| Connectivity and persistence / Positive minus neutral | Left amygdala-anterior vmPFC (positive − neutral) | Left negative persistence | 127 | 0.01615 | [−0.04739, 0.07969] | .6184 | .6184 |
| Connectivity and persistence / Positive minus neutral | Left amygdala-posterior vmPFC (positive − neutral) | Left negative persistence | 127 | 0.01818 | [−0.05333, 0.08968] | .6183 | .6183 |
| Connectivity and persistence / Right amygdala | Right amygdala-anterior vmPFC (negative − neutral) | Left negative persistence | 127 | −0.00894 | [−0.08245, 0.06457] | .8116 | .8116 |
| Connectivity and persistence / Right amygdala | Right amygdala-posterior vmPFC (negative − neutral) | Left negative persistence | 127 | −0.01104 | [−0.08577, 0.06368] | .7720 | .7720 |
| Connectivity and persistence / Right hemisphere | Left amygdala-anterior vmPFC (negative − neutral) | Right negative persistence | 127 | −0.03466 | [−0.09793, 0.02861] | .2830 | .2830 |
| Connectivity and persistence / Right hemisphere | Left amygdala-posterior vmPFC (negative − neutral) | Right negative persistence | 127 | −0.00576 | [−0.07012, 0.05860] | .8608 | .8608 |
| Potential-overlap exclusion subsample | Left negative persistence | Diary PA | 55 | −0.54763 | [−1.42895, 0.33369] | .1116 | .2233 |
| Potential-overlap exclusion subsample | Left negative persistence | Diary NA | 55 | 0.22297 | [−0.04340, 0.48935] | .0504 | .1009 |
| Potential-overlap exclusion subsample / log-NA robustness | Left negative persistence | Diary log NA | 55 | 1.89113 | [−0.79940, 4.58167] | .0842 | .1683 |
| Exploratory persistence moderation / reappraisal / log-NA robustness | Left negative persistence × Reappraisal | Diary log NA | 80 | −1.52805 | [−3.84325, 0.78714] | .1958 | .1958 |
| Exploratory persistence moderation / PANAS / reappraisal | Left negative persistence × Reappraisal | PANAS PA | 126 | 0.00423 | [−0.77819, 0.78666] | .9915 | .9915 |
| Exploratory persistence moderation / PANAS / reappraisal | Left negative persistence × Reappraisal | PANAS NA | 126 | −0.25886 | Unavailable | — | — |
| Exploratory persistence moderation / PANAS / reappraisal / log-NA robustness | Left negative persistence × Reappraisal | PANAS log NA | 126 | −0.16159 | [−0.48817, 0.16499] | .3322 | .3322 |
| Exploratory persistence moderation / PANAS / suppression | Left negative persistence × Suppression | PANAS PA | 126 | 0.36977 | [−0.36026, 1.09979] | .3208 | .3208 |

| **Analysis / specification** | **Predictor / interaction** | **Outcome** | ***N*** | **β** | **95% CI** | ***p* (analysis)** | ***p* (2-sided)** |
|---|---|---|---|---|---|---|---|
| Exploratory persistence moderation / PANAS / suppression | Left negative persistence × Suppression | PANAS NA | 126 | −0.09392 | [−0.60908, 0.42123] | .7208 | .7208 |
| Exploratory persistence moderation / PANAS / suppression / log-NA robustness | Left negative persistence × Suppression | PANAS log NA | 126 | −0.02756 | [−0.32627, 0.27115] | .8565 | .8565 |
| Exploratory persistence moderation / Right hemisphere / reappraisal | Right negative persistence × Reappraisal | Diary PA | 80 | 0.52373 | [−0.54615, 1.59362] | .3373 | .3373 |
| Exploratory persistence moderation / Right hemisphere / reappraisal | Right negative persistence × Reappraisal | Diary NA | 80 | 0.00012 | [−0.29960, 0.29985] | .9994 | .9994 |
| Exploratory persistence moderation / Right hemisphere / reappraisal / log-NA robustness | Right negative persistence × Reappraisal | Diary log NA | 80 | −1.14875 | [−3.75315, 1.45564] | .3873 | .3873 |
| Exploratory persistence moderation / Right hemisphere / suppression | Right negative persistence × Suppression | Diary PA | 80 | 0.44114 | [−0.45640, 1.33869] | .3354 | .3354 |
| Exploratory persistence moderation / Right hemisphere / suppression | Right negative persistence × Suppression | Diary NA | 80 | −0.11803 | [−0.40470, 0.16864] | .4197 | .4197 |
| Exploratory persistence moderation / Right hemisphere / suppression / log-NA robustness | Right negative persistence × Suppression | Diary log NA | 80 | −0.16730 | [−2.70571, 2.37112] | .8972 | .8972 |
| Exploratory persistence moderation / suppression / log-NA robustness | Left negative persistence × Suppression | Diary log NA | 80 | 0.31763 | [−1.91546, 2.55071] | .7804 | .7804 |
| Exploratory connectivity moderation / reappraisal / log-NA robustness | Left amygdala-anterior vmPFC (negative − neutral) × Reappraisal | Diary log NA | 80 | 0.29688 | [−0.89337, 1.48713] | .6249 | .6249 |
| Exploratory connectivity moderation / reappraisal / log-NA robustness | Left amygdala-posterior vmPFC (negative − neutral) × Reappraisal | Diary log NA | 80 | 0.02600 | [−0.98774, 1.03974] | .9599 | .9599 |
| Exploratory connectivity moderation / PANAS / reappraisal | Left amygdala-anterior vmPFC (negative − neutral) × Reappraisal | PANAS PA | 126 | −0.01263 | [−0.31937, 0.29411] | .9357 | .9357 |

| Analysis / specification | Predictor / interaction | Outcome | *N* | β | 95% CI | *p* (analysis) | *p* (2-sided) |
|---|---|---|---|---|---|---|---|
| Exploratory connectivity moderation / PANAS / reappraisal | Left amygdala-anterior vmPFC (negative − neutral) × Reappraisal | PANAS NA | 126 | −0.11240 | [−0.33666, 0.11186] | .3259 | .3259 |
| Exploratory connectivity moderation / PANAS / reappraisal / log-NA robustness | Left amygdala-anterior vmPFC (negative − neutral) × Reappraisal | PANAS log NA | 126 | −0.05319 | [−0.18573, 0.07935] | .4315 | .4315 |
| Exploratory connectivity moderation / PANAS / reappraisal | Left amygdala-posterior vmPFC (negative − neutral) × Reappraisal | PANAS PA | 126 | 0.07756 | [−0.23475, 0.38988] | .6264 | .6264 |
| Exploratory connectivity moderation / PANAS / reappraisal | Left amygdala-posterior vmPFC (negative − neutral) × Reappraisal | PANAS NA | 126 | −0.15728 | [−0.37588, 0.06131] | .1585 | .1585 |
| Exploratory connectivity moderation / PANAS / reappraisal / log-NA robustness | Left amygdala-posterior vmPFC (negative − neutral) × Reappraisal | PANAS log NA | 126 | −0.06514 | [−0.19365, 0.06336] | .3204 | .3204 |
| Exploratory connectivity moderation / PANAS / suppression | Left amygdala-anterior vmPFC (negative − neutral) × Suppression | PANAS PA | 126 | −0.06674 | [−0.34337, 0.20989] | .6363 | .6363 |
| Exploratory connectivity moderation / PANAS / suppression | Left amygdala-anterior vmPFC (negative − neutral) × Suppression | PANAS NA | 126 | −0.13017 | [−0.33223, 0.07189] | .2067 | .2067 |
| Exploratory connectivity moderation / PANAS / suppression / log-NA robustness | Left amygdala-anterior vmPFC (negative − neutral) × Suppression | PANAS log NA | 126 | −0.05184 | [−0.16608, 0.06240] | .3738 | .3738 |
| Exploratory connectivity moderation / PANAS / suppression | Left amygdala-posterior vmPFC (negative − neutral) × Suppression | PANAS PA | 126 | 0.09530 | [−0.19135, 0.38195] | .5147 | .5147 |
| Exploratory connectivity moderation / PANAS / suppression | Left amygdala-posterior vmPFC (negative − neutral) × Suppression | PANAS NA | 126 | −0.24552 | [−0.44201, −0.04903] | .0143 | .0143 |
| Exploratory connectivity moderation / PANAS / suppression / log-NA robustness | Left amygdala-posterior vmPFC (negative − neutral) × Suppression | PANAS log NA | 126 | −0.12808 | [−0.24286, −0.01329] | .0287 | .0287 |
| Exploratory connectivity moderation / Right amygdala / reappraisal | Right amygdala-anterior vmPFC (negative − neutral) × Reappraisal | Diary PA | 80 | 0.21591 | [−0.16660, 0.59841] | .2686 | .2686 |

| Analysis / specification | Predictor / interaction | Outcome | *N* | β | 95% CI | *p* (analysis) | *p* (2-sided) |
|---|---|---|---|---|---|---|---|
| Exploratory connectivity moderation / Right amygdala / reappraisal | Right amygdala-anterior vmPFC (negative − neutral) × Reappraisal | Diary NA | 80 | −0.02538 | [−0.12325, 0.07249] | .6112 | .6112 |
| Exploratory connectivity moderation / Right amygdala / reappraisal / log-NA robustness | Right amygdala-anterior vmPFC (negative − neutral) × Reappraisal | Diary log NA | 80 | −0.33339 | [−1.18470, 0.51792] | .4427 | .4427 |
| Exploratory connectivity moderation / Right amygdala / reappraisal | Right amygdala-posterior vmPFC (negative − neutral) × Reappraisal | Diary PA | 80 | 0.12484 | [−0.38712, 0.63680] | .6327 | .6327 |
| Exploratory connectivity moderation / Right amygdala / reappraisal | Right amygdala-posterior vmPFC (negative − neutral) × Reappraisal | Diary NA | 80 | 0.03659 | [−0.11059, 0.18377] | .6261 | .6261 |
| Exploratory connectivity moderation / Right amygdala / reappraisal / log-NA robustness | Right amygdala-posterior vmPFC (negative − neutral) × Reappraisal | Diary log NA | 80 | −0.58786 | [−1.90226, 0.72653] | .3807 | .3807 |
| Exploratory connectivity moderation / Right amygdala / suppression | Right amygdala-anterior vmPFC (negative − neutral) × Suppression | Diary PA | 80 | 0.24482 | [−0.09860, 0.58824] | .1623 | .1623 |
| Exploratory connectivity moderation / Right amygdala / suppression | Right amygdala-anterior vmPFC (negative − neutral) × Suppression | Diary NA | 80 | −0.06553 | [−0.16517, 0.03411] | .1974 | .1974 |
| Exploratory connectivity moderation / Right amygdala / suppression / log-NA robustness | Right amygdala-anterior vmPFC (negative − neutral) × Suppression | Diary log NA | 80 | −0.49492 | [−1.30919, 0.31934] | .2335 | .2335 |
| Exploratory connectivity moderation / Right amygdala / suppression | Right amygdala-posterior vmPFC (negative − neutral) × Suppression | Diary PA | 80 | −0.08361 | [−0.44563, 0.27842] | .6508 | .6508 |
| Exploratory connectivity moderation / Right amygdala / suppression | Right amygdala-posterior vmPFC (negative − neutral) × Suppression | Diary NA | 80 | 0.03432 | [−0.06446, 0.13310] | .4959 | .4959 |
| Exploratory connectivity moderation / Right amygdala / suppression / log-NA robustness | Right amygdala-posterior vmPFC (negative − neutral) × Suppression | Diary log NA | 80 | −0.01224 | [−0.86721, 0.84274] | .9776 | .9776 |
| Exploratory connectivity moderation / suppression / log-NA robustness | Left amygdala-anterior vmPFC (negative − neutral) × Suppression | Diary log NA | 80 | 0.11896 | [−0.75680, 0.99473] | .7901 | .7901 |

| **Analysis / specification** | **Predictor / interaction** | **Outcome** | ***N*** | **β** | **95% CI** | ***p* (analysis)** | ***p* (2-sided)** |
|---|---|---|---|---|---|---|---|
| Exploratory connectivity moderation / suppression / log-NA robustness | Left amygdala-posterior vmPFC (negative − neutral) × Suppression | Diary log NA | 80 | 0.52335 | [−0.37033, 1.41703] | .2511 | .2511 |
| Motion association / fd mean across runs | fd mean across runs | Left negative persistence | 127 | 0.02165 | [−0.21609, 0.25940] | .8583 | .8583 |
| Motion association / fd mean across runs | fd mean across runs | Left amygdala-anterior vmPFC (negative − neutral) | 127 | −0.00507 | [−0.62958, 0.61943] | .9873 | .9873 |
| Motion association / fd mean across runs | fd mean across runs | Left amygdala-posterior vmPFC (negative − neutral) | 127 | 0.37606 | [−0.24693, 0.99904] | .2368 | .2368 |
| Adjusted for fd mean across runs | Left negative persistence | Diary NA | 81 | 0.24330 | [−0.02454, 0.51113] | .0375 | .0750 |
| Adjusted for fd mean across runs | Age (neuroscience) | Left negative persistence | 127 | −0.00265 | [−0.00515, −0.00016] | .0186 | .0372 |
| Adjusted for fd mean across runs | Left amygdala-anterior vmPFC (negative − neutral) | Diary NA | 81 | −0.09979 | [−0.21725, 0.01768] | .0480 | .0959 |
| Adjusted for fd mean across runs / log-NA robustness | Left amygdala-anterior vmPFC (negative − neutral) | Diary log NA | 81 | −1.02642 | [−1.98560, −0.06725] | .0180 | .0360 |
| Motion association / fd neg mean | fd neg mean | Left negative persistence | 127 | 0.01201 | [−0.22056, 0.24458] | .9194 | .9194 |
| Motion association / fd neg mean | fd neg mean | Left amygdala-anterior vmPFC (negative − neutral) | 127 | 0.16096 | [−0.45096, 0.77288] | .6062 | .6062 |
| Motion association / fd neg mean | fd neg mean | Left amygdala-posterior vmPFC (negative − neutral) | 127 | 0.47355 | [−0.13781, 1.08491] | .1290 | .1290 |
| Adjusted for fd neg mean | Left negative persistence | Diary NA | 81 | 0.26669 | [−0.00287, 0.53625] | .0262 | .0525 |
| Adjusted for fd neg mean | Age (neuroscience) | Left negative persistence | 127 | −0.00263 | [−0.00511, −0.00014] | .0193 | .0385 |
| Adjusted for fd neg mean | Left amygdala-anterior vmPFC (negative − neutral) | Diary NA | 81 | −0.10520 | [−0.22310, 0.01270] | .0402 | .0803 |
| Adjusted for fd neg mean / log-NA robustness | Left amygdala-anterior vmPFC (negative − neutral) | Diary log NA | 81 | −1.06198 | [−2.01974, −0.10422] | .0149 | .0298 |

### Table S4. MR1 sensitivity results: OLS models

MR1 sensitivity analyses retain OLS as the primary adjusted method.

| Analysis / specification | Predictor / interaction | Outcome | *N* | β | 95% CI | *p* (analysis) | *p* (2-sided) |
|---|---|---|---|---|---|---|---|
| Persistence and affect / log-NA robustness | Left negative persistence | Diary log NA | 48 | 1.42413 | [−1.10552, 3.95379] | .1311 | .2622 |
| Persistence and affect / PANAS | Left negative persistence | PANAS PA | 114 | 0.59713 | [−0.03547, 1.22973] | .9680 | .0640 |
| Persistence and affect / PANAS | Left negative persistence | PANAS NA | 114 | 0.12091 | [−0.38474, 0.62656] | .3182 | .6364 |
| Persistence and affect / PANAS / log-NA robustness | Left negative persistence | PANAS log NA | 114 | 0.08719 | [−0.21954, 0.39393] | .2871 | .5742 |
| Persistence and affect / Right hemisphere | Right negative persistence | Diary PA | 48 | −1.28052 | [−2.55468, −0.00635] | .0489 | .0489 |
| Persistence and affect / Right hemisphere | Right negative persistence | Diary NA | 48 | 0.38379 | [−0.11593, 0.88352] | .1286 | .1286 |
| Persistence and affect / Right hemisphere / log-NA robustness | Right negative persistence | Diary log NA | 48 | 1.31623 | [−1.58094, 4.21340] | .3642 | .3642 |
| Age and persistence / Positive persistence | Age (neuroscience) | Left positive persistence | 115 | −0.00034 | [−0.00263, 0.00194] | .7660 | .7660 |
| Age and persistence / Positive persistence | Age (neuroscience) | Right positive persistence | 115 | −0.00027 | [−0.00240, 0.00187] | .8060 | .8060 |
| Age and persistence / Right hemisphere | Age (neuroscience) | Right negative persistence | 115 | −0.00061 | [−0.00284, 0.00161] | .5865 | .5865 |
| Connectivity and affect / log-NA robustness | Left amygdala-anterior vmPFC (negative − neutral) | Diary log NA | 48 | −0.34007 | [−1.49995, 0.81982] | .2785 | .5570 |
| Connectivity and affect / log-NA robustness | Left amygdala-posterior vmPFC (negative − neutral) | Diary log NA | 48 | 0.38046 | [−0.93638, 1.69730] | .7186 | .5628 |
| Connectivity and affect / Negative condition | Left amygdala-anterior vmPFC neg | Diary PA | 48 | 0.31251 | [−0.41350, 1.03853] | .1949 | .3897 |
| Connectivity and affect / Negative condition | Left amygdala-anterior vmPFC neg | Diary NA | 48 | −0.05934 | [−0.34056, 0.22188] | .3361 | .6722 |
| Connectivity and affect / Negative condition / log-NA robustness | Left amygdala-anterior vmPFC neg | Diary log NA | 48 | −0.96107 | [−2.53647, 0.61433] | .1125 | .2250 |
| Connectivity and affect / Negative condition | Left amygdala-posterior vmPFC neg | Diary PA | 48 | 0.24936 | [−0.43862, 0.93735] | .2342 | .4683 |
| Connectivity and affect / Negative condition | Left amygdala-posterior vmPFC neg | Diary NA | 48 | −0.18924 | [−0.44885, 0.07036] | .0743 | .1486 |

| Analysis / specification | Predictor / interaction | Outcome | *N* | β | 95% CI | *p* (analysis) | *p* (2-sided) |
|---|---|---|---|---|---|---|---|
| Connectivity and affect / Negative condition / log-NA robustness | Left amygdala-posterior vmPFC neg | Diary log NA | 48 | −0.90237 | [−2.39167, 0.58693] | .1140 | .2281 |
| Connectivity and affect / PANAS | Left amygdala-anterior vmPFC (negative − neutral) | PANAS PA | 114 | 0.18617 | [−0.14272, 0.51505] | .1321 | .2643 |
| Connectivity and affect / PANAS | Left amygdala-anterior vmPFC (negative − neutral) | PANAS NA | 114 | −0.03829 | [−0.29864, 0.22206] | .3856 | .7712 |
| Connectivity and affect / PANAS / log-NA robustness | Left amygdala-anterior vmPFC (negative − neutral) | PANAS log NA | 114 | −0.01087 | [−0.16892, 0.14718] | .4459 | .8918 |
| Connectivity and affect / PANAS | Left amygdala-posterior vmPFC (negative − neutral) | PANAS PA | 114 | 0.22603 | [−0.11173, 0.56378] | .0937 | .1874 |
| Connectivity and affect / PANAS | Left amygdala-posterior vmPFC (negative − neutral) | PANAS NA | 114 | −0.10144 | [−0.36883, 0.16595] | .2268 | .4536 |
| Connectivity and affect / PANAS / log-NA robustness | Left amygdala-posterior vmPFC (negative − neutral) | PANAS log NA | 114 | −0.07360 | [−0.23568, 0.08849] | .1850 | .3701 |
| Connectivity and affect / Positive minus neutral | Left amygdala-anterior vmPFC (positive − neutral) | Diary PA | 48 | 0.11567 | [−0.39276, 0.62410] | .6483 | .6483 |
| Connectivity and affect / Positive minus neutral | Left amygdala-anterior vmPFC (positive − neutral) | Diary NA | 48 | −0.11839 | [−0.31089, 0.07410] | .2213 | .2213 |
| Connectivity and affect / Positive minus neutral / log-NA robustness | Left amygdala-anterior vmPFC (positive − neutral) | Diary log NA | 48 | −0.17423 | [−1.28902, 0.94056] | .7539 | .7539 |
| Connectivity and affect / Positive minus neutral | Left amygdala-posterior vmPFC (positive − neutral) | Diary PA | 48 | 0.00035 | [−0.55947, 0.56016] | .9990 | .9990 |
| Connectivity and affect / Positive minus neutral | Left amygdala-posterior vmPFC (positive − neutral) | Diary NA | 48 | 0.06718 | [−0.14712, 0.28148] | .5302 | .5302 |
| Connectivity and affect / Positive minus neutral / log-NA robustness | Left amygdala-posterior vmPFC (positive − neutral) | Diary log NA | 48 | 0.48219 | [−0.73414, 1.69851] | .4280 | .4280 |
| Connectivity and affect / Right amygdala | Right amygdala-anterior vmPFC (negative − neutral) | Diary PA | 48 | 0.01313 | [−0.61730, 0.64355] | .9667 | .9667 |
| Connectivity and affect / Right amygdala | Right amygdala-anterior vmPFC (negative − neutral) | Diary NA | 48 | 0.23351 | [0.00245, 0.46457] | .0477 | .0477 |
| Connectivity and affect / Right amygdala / log-NA robustness | Right amygdala-anterior vmPFC (negative − neutral) | Diary log NA | 48 | 0.95138 | [−0.39603, 2.29880] | .1615 | .1615 |

| **Analysis / specification** | **Predictor / interaction** | **Outcome** | ***N*** | **β** | **95% CI** | ***p* (analysis)** | ***p* (2-sided)** |
|---|---|---|---|---|---|---|---|
| Connectivity and affect / Right amygdala | Right amygdala-posterior vmPFC (negative − neutral) | Diary PA | 48 | −0.16503 | [−0.86411, 0.53404] | .6361 | .6361 |
| Connectivity and affect / Right amygdala | Right amygdala-posterior vmPFC (negative − neutral) | Diary NA | 48 | 0.14435 | [−0.12144, 0.41014] | .2791 | .2791 |
| Connectivity and affect / Right amygdala / log-NA robustness | Right amygdala-posterior vmPFC (negative − neutral) | Diary log NA | 48 | 0.78067 | [−0.73441, 2.29575] | .3042 | .3042 |

### Table S5. M3 main results: OLS and correlation comparisons

| Analysis / specification | Association / interaction | OLS | Pearson correlation |
|---|---|---|---|
| Age and affect | Age (diary) → Diary PA | $N = 1168$; $\beta = 0.00996$<br>95% CI [0.00588, 0.01405]<br>$p$ (analysis) < .001; $p$ (2-sided) < .001 | $N = 1174$; $r = 0.14477$<br>95% CI [0.08829, 0.20033]<br>$p$ (analysis) < .001; $p$ (2-sided) < .001 |
| Age and affect | Age (diary) → Diary NA | $N = 1168$; $\beta = -0.00310$<br>95% CI [−0.00454, −0.00166]<br>$p$ (analysis) < .001; $p$ (2-sided) < .001 | $N = 1174$; $r = -0.11052$<br>95% CI [−0.16668, −0.05365]<br>$p$ (analysis) < .001; $p$ (2-sided) < .001 |
| Persistence and affect | Left negative persistence → Diary PA | $N = 81$; $\beta = -1.60083$<br>95% CI [−2.83030, −0.37135]<br>$p$ (analysis) .0058; $p$ (2-sided) .0115 | $N = 81$; $r = -0.29326$<br>95% CI [−0.48082, −0.08004]<br>$p$ (analysis) .0039; $p$ (2-sided) .0079 |
| Persistence and affect | Left negative persistence → Diary NA | $N = 81$; $\beta = 0.37675$<br>95% CI [0.08008, 0.67342]<br>$p$ (analysis) .0068; $p$ (2-sided) .0136 | $N = 81$; $r = 0.21669$<br>95% CI [−0.00175, 0.41538]<br>$p$ (analysis) .0260; $p$ (2-sided) .0520 |
| Age and persistence | Age (neuroscience) → Left negative persistence | $N = 127$; $\beta = -0.00203$<br>95% CI [−0.00490, 0.00085]<br>$p$ (analysis) .0824; $p$ (2-sided) .1649 | $N = 127$; $r = -0.17607$<br>95% CI [−0.33986, −0.00191]<br>$p$ (analysis) .0238; $p$ (2-sided) .0477 |
| Connectivity and affect | Left amygdala-anterior vmPFC (negative − neutral) → Diary PA | $N = 81$; $\beta = 0.16032$<br>95% CI [−0.34480, 0.66544]<br>$p$ (analysis) .2642; $p$ (2-sided) .5284 | $N = 81$; $r = -0.02228$<br>95% CI [−0.23947, 0.19703]<br>$p$ (analysis) .5783; $p$ (2-sided) .8435 |
| Connectivity and affect | Left amygdala-anterior vmPFC (negative − neutral) → Diary NA | $N = 81$; $\beta = -0.08371$<br>95% CI [−0.20392, 0.03649]<br>$p$ (analysis) .0845; $p$ (2-sided) .1690 | $N = 81$; $r = -0.14858$<br>95% CI [−0.35540, 0.07211]<br>$p$ (analysis) .0928; $p$ (2-sided) .1856 |
| Connectivity and affect | Left amygdala-posterior vmPFC (negative − neutral) → Diary PA | $N = 81$; $\beta = 0.05873$<br>95% CI [−0.45194, 0.56940]<br>$p$ (analysis) .4095; $p$ (2-sided) .8190 | $N = 81$; $r = 0.03223$<br>95% CI [−0.18744, 0.24883]<br>$p$ (analysis) .3876; $p$ (2-sided) .7752 |
| Connectivity and affect | Left amygdala-posterior vmPFC (negative − neutral) → Diary NA | $N = 81$; $\beta = -0.05429$<br>95% CI [−0.17654, 0.06797]<br>$p$ (analysis) .1892; $p$ (2-sided) .3784 | $N = 81$; $r = -0.10591$<br>95% CI [−0.31693, 0.11510]<br>$p$ (analysis) .1733; $p$ (2-sided) .3467 |
| Connectivity and persistence | Left amygdala-anterior vmPFC (negative − neutral) → Left negative persistence | $N = 127$; $\beta = -0.01658$<br>95% CI [−0.09234, 0.05917]<br>$p$ (analysis) .3325; $p$ (2-sided) .6650 | $N = 127$; $r = -0.03135$<br>95% CI [−0.20445, 0.14365]<br>$p$ (analysis) .3632; $p$ (2-sided) .7264 |

| Analysis / specification | Association / interaction | OLS | Pearson correlation |
|---|---|---|---|
| Connectivity and persistence | Left amygdala-posterior vmPFC (negative − neutral) → Left negative persistence | $N = 127$; β = 0.02196<br>95% CI [−0.05162, 0.09554]<br>$p$ (analysis) .7224; $p$ (2-sided) .5552 | $N = 127$; $r = 0.00870$<br>95% CI [−0.16576, 0.18264]<br>$p$ (analysis) .5387; $p$ (2-sided) .9226 |
| Exploratory persistence moderation / reappraisal | Left negative persistence × Reappraisal → Diary PA | $N = 80$; β interaction = 0.97885<br>95% CI [−0.18111, 2.13881]<br>$p$ (analysis) .0967; $p$ (2-sided) .0967 | Not applicable |
| Exploratory persistence moderation / reappraisal | Left negative persistence × Reappraisal → Diary NA | $N = 80$; β interaction = −0.13903<br>95% CI [−0.41519, 0.13713]<br>$p$ (analysis) .3182; $p$ (2-sided) .3182 | Not applicable |
| Exploratory persistence moderation / suppression | Left negative persistence × Suppression → Diary PA | $N = 80$; β interaction = 0.34426<br>95% CI [−0.76159, 1.45011]<br>$p$ (analysis) .5360; $p$ (2-sided) .5360 | Not applicable |
| Exploratory persistence moderation / suppression | Left negative persistence × Suppression → Diary NA | $N = 80$; β interaction = 0.11054<br>95% CI [−0.15732, 0.37841]<br>$p$ (analysis) .4126; $p$ (2-sided) .4126 | Not applicable |
| Exploratory connectivity moderation / reappraisal | Left amygdala-anterior vmPFC (negative − neutral) × Reappraisal → Diary PA | $N = 80$; β interaction = −0.24773<br>95% CI [−0.87566, 0.38020]<br>$p$ (analysis) .4333; $p$ (2-sided) .4333 | Not applicable |
| Exploratory connectivity moderation / reappraisal | Left amygdala-anterior vmPFC (negative − neutral) × Reappraisal → Diary NA | $N = 80$; β interaction = 0.05817<br>95% CI [−0.08935, 0.20570]<br>$p$ (analysis) .4336; $p$ (2-sided) .4336 | Not applicable |
| Exploratory connectivity moderation / reappraisal | Left amygdala-posterior vmPFC (negative − neutral) × Reappraisal → Diary PA | $N = 80$; β interaction = 0.05342<br>95% CI [−0.48520, 0.59203]<br>$p$ (analysis) .8435; $p$ (2-sided) .8435 | Not applicable |
| Exploratory connectivity moderation / reappraisal | Left amygdala-posterior vmPFC (negative − neutral) × Reappraisal → Diary NA | $N = 80$; β interaction = 0.02210<br>95% CI [−0.10392, 0.14811]<br>$p$ (analysis) .7271; $p$ (2-sided) .7271 | Not applicable |
| Exploratory connectivity moderation / suppression | Left amygdala-anterior vmPFC (negative − neutral) × Suppression → Diary PA | $N = 80$; β interaction = 0.26279<br>95% CI [−0.18308, 0.70867]<br>$p$ (analysis) .2432; $p$ (2-sided) .2432 | Not applicable |

| Analysis / specification | Association / interaction | OLS | Pearson correlation |
|---|---|---|---|
| Exploratory connectivity moderation / suppression | Left amygdala-anterior vmPFC (negative − neutral) × Suppression → Diary NA | $N = 80$; β interaction = −0.02246<br>95% CI [−0.12962, 0.08470]<br>*p* (analysis) .6767; *p* (2-sided) .6767 | Not applicable |
| Exploratory connectivity moderation / suppression | Left amygdala-posterior vmPFC (negative − neutral) × Suppression → Diary PA | $N = 80$; β interaction = −0.22139<br>95% CI [−0.69664, 0.25386]<br>*p* (analysis) .3554; *p* (2-sided) .3554 | Not applicable |
| Exploratory connectivity moderation / suppression | Left amygdala-posterior vmPFC (negative − neutral) × Suppression → Diary NA | $N = 80$; β interaction = 0.09287<br>95% CI [−0.01948, 0.20522]<br>*p* (analysis) .1035; *p* (2-sided) .1035 | Not applicable |

### Table S6. MR1 main results: OLS and correlation comparisons

| Analysis / specification | Association / interaction | OLS | Pearson correlation |
|---|---|---|---|
| Persistence and affect | Left negative persistence → Diary PA | *N* = 48; β = −0.19538<br>95% CI [−1.36712, 0.97635]<br>*p* (analysis) .3690; *p* (2-sided) .7380 | *N* = 48; *r* = −0.00922<br>95% CI [−0.29259, 0.27564]<br>*p* (analysis) .4752; *p* (2-sided) .9504 |
| Persistence and affect | Left negative persistence → Diary NA | *N* = 48; β = 0.31133<br>95% CI [−0.12922, 0.75188]<br>*p* (analysis) .0806; *p* (2-sided) .1611 | *N* = 48; *r* = 0.19939<br>95% CI [−0.08984, 0.45760]<br>*p* (analysis) .0871; *p* (2-sided) .1743 |
| Age and persistence | Age (neuroscience) → Left negative persistence | *N* = 115; β = 0.00134<br>95% CI [−0.00157, 0.00425]<br>*p* (analysis) .8177; *p* (2-sided) .3646 | *N* = 115; *r* = 0.06877<br>95% CI [−0.11580, 0.24875]<br>*p* (analysis) .7674; *p* (2-sided) .4652 |
| Connectivity and affect | Left amygdala-anterior vmPFC (negative − neutral) → Diary PA | *N* = 48; β = 0.05918<br>95% CI [−0.47246, 0.59083]<br>*p* (analysis) .4116; *p* (2-sided) .8232 | *N* = 48; *r* = 0.00648<br>95% CI [−0.27817, 0.29008]<br>*p* (analysis) .4826; *p* (2-sided) .9651 |
| Connectivity and affect | Left amygdala-anterior vmPFC (negative − neutral) → Diary NA | *N* = 48; β = −0.08013<br>95% CI [−0.28321, 0.12294]<br>*p* (analysis) .2150; *p* (2-sided) .4301 | *N* = 48; *r* = −0.05122<br>95% CI [−0.33055, 0.23635]<br>*p* (analysis) .3648; *p* (2-sided) .7295 |
| Connectivity and affect | Left amygdala-posterior vmPFC (negative − neutral) → Diary PA | *N* = 48; β = −0.14926<br>95% CI [−0.75131, 0.45279]<br>*p* (analysis) .6904; *p* (2-sided) .6193 | *N* = 48; *r* = −0.06617<br>95% CI [−0.34384, 0.22215]<br>*p* (analysis) .6725; *p* (2-sided) .6550 |
| Connectivity and affect | Left amygdala-posterior vmPFC (negative − neutral) → Diary NA | *N* = 48; β = 0.02214<br>95% CI [−0.21006, 0.25433]<br>*p* (analysis) .5759; *p* (2-sided) .8483 | *N* = 48; *r* = 0.04452<br>95% CI [−0.24269, 0.32455]<br>*p* (analysis) .6181; *p* (2-sided) .7638 |
| Connectivity and persistence | Left amygdala-anterior vmPFC (negative − neutral) → Left negative persistence | *N* = 115; β = 0.03129<br>95% CI [−0.06716, 0.12974]<br>*p* (analysis) .7350; *p* (2-sided) .5300 | *N* = 115; *r* = 0.06432<br>95% CI [−0.12021, 0.24455]<br>*p* (analysis) .7527; *p* (2-sided) .4947 |
| Connectivity and persistence | Left amygdala-posterior vmPFC (negative − neutral) → Left negative persistence | *N* = 115; β = −0.01751<br>95% CI [−0.11901, 0.08399]<br>*p* (analysis) .3665; *p* (2-sided) .7330 | *N* = 115; *r* = −0.02788<br>95% CI [−0.20992, 0.15603]<br>*p* (analysis) .3837; *p* (2-sided) .7674 |

### Table S7. M3 sensitivity results: OLS and correlation comparisons

OLS results cannot resolve unavailable inference in an MLM. Motion-adjusted models and moderation have no zero-order correlation counterpart.

| Analysis / specification | Association / interaction | OLS | Pearson correlation |
|---|---|---|---|
| Age and affect / log-NA robustness | Age (diary) → Diary log NA | $N = 1168$; $\beta = -0.02338$<br>95% CI [−0.03199, −0.01478]<br>$p$ (analysis) < .001; $p$ (2-sided) < .001 | $N = 1174$; $r = -0.14670$<br>95% CI [−0.20221, −0.09024]<br>$p$ (analysis) < .001; $p$ (2-sided) < .001 |
| Age and affect / Neuroscience subsample | Age (neuroscience) → Diary PA | $N = 137$; $\beta = 0.01240$<br>95% CI [−0.00205, 0.02685]<br>$p$ (analysis) .0459; $p$ (2-sided) .0918 | $N = 137$; $r = 0.09897$<br>95% CI [−0.06990, 0.26233]<br>$p$ (analysis) .1249; $p$ (2-sided) .2499 |
| Age and affect / Neuroscience subsample | Age (neuroscience) → Diary NA | $N = 137$; $\beta = -0.00328$<br>95% CI [−0.00892, 0.00236]<br>$p$ (analysis) .1256; $p$ (2-sided) .2512 | $N = 137$; $r = -0.07276$<br>95% CI [−0.23758, 0.09612]<br>$p$ (analysis) .1991; $p$ (2-sided) .3981 |
| Age and affect / Neuroscience subsample / log-NA robustness | Age (neuroscience) → Diary log NA | $N = 137$; $\beta = -0.04397$<br>95% CI [−0.07583, −0.01212]<br>$p$ (analysis) .0036; $p$ (2-sided) .0072 | $N = 137$; $r = -0.18309$<br>95% CI [−0.34035, −0.01586]<br>$p$ (analysis) .0161; $p$ (2-sided) .0322 |
| Age and affect / PANAS | Age (neuroscience) → PANAS PA | $N = 230$; $\beta = 0.00212$<br>95% CI [−0.00999, 0.01424]<br>$p$ (analysis) .3651; $p$ (2-sided) .7302 | $N = 230$; $r = 0.00315$<br>95% CI [−0.12626, 0.13246]<br>$p$ (analysis) .4810; $p$ (2-sided) .9621 |
| Age and affect / PANAS | Age (neuroscience) → PANAS NA | $N = 230$; $\beta = -0.00553$<br>95% CI [−0.01309, 0.00203]<br>$p$ (analysis) .0753; $p$ (2-sided) .1505 | $N = 230$; $r = -0.11821$<br>95% CI [−0.24384, 0.01132]<br>$p$ (analysis) .0368; $p$ (2-sided) .0736 |
| Age and affect / PANAS / log-NA robustness | Age (neuroscience) → PANAS log NA | $N = 230$; $\beta = -0.00414$<br>95% CI [−0.00872, 0.00044]<br>$p$ (analysis) .0380; $p$ (2-sided) .0760 | $N = 230$; $r = -0.14396$<br>95% CI [−0.26832, −0.01488]<br>$p$ (analysis) .0145; $p$ (2-sided) .0291 |
| Persistence and affect / log-NA robustness | Left negative persistence → Diary log NA | $N = 81$; $\beta = 2.62129$<br>95% CI [−0.00429, 5.24688]<br>$p$ (analysis) .0252; $p$ (2-sided) .0504 | $N = 81$; $r = 0.18882$<br>95% CI [−0.03080, 0.39105]<br>$p$ (analysis) .0457; $p$ (2-sided) .0914 |
| Persistence and affect / Positive persistence | Left positive persistence → Diary PA | $N = 81$; $\beta = 0.09824$<br>95% CI [−1.23250, 1.42897]<br>$p$ (analysis) .8832; $p$ (2-sided) .8832 | $N = 81$; $r = 0.01264$<br>95% CI [−0.20628, 0.23035]<br>$p$ (analysis) .9108; $p$ (2-sided) .9108 |

| Analysis / specification | Association / interaction | OLS | Pearson correlation |
|---|---|---|---|
| Persistence and affect / Positive persistence | Left positive persistence → Diary NA | *N* = 81; β = 0.01141<br>95% CI [−0.30900, 0.33182]<br>*p* (analysis) .9435; *p* (2-sided) .9435 | *N* = 81; *r* = −0.02991<br>95% CI [−0.24664, 0.18968]<br>*p* (analysis) .7910; *p* (2-sided) .7910 |
| Persistence and affect / Positive persistence / log-NA robustness | Left positive persistence → Diary log NA | *N* = 81; β = −0.93405<br>95% CI [−3.71100, 1.84290]<br>*p* (analysis) .5041; *p* (2-sided) .5041 | *N* = 81; *r* = −0.11776<br>95% CI [−0.32768, 0.10325]<br>*p* (analysis) .2951; *p* (2-sided) .2951 |
| Persistence and affect / PANAS | Left negative persistence → PANAS PA | *N* = 127; β = −0.74941<br>95% CI [−1.62637, 0.12756]<br>*p* (analysis) .0466; *p* (2-sided) .0931 | *N* = 127; *r* = −0.17660<br>95% CI [−0.34034, −0.00246]<br>*p* (analysis) .0235; *p* (2-sided) .0470 |
| Persistence and affect / PANAS | Left negative persistence → PANAS NA | *N* = 127; β = 0.00069<br>95% CI [−0.64804, 0.64941]<br>*p* (analysis) .4992; *p* (2-sided) .9983 | *N* = 127; *r* = 0.02849<br>95% CI [−0.14645, 0.20171]<br>*p* (analysis) .3752; *p* (2-sided) .7505 |
| Persistence and affect / PANAS / log-NA robustness | Left negative persistence → PANAS log NA | *N* = 127; β = 0.04272<br>95% CI [−0.32945, 0.41489]<br>*p* (analysis) .4102; *p* (2-sided) .8203 | *N* = 127; *r* = 0.06198<br>95% CI [−0.11346, 0.23368]<br>*p* (analysis) .2444; *p* (2-sided) .4888 |
| Persistence and affect / Right hemisphere | Right negative persistence → Diary PA | *N* = 81; β = −0.61435<br>95% CI [−1.95692, 0.72822]<br>*p* (analysis) .3642; *p* (2-sided) .3642 | *N* = 81; *r* = −0.03854<br>95% CI [−0.25475, 0.18133]<br>*p* (analysis) .7326; *p* (2-sided) .7326 |
| Persistence and affect / Right hemisphere | Right negative persistence → Diary NA | *N* = 81; β = −0.03142<br>95% CI [−0.35661, 0.29377]<br>*p* (analysis) .8476; *p* (2-sided) .8476 | *N* = 81; *r* = −0.05297<br>95% CI [−0.26821, 0.16732]<br>*p* (analysis) .6386; *p* (2-sided) .6386 |
| Persistence and affect / Right hemisphere / log-NA robustness | Right negative persistence → Diary log NA | *N* = 81; β = 0.88425<br>95% CI [−1.93614, 3.70464]<br>*p* (analysis) .5334; *p* (2-sided) .5334 | *N* = 81; *r* = 0.05961<br>95% CI [−0.16083, 0.27439]<br>*p* (analysis) .5971; *p* (2-sided) .5971 |
| Persistence and affect / Regional activation | Left amygdala neg image → Diary PA | *N* = 81; β = 0.55455<br>95% CI [−1.00840, 2.11750]<br>*p* (analysis) .4811; *p* (2-sided) .4811 | *N* = 81; *r* = 0.10988<br>95% CI [−0.11113, 0.32054]<br>*p* (analysis) .3288; *p* (2-sided) .3288 |
| Persistence and affect / Regional activation | Left amygdala neg image → Diary NA | *N* = 81; β = −0.11592<br>95% CI [−0.49255, 0.26071]<br>*p* (analysis) .5409; *p* (2-sided) .5409 | *N* = 81; *r* = −0.04899<br>95% CI [−0.26451, 0.17119]<br>*p* (analysis) .6641; *p* (2-sided) .6641 |

| Analysis / specification | Association / interaction | OLS | Pearson correlation |
|---|---|---|---|
| Persistence and affect / Regional activation / log-NA robustness | Left amygdala neg image → Diary log NA | $N = 81$; β = −0.81114<br>95% CI [−4.08990, 2.46762]<br>$p$ (analysis) .6229; $p$ (2-sided) .6229 | $N = 81$; $r = −0.03672$<br>95% CI [−0.25304, 0.18310]<br>$p$ (analysis) .7448; $p$ (2-sided) .7448 |
| Persistence and affect / Regional activation | anterior vmPFC neg image → Diary PA | $N = 81$; β = 0.30514<br>95% CI [−0.92929, 1.53958]<br>$p$ (analysis) .6232; $p$ (2-sided) .6232 | $N = 81$; $r = 0.10201$<br>95% CI [−0.11899, 0.31338]<br>$p$ (analysis) .3649; $p$ (2-sided) .3649 |
| Persistence and affect / Regional activation | anterior vmPFC neg image → Diary NA | $N = 81$; β = −0.02387<br>95% CI [−0.32155, 0.27381]<br>$p$ (analysis) .8733; $p$ (2-sided) .8733 | $N = 81$; $r = 0.04446$<br>95% CI [−0.17559, 0.26029]<br>$p$ (analysis) .6935; $p$ (2-sided) .6935 |
| Persistence and affect / Regional activation / log-NA robustness | anterior vmPFC neg image → Diary log NA | $N = 81$; β = 0.44765<br>95% CI [−2.13930, 3.03461]<br>$p$ (analysis) .7308; $p$ (2-sided) .7308 | $N = 81$; $r = 0.05332$<br>95% CI [−0.16697, 0.26855]<br>$p$ (analysis) .6364; $p$ (2-sided) .6364 |
| Persistence and affect / Regional activation | posterior vmPFC neg image → Diary PA | $N = 81$; β = 0.90100<br>95% CI [−0.40219, 2.20419]<br>$p$ (analysis) .1721; $p$ (2-sided) .1721 | $N = 81$; $r = 0.12938$<br>95% CI [−0.09156, 0.33817]<br>$p$ (analysis) .2497; $p$ (2-sided) .2497 |
| Persistence and affect / Regional activation | posterior vmPFC neg image → Diary NA | $N = 81$; β = −0.17087<br>95% CI [−0.48635, 0.14461]<br>$p$ (analysis) .2834; $p$ (2-sided) .2834 | $N = 81$; $r = −0.06604$<br>95% CI [−0.28035, 0.15453]<br>$p$ (analysis) .5580; $p$ (2-sided) .5580 |
| Persistence and affect / Regional activation / log-NA robustness | posterior vmPFC neg image → Diary log NA | $N = 81$; β = 0.37381<br>95% CI [−2.39281, 3.14042]<br>$p$ (analysis) .7881; $p$ (2-sided) .7881 | $N = 81$; $r = 0.06102$<br>95% CI [−0.15945, 0.27570]<br>$p$ (analysis) .5884; $p$ (2-sided) .5884 |
| Persistence and affect / vmPFC persistence | ant vmPFC neg image-to-face persistence (Fisher $z$) → Diary PA | $N = 81$; β = 0.25385<br>95% CI [−0.76532, 1.27303]<br>$p$ (analysis) .6206; $p$ (2-sided) .6206 | $N = 81$; $r = 0.02630$<br>95% CI [−0.19316, 0.24325]<br>$p$ (analysis) .8157; $p$ (2-sided) .8157 |
| Persistence and affect / vmPFC persistence | ant vmPFC neg image-to-face persistence (Fisher $z$) → Diary NA | $N = 81$; β = −0.06482<br>95% CI [−0.31012, 0.18048]<br>$p$ (analysis) .5995; $p$ (2-sided) .5995 | $N = 81$; $r = −0.09660$<br>95% CI [−0.30844, 0.12437]<br>$p$ (analysis) .3909; $p$ (2-sided) .3909 |
| Persistence and affect / vmPFC persistence / log-NA robustness | ant vmPFC neg image-to-face persistence (Fisher $z$) → Diary log NA | $N = 81$; β = −0.49861<br>95% CI [−2.63290, 1.63569]<br>$p$ (analysis) .6424; $p$ (2-sided) .6424 | $N = 81$; $r = −0.02645$<br>95% CI [−0.24339, 0.19302]<br>$p$ (analysis) .8147; $p$ (2-sided) .8147 |

| Analysis / specification | Association / interaction | OLS | Pearson correlation |
|---|---|---|---|
| Persistence and affect / vmPFC persistence | post vmPFC neg image-to-face persistence (Fisher $z$) → Diary PA | $N = 81$; $\beta = -0.30787$<br>95% CI [−1.44893, 0.83318]<br>$p$ (analysis) .5918; $p$ (2-sided) .5918 | $N = 81$; $r = -0.01700$<br>95% CI [−0.23447, 0.20210]<br>$p$ (analysis) .8803; $p$ (2-sided) .8803 |
| Persistence and affect / vmPFC persistence | post vmPFC neg image-to-face persistence (Fisher $z$) → Diary NA | $N = 81$; $\beta = 0.03154$<br>95% CI [−0.24367, 0.30674]<br>$p$ (analysis) .8197; $p$ (2-sided) .8197 | $N = 81$; $r = 0.01270$<br>95% CI [−0.20623, 0.23041]<br>$p$ (analysis) .9104; $p$ (2-sided) .9104 |
| Persistence and affect / vmPFC persistence / log-NA robustness | post vmPFC neg image-to-face persistence (Fisher $z$) → Diary log NA | $N = 81$; $\beta = 1.36420$<br>95% CI [−1.00614, 3.73454]<br>$p$ (analysis) .2546; $p$ (2-sided) .2546 | $N = 81$; $r = 0.12765$<br>95% CI [−0.09330, 0.33662]<br>$p$ (analysis) .2561; $p$ (2-sided) .2561 |
| Age and persistence / Positive persistence | Age (neuroscience) → Left positive persistence | $N = 127$; $\beta = -0.00001$<br>95% CI [−0.00283, 0.00280]<br>$p$ (analysis) .9918; $p$ (2-sided) .9918 | $N = 127$; $r = -0.13593$<br>95% CI [−0.30297, 0.03922]<br>$p$ (analysis) .1276; $p$ (2-sided) .1276 |
| Age and persistence / Positive persistence | Age (neuroscience) → Right positive persistence | $N = 127$; $\beta = 0.00030$<br>95% CI [−0.00231, 0.00290]<br>$p$ (analysis) .8224; $p$ (2-sided) .8224 | $N = 127$; $r = 0.00532$<br>95% CI [−0.16905, 0.17937]<br>$p$ (analysis) .9527; $p$ (2-sided) .9527 |
| Age and persistence / Right hemisphere | Age (neuroscience) → Right negative persistence | $N = 127$; $\beta = 0.00032$<br>95% CI [−0.00254, 0.00317]<br>$p$ (analysis) .8269; $p$ (2-sided) .8269 | $N = 127$; $r = 0.04128$<br>95% CI [−0.13390, 0.21396]<br>$p$ (analysis) .6449; $p$ (2-sided) .6449 |
| Age and persistence / vmPFC persistence | Age (neuroscience) → ant vmPFC neg image-to-face persistence (Fisher $z$) | $N = 127$; $\beta = 0.00061$<br>95% CI [−0.00275, 0.00397]<br>$p$ (analysis) .7191; $p$ (2-sided) .7191 | $N = 127$; $r = -0.05400$<br>95% CI [−0.22609, 0.12136]<br>$p$ (analysis) .5465; $p$ (2-sided) .5465 |
| Age and persistence / vmPFC persistence | Age (neuroscience) → post vmPFC neg image-to-face persistence (Fisher $z$) | $N = 127$; $\beta = 0.00059$<br>95% CI [−0.00246, 0.00364]<br>$p$ (analysis) .7015; $p$ (2-sided) .7015 | $N = 127$; $r = 0.01663$<br>95% CI [−0.15804, 0.19030]<br>$p$ (analysis) .8528; $p$ (2-sided) .8528 |
| Connectivity and affect / log-NA robustness | Left amygdala-anterior vmPFC (negative − neutral) → Diary log NA | $N = 81$; $\beta = -0.99242$<br>95% CI [−2.02436, 0.03952]<br>$p$ (analysis) .0296; $p$ (2-sided) .0592 | $N = 81$; $r = -0.18943$<br>95% CI [−0.39158, 0.03017]<br>$p$ (analysis) .0452; $p$ (2-sided) .0903 |
| Connectivity and affect / log-NA robustness | Left amygdala-posterior vmPFC (negative − neutral) → Diary log NA | $N = 81$; $\beta = -0.41163$<br>95% CI [−1.47636, 0.65310]<br>$p$ (analysis) .2214; $p$ (2-sided) .4429 | $N = 81$; $r = -0.14738$<br>95% CI [−0.35433, 0.07333]<br>$p$ (analysis) .0946; $p$ (2-sided) .1892 |

| Analysis / specification | Association / interaction | OLS | Pearson correlation |
|---|---|---|---|
| Connectivity and affect / Negative condition | Left amygdala-anterior vmPFC neg → Diary PA | $N = 81$; β = 0.19522<br>95% CI [−0.35827, 0.74871]<br>$p$ (analysis) .2418; $p$ (2-sided) .4837 | $N = 81$; $r = 0.00213$<br>95% CI [−0.21632, 0.22038]<br>$p$ (analysis) .4925; $p$ (2-sided) .9849 |
| Connectivity and affect / Negative condition | Left amygdala-anterior vmPFC neg → Diary NA | $N = 81$; β = −0.04185<br>95% CI [−0.17520, 0.09151]<br>$p$ (analysis) .2665; $p$ (2-sided) .5331 | $N = 81$; $r = −0.03170$<br>95% CI [−0.24833, 0.18795]<br>$p$ (analysis) .3894; $p$ (2-sided) .7788 |
| Connectivity and affect / Negative condition / log-NA robustness | Left amygdala-anterior vmPFC neg → Diary log NA | $N = 81$; β = −0.81356<br>95% CI [−1.95921, 0.33209]<br>$p$ (analysis) .0805; $p$ (2-sided) .1609 | $N = 81$; $r = −0.12079$<br>95% CI [−0.33042, 0.10020]<br>$p$ (analysis) .1414; $p$ (2-sided) .2828 |
| Connectivity and affect / Negative condition | Left amygdala-posterior vmPFC neg → Diary PA | $N = 81$; β = 0.31680<br>95% CI [−0.33020, 0.96380]<br>$p$ (analysis) .1659; $p$ (2-sided) .3318 | $N = 81$; $r = 0.07924$<br>95% CI [−0.14156, 0.29253]<br>$p$ (analysis) .2410; $p$ (2-sided) .4820 |
| Connectivity and affect / Negative condition | Left amygdala-posterior vmPFC neg → Diary NA | $N = 81$; β = −0.05094<br>95% CI [−0.20733, 0.10546]<br>$p$ (analysis) .2588; $p$ (2-sided) .5177 | $N = 81$; $r = 0.00301$<br>95% CI [−0.21548, 0.22122]<br>$p$ (analysis) .5107; $p$ (2-sided) .9787 |
| Connectivity and affect / Negative condition / log-NA robustness | Left amygdala-posterior vmPFC neg → Diary log NA | $N = 81$; β = −0.92448<br>95% CI [−2.26965, 0.42069]<br>$p$ (analysis) .0873; $p$ (2-sided) .1746 | $N = 81$; $r = −0.09915$<br>95% CI [−0.31077, 0.12183]<br>$p$ (analysis) .1892; $p$ (2-sided) .3785 |
| Connectivity and affect / PANAS | Left amygdala-anterior vmPFC (negative − neutral) → PANAS PA | $N = 127$; β = −0.04172<br>95% CI [−0.38326, 0.29983]<br>$p$ (analysis) .5955; $p$ (2-sided) .8091 | $N = 127$; $r = −0.13159$<br>95% CI [−0.29895, 0.04362]<br>$p$ (analysis) .9299; $p$ (2-sided) .1403 |
| Connectivity and affect / PANAS | Left amygdala-anterior vmPFC (negative − neutral) → PANAS NA | $N = 127$; β = −0.38494<br>95% CI [−0.62228, −0.14759]<br>$p$ (analysis) < .001; $p$ (2-sided) .0017 | $N = 127$; $r = −0.18053$<br>95% CI [−0.34393, −0.00652]<br>$p$ (analysis) .0211; $p$ (2-sided) .0422 |
| Connectivity and affect / PANAS / log-NA robustness | Left amygdala-anterior vmPFC (negative − neutral) → PANAS log NA | $N = 127$; β = −0.21743<br>95% CI [−0.35385, −0.08102]<br>$p$ (analysis) .0010; $p$ (2-sided) .0021 | $N = 127$; $r = −0.16374$<br>95% CI [−0.32858, 0.01078]<br>$p$ (analysis) .0329; $p$ (2-sided) .0659 |
| Connectivity and affect / PANAS | Left amygdala-posterior vmPFC (negative − neutral) → PANAS PA | $N = 127$; β = 0.14671<br>95% CI [−0.18413, 0.47755]<br>$p$ (analysis) .1906; $p$ (2-sided) .3811 | $N = 127$; $r = 0.04484$<br>95% CI [−0.13040, 0.21735]<br>$p$ (analysis) .3083; $p$ (2-sided) .6167 |

| Analysis / specification | Association / interaction | OLS | Pearson correlation |
|---|---|---|---|
| Connectivity and affect / PANAS | Left amygdala-posterior vmPFC (negative − neutral) → PANAS NA | $N$ = 127; β = −0.37315<br>95% CI [−0.60393, −0.14237]<br>$p$ (analysis) < .001; $p$ (2-sided) .0018 | $N$ = 127; $r$ = −0.20868<br>95% CI [−0.36947, −0.03577]<br>$p$ (analysis) .0093; $p$ (2-sided) .0185 |
| Connectivity and affect / PANAS / log-NA robustness | Left amygdala-posterior vmPFC (negative − neutral) → PANAS log NA | $N$ = 127; β = −0.20272<br>95% CI [−0.33585, −0.06959]<br>$p$ (analysis) .0016; $p$ (2-sided) .0032 | $N$ = 127; $r$ = −0.18406<br>95% CI [−0.34714, −0.01017]<br>$p$ (analysis) .0192; $p$ (2-sided) .0383 |
| Connectivity and affect / Positive minus neutral | Left amygdala-anterior vmPFC (positive − neutral) → Diary PA | $N$ = 81; β = −0.09495<br>95% CI [−0.52681, 0.33691]<br>$p$ (analysis) .6621; $p$ (2-sided) .6621 | $N$ = 81; $r$ = −0.08614<br>95% CI [−0.29887, 0.13475]<br>$p$ (analysis) .4445; $p$ (2-sided) .4445 |
| Connectivity and affect / Positive minus neutral | Left amygdala-anterior vmPFC (positive − neutral) → Diary NA | $N$ = 81; β = −0.01101<br>95% CI [−0.11510, 0.09307]<br>$p$ (analysis) .8333; $p$ (2-sided) .8333 | $N$ = 81; $r$ = −0.05192<br>95% CI [−0.26724, 0.16834]<br>$p$ (analysis) .6453; $p$ (2-sided) .6453 |
| Connectivity and affect / Positive minus neutral / log-NA robustness | Left amygdala-anterior vmPFC (positive − neutral) → Diary log NA | $N$ = 81; β = −0.43488<br>95% CI [−1.33395, 0.46420]<br>$p$ (analysis) .3376; $p$ (2-sided) .3376 | $N$ = 81; $r$ = −0.09500<br>95% CI [−0.30698, 0.12596]<br>$p$ (analysis) .3989; $p$ (2-sided) .3989 |
| Connectivity and affect / Positive minus neutral | Left amygdala-posterior vmPFC (positive − neutral) → Diary PA | $N$ = 81; β = −0.05569<br>95% CI [−0.56110, 0.44971]<br>$p$ (analysis) .8265; $p$ (2-sided) .8265 | $N$ = 81; $r$ = 0.00944<br>95% CI [−0.20934, 0.22732]<br>$p$ (analysis) .9333; $p$ (2-sided) .9333 |
| Connectivity and affect / Positive minus neutral | Left amygdala-posterior vmPFC (positive − neutral) → Diary NA | $N$ = 81; β = −0.02074<br>95% CI [−0.14235, 0.10087]<br>$p$ (analysis) .7345; $p$ (2-sided) .7345 | $N$ = 81; $r$ = −0.08501<br>95% CI [−0.29783, 0.13586]<br>$p$ (analysis) .4505; $p$ (2-sided) .4505 |
| Connectivity and affect / Positive minus neutral / log-NA robustness | Left amygdala-posterior vmPFC (positive − neutral) → Diary log NA | $N$ = 81; β = −0.18233<br>95% CI [−1.23991, 0.87525]<br>$p$ (analysis) .7317; $p$ (2-sided) .7317 | $N$ = 81; $r$ = −0.07970<br>95% CI [−0.29295, 0.14111]<br>$p$ (analysis) .4794; $p$ (2-sided) .4794 |
| Connectivity and affect / Right amygdala | Right amygdala-anterior vmPFC (negative − neutral) → Diary PA | $N$ = 81; β = 0.06854<br>95% CI [−0.46705, 0.60412]<br>$p$ (analysis) .7991; $p$ (2-sided) .7991 | $N$ = 81; $r$ = 0.06220<br>95% CI [−0.15830, 0.27679]<br>$p$ (analysis) .5812; $p$ (2-sided) .5812 |
| Connectivity and affect / Right amygdala | Right amygdala-anterior vmPFC (negative − neutral) → Diary NA | $N$ = 81; β = 0.04457<br>95% CI [−0.08396, 0.17310]<br>$p$ (analysis) .4911; $p$ (2-sided) .4911 | $N$ = 81; $r$ = 0.04307<br>95% CI [−0.17694, 0.25899]<br>$p$ (analysis) .7026; $p$ (2-sided) .7026 |

| Analysis / specification | Association / interaction | OLS | Pearson correlation |
|---|---|---|---|
| Connectivity and affect / Right amygdala / log-NA robustness | Right amygdala-anterior vmPFC (negative − neutral) → Diary log NA | $N = 81$; β = 0.62507<br>95% CI [−0.48609, 1.73623]<br>$p$ (analysis) .2654; $p$ (2-sided) .2654 | $N = 81$; $r = 0.03896$<br>95% CI [−0.18093, 0.25514]<br>$p$ (analysis) .7298; $p$ (2-sided) .7298 |
| Connectivity and affect / Right amygdala | Right amygdala-posterior vmPFC (negative − neutral) → Diary PA | $N = 81$; β = −0.06408<br>95% CI [−0.64532, 0.51716]<br>$p$ (analysis) .8264; $p$ (2-sided) .8264 | $N = 81$; $r = 0.02975$<br>95% CI [−0.18983, 0.24650]<br>$p$ (analysis) .7920; $p$ (2-sided) .7920 |
| Connectivity and affect / Right amygdala | Right amygdala-posterior vmPFC (negative − neutral) → Diary NA | $N = 81$; β = 0.03896<br>95% CI [−0.10069, 0.17861]<br>$p$ (analysis) .5793; $p$ (2-sided) .5793 | $N = 81$; $r = 0.00067$<br>95% CI [−0.21771, 0.21899]<br>$p$ (analysis) .9953; $p$ (2-sided) .9953 |
| Connectivity and affect / Right amygdala / log-NA robustness | Right amygdala-posterior vmPFC (negative − neutral) → Diary log NA | $N = 81$; β = 0.68183<br>95% CI [−0.52376, 1.88742]<br>$p$ (analysis) .2628; $p$ (2-sided) .2628 | $N = 81$; $r = 0.00481$<br>95% CI [−0.21376, 0.22293]<br>$p$ (analysis) .9660; $p$ (2-sided) .9660 |
| Anterior/posterior contrast / diary | Left amygdala-ant minus posterior vmPFC (negative − neutral) → Diary PA | $N = 81$; β = 0.11503<br>95% CI [−0.42085, 0.65091]<br>$p$ (analysis) .3348; $p$ (2-sided) .6696 | $N = 81$; $r = −0.05481$<br>95% CI [−0.26992, 0.16552]<br>$p$ (analysis) .6865; $p$ (2-sided) .6270 |
| Anterior/posterior contrast / diary | Left amygdala-ant minus posterior vmPFC (negative − neutral) → Diary NA | $N = 81$; β = −0.03400<br>95% CI [−0.16292, 0.09492]<br>$p$ (analysis) .3001; $p$ (2-sided) .6002 | $N = 81$; $r = −0.04342$<br>95% CI [−0.25931, 0.17660]<br>$p$ (analysis) .3502; $p$ (2-sided) .7003 |
| Anterior/posterior contrast / diary / log-NA robustness | Left amygdala-ant minus posterior vmPFC (negative − neutral) → Diary log NA | $N = 81$; β = −0.65905<br>95% CI [−1.77066, 0.45257]<br>$p$ (analysis) .1204; $p$ (2-sided) .2407 | $N = 81$; $r = −0.04297$<br>95% CI [−0.25889, 0.17705]<br>$p$ (analysis) .3517; $p$ (2-sided) .7033 |
| Anterior/posterior contrast / diary | Right amygdala-ant minus posterior vmPFC (negative − neutral) → Diary PA | $N = 81$; β = 0.12600<br>95% CI [−0.41554, 0.66754]<br>$p$ (analysis) .3219; $p$ (2-sided) .6437 | $N = 81$; $r = 0.03298$<br>95% CI [−0.18671, 0.24954]<br>$p$ (analysis) .3850; $p$ (2-sided) .7700 |
| Anterior/posterior contrast / diary | Right amygdala-ant minus posterior vmPFC (negative − neutral) → Diary NA | $N = 81$; β = 0.01176<br>95% CI [−0.11879, 0.14232]<br>$p$ (analysis) .5711; $p$ (2-sided) .8577 | $N = 81$; $r = 0.04167$<br>95% CI [−0.17830, 0.25768]<br>$p$ (analysis) .6441; $p$ (2-sided) .7118 |
| Anterior/posterior contrast / diary / log-NA robustness | Right amygdala-ant minus posterior vmPFC (negative − neutral) → Diary log NA | $N = 81$; β = 0.04712<br>95% CI [−1.08850, 1.18275]<br>$p$ (analysis) .5329; $p$ (2-sided) .9342 | $N = 81$; $r = 0.03373$<br>95% CI [−0.18599, 0.25023]<br>$p$ (analysis) .6175; $p$ (2-sided) .7650 |

| Analysis / specification | Association / interaction | OLS | Pearson correlation |
|---|---|---|---|
| Anterior/posterior contrast / panas | Left amygdala-ant minus posterior vmPFC (negative − neutral) → PANAS PA | *N* = 127; β = −0.20441<br>95% CI [−0.55009, 0.14127]<br>*p* (analysis) .8782; *p* (2-sided) .2435 | *N* = 127; *r* = −0.18807<br>95% CI [−0.35079, −0.01433]<br>*p* (analysis) .9829; *p* (2-sided) .0342 |
| Anterior/posterior contrast / panas | Left amygdala-ant minus posterior vmPFC (negative − neutral) → PANAS NA | *N* = 127; β = 0.01035<br>95% CI [−0.24351, 0.26420]<br>*p* (analysis) .5321; *p* (2-sided) .9357 | *N* = 127; *r* = 0.02765<br>95% CI [−0.14727, 0.20090]<br>*p* (analysis) .6212; *p* (2-sided) .7576 |
| Anterior/posterior contrast / panas / log-NA robustness | Left amygdala-ant minus posterior vmPFC (negative − neutral) → PANAS log NA | *N* = 127; β = −0.00300<br>95% CI [−0.14868, 0.14267]<br>*p* (analysis) .4837; *p* (2-sided) .9675 | *N* = 127; *r* = 0.01957<br>95% CI [−0.15518, 0.19312]<br>*p* (analysis) .5864; *p* (2-sided) .8271 |
| Anterior/posterior contrast / panas | Right amygdala-ant minus posterior vmPFC (negative − neutral) → PANAS PA | *N* = 127; β = −0.21885<br>95% CI [−0.59140, 0.15370]<br>*p* (analysis) .8767; *p* (2-sided) .2466 | *N* = 127; *r* = −0.09295<br>95% CI [−0.26291, 0.08260]<br>*p* (analysis) .8507; *p* (2-sided) .2986 |
| Anterior/posterior contrast / panas | Right amygdala-ant minus posterior vmPFC (negative − neutral) → PANAS NA | *N* = 127; β = −0.10163<br>95% CI [−0.37447, 0.17121]<br>*p* (analysis) .2308; *p* (2-sided) .4617 | *N* = 127; *r* = −0.01128<br>95% CI [−0.18513, 0.16325]<br>*p* (analysis) .4499; *p* (2-sided) .8998 |
| Anterior/posterior contrast / panas / log-NA robustness | Right amygdala-ant minus posterior vmPFC (negative − neutral) → PANAS log NA | *N* = 127; β = −0.06414<br>95% CI [−0.22061, 0.09234]<br>*p* (analysis) .2090; *p* (2-sided) .4181 | *N* = 127; *r* = −0.02140<br>95% CI [−0.19489, 0.15338]<br>*p* (analysis) .4056; *p* (2-sided) .8112 |
| Connectivity and persistence / Negative condition | Left amygdala-anterior vmPFC neg → Left negative persistence | *N* = 127; β = −0.00588<br>95% CI [−0.09169, 0.07993]<br>*p* (analysis) .4461; *p* (2-sided) .8921 | *N* = 127; *r* = 0.00162<br>95% CI [−0.17265, 0.17578]<br>*p* (analysis) .5072; *p* (2-sided) .9856 |
| Connectivity and persistence / Negative condition | Left amygdala-posterior vmPFC neg → Left negative persistence | *N* = 127; β = 0.04687<br>95% CI [−0.04852, 0.14227]<br>*p* (analysis) .8340; *p* (2-sided) .3320 | *N* = 127; *r* = 0.03259<br>95% CI [−0.14243, 0.20564]<br>*p* (analysis) .6420; *p* (2-sided) .7160 |
| Connectivity and persistence / Positive persistence | Left amygdala-anterior vmPFC (negative − neutral) → Left positive persistence | *N* = 127; β = −0.01360<br>95% CI [−0.08780, 0.06060]<br>*p* (analysis) .7169; *p* (2-sided) .7169 | *N* = 127; *r* = −0.05373<br>95% CI [−0.22583, 0.12162]<br>*p* (analysis) .5485; *p* (2-sided) .5485 |
| Connectivity and persistence / Positive persistence | Left amygdala-posterior vmPFC (negative − neutral) → Left positive persistence | *N* = 127; β = −0.02910<br>95% CI [−0.10105, 0.04284]<br>*p* (analysis) .4242; *p* (2-sided) .4242 | *N* = 127; *r* = −0.07869<br>95% CI [−0.24948, 0.09685]<br>*p* (analysis) .3792; *p* (2-sided) .3792 |

| Analysis / specification | Association / interaction | OLS | Pearson correlation |
|---|---|---|---|
| Connectivity and persistence / Positive minus neutral | Left amygdala-anterior vmPFC (positive − neutral) → Left negative persistence | $N = 127$; $\beta = 0.00806$<br>95% CI [−0.06439, 0.08052]<br>$p$ (analysis) .8257; $p$ (2-sided) .8257 | $N = 127$; $r = 0.04792$<br>95% CI [−0.12736, 0.22029]<br>$p$ (analysis) .5927; $p$ (2-sided) .5927 |
| Connectivity and persistence / Positive minus neutral | Left amygdala-posterior vmPFC (positive − neutral) → Left negative persistence | $N = 127$; $\beta = 0.01773$<br>95% CI [−0.05979, 0.09525]<br>$p$ (analysis) .6511; $p$ (2-sided) .6511 | $N = 127$; $r = 0.01816$<br>95% CI [−0.15655, 0.19177]<br>$p$ (analysis) .8394; $p$ (2-sided) .8394 |
| Connectivity and persistence / Right amygdala | Right amygdala-anterior vmPFC (negative − neutral) → Left negative persistence | $N = 127$; $\beta = -0.00443$<br>95% CI [−0.08765, 0.07878]<br>$p$ (analysis) .9161; $p$ (2-sided) .9161 | $N = 127$; $r = -0.05582$<br>95% CI [−0.22782, 0.11956]<br>$p$ (analysis) .5331; $p$ (2-sided) .5331 |
| Connectivity and persistence / Right amygdala | Right amygdala-posterior vmPFC (negative − neutral) → Left negative persistence | $N = 127$; $\beta = -0.00373$<br>95% CI [−0.08525, 0.07779]<br>$p$ (analysis) .9279; $p$ (2-sided) .9279 | $N = 127$; $r = -0.05488$<br>95% CI [−0.22693, 0.12049]<br>$p$ (analysis) .5400; $p$ (2-sided) .5400 |
| Connectivity and persistence / Right hemisphere | Left amygdala-anterior vmPFC (negative − neutral) → Right negative persistence | $N = 127$; $\beta = -0.03278$<br>95% CI [−0.10782, 0.04225]<br>$p$ (analysis) .3882; $p$ (2-sided) .3882 | $N = 127$; $r = -0.09253$<br>95% CI [−0.26251, 0.08303]<br>$p$ (analysis) .3008; $p$ (2-sided) .3008 |
| Connectivity and persistence / Right hemisphere | Left amygdala-posterior vmPFC (negative − neutral) → Right negative persistence | $N = 127$; $\beta = 0.00517$<br>95% CI [−0.06803, 0.07837]<br>$p$ (analysis) .8889; $p$ (2-sided) .8889 | $N = 127$; $r = -0.02372$<br>95% CI [−0.19712, 0.15112]<br>$p$ (analysis) .7912; $p$ (2-sided) .7912 |
| Potential-overlap exclusion subsample | Left negative persistence → Diary PA | $N = 55$; $\beta = -1.26734$<br>95% CI [−2.63794, 0.10325]<br>$p$ (analysis) .0345; $p$ (2-sided) .0690 | $N = 55$; $r = -0.28307$<br>95% CI [−0.51006, −0.01922]<br>$p$ (analysis) .0181; $p$ (2-sided) .0362 |
| Potential-overlap exclusion subsample | Left negative persistence → Diary NA | $N = 55$; $\beta = 0.30801$<br>95% CI [−0.01862, 0.63463]<br>$p$ (analysis) .0319; $p$ (2-sided) .0639 | $N = 55$; $r = 0.20010$<br>95% CI [−0.06885, 0.44194]<br>$p$ (analysis) .0715; $p$ (2-sided) .1430 |
| Potential-overlap exclusion subsample / log-NA robustness | Left negative persistence → Diary log NA | $N = 55$; $\beta = 2.46310$<br>95% CI [−0.39725, 5.32345]<br>$p$ (analysis) .0448; $p$ (2-sided) .0895 | $N = 55$; $r = 0.20368$<br>95% CI [−0.06513, 0.44494]<br>$p$ (analysis) .0679; $p$ (2-sided) .1358 |
| Exploratory persistence moderation / reappraisal / log-NA robustness | Left negative persistence × Reappraisal → Diary log NA | $N = 80$; $\beta$ interaction = −1.08700<br>95% CI [−3.51054, 1.33653]<br>$p$ (analysis) .3734; $p$ (2-sided) .3734 | Not applicable |

| Analysis / specification | Association / interaction | OLS | Pearson correlation |
|---|---|---|---|
| Exploratory persistence moderation / PANAS / reappraisal | Left negative persistence × Reappraisal → PANAS PA | *N* = 126; β interaction = 0.03523<br>95% CI [−0.78355, 0.85401]<br>*p* (analysis) .9321; *p* (2-sided) .9321 | Not applicable |
| Exploratory persistence moderation / PANAS / reappraisal | Left negative persistence × Reappraisal → PANAS NA | *N* = 126; β interaction = −0.17960<br>95% CI [−0.78711, 0.42791]<br>*p* (analysis) .5588; *p* (2-sided) .5588 | Not applicable |
| Exploratory persistence moderation / PANAS / reappraisal / log-NA robustness | Left negative persistence × Reappraisal → PANAS log NA | *N* = 126; β interaction = −0.10983<br>95% CI [−0.45747, 0.23781]<br>*p* (analysis) .5321; *p* (2-sided) .5321 | Not applicable |
| Exploratory persistence moderation / PANAS / suppression | Left negative persistence × Suppression → PANAS PA | *N* = 126; β interaction = 0.36327<br>95% CI [−0.42361, 1.15014]<br>*p* (analysis) .3618; *p* (2-sided) .3618 | Not applicable |
| Exploratory persistence moderation / PANAS / suppression | Left negative persistence × Suppression → PANAS NA | *N* = 126; β interaction = −0.09756<br>95% CI [−0.67576, 0.48064]<br>*p* (analysis) .7385; *p* (2-sided) .7385 | Not applicable |
| Exploratory persistence moderation / PANAS / suppression / log-NA robustness | Left negative persistence × Suppression → PANAS log NA | *N* = 126; β interaction = −0.01450<br>95% CI [−0.34559, 0.31659]<br>*p* (analysis) .9309; *p* (2-sided) .9309 | Not applicable |
| Exploratory persistence moderation / Right hemisphere / reappraisal | Right negative persistence × Reappraisal → Diary PA | *N* = 80; β interaction = 0.63056<br>95% CI [−0.70193, 1.96304]<br>*p* (analysis) .3478; *p* (2-sided) .3478 | Not applicable |
| Exploratory persistence moderation / Right hemisphere / reappraisal | Right negative persistence × Reappraisal → Diary NA | *N* = 80; β interaction = −0.01394<br>95% CI [−0.33193, 0.30405]<br>*p* (analysis) .9305; *p* (2-sided) .9305 | Not applicable |
| Exploratory persistence moderation / Right hemisphere / reappraisal / log-NA robustness | Right negative persistence × Reappraisal → Diary log NA | *N* = 80; β interaction = −1.21923<br>95% CI [−3.92344, 1.48498]<br>*p* (analysis) .3709; *p* (2-sided) .3709 | Not applicable |
| Exploratory persistence moderation / Right hemisphere / suppression | Right negative persistence × Suppression → Diary PA | *N* = 80; β interaction = 0.43697<br>95% CI [−0.84657, 1.72051]<br>*p* (analysis) .4987; *p* (2-sided) .4987 | Not applicable |

| Analysis / specification | Association / interaction | OLS | Pearson correlation |
|---|---|---|---|
| Exploratory persistence moderation / Right hemisphere / suppression | Right negative persistence × Suppression → Diary NA | $N$ = 80; β interaction = −0.08832<br>95% CI [−0.40084, 0.22420]<br>$p$ (analysis) .5742; $p$ (2-sided) .5742 | Not applicable |
| Exploratory persistence moderation / Right hemisphere / suppression / log-NA robustness | Right negative persistence × Suppression → Diary log NA | $N$ = 80; β interaction = −0.06945<br>95% CI [−2.77555, 2.63664]<br>$p$ (analysis) .9592; $p$ (2-sided) .9592 | Not applicable |
| Exploratory persistence moderation / suppression / log-NA robustness | Left negative persistence × Suppression → Diary log NA | $N$ = 80; β interaction = 0.37111<br>95% CI [−2.00985, 2.75206]<br>$p$ (analysis) .7564; $p$ (2-sided) .7564 | Not applicable |
| Exploratory connectivity moderation / reappraisal / log-NA robustness | Left amygdala-anterior vmPFC (negative − neutral) × Reappraisal → Diary log NA | $N$ = 80; β interaction = 0.34014<br>95% CI [−0.90654, 1.58682]<br>$p$ (analysis) .5874; $p$ (2-sided) .5874 | Not applicable |
| Exploratory connectivity moderation / reappraisal / log-NA robustness | Left amygdala-posterior vmPFC (negative − neutral) × Reappraisal → Diary log NA | $N$ = 80; β interaction = 0.20251<br>95% CI [−0.87901, 1.28404]<br>$p$ (analysis) .7095; $p$ (2-sided) .7095 | Not applicable |
| Exploratory connectivity moderation / PANAS / reappraisal | Left amygdala-anterior vmPFC (negative − neutral) × Reappraisal → PANAS PA | $N$ = 126; β interaction = 0.06103<br>95% CI [−0.28979, 0.41185]<br>$p$ (analysis) .7307; $p$ (2-sided) .7307 | Not applicable |
| Exploratory connectivity moderation / PANAS / reappraisal | Left amygdala-anterior vmPFC (negative − neutral) × Reappraisal → PANAS NA | $N$ = 126; β interaction = −0.17171<br>95% CI [−0.41330, 0.06988]<br>$p$ (analysis) .1616; $p$ (2-sided) .1616 | Not applicable |
| Exploratory connectivity moderation / PANAS / reappraisal / log-NA robustness | Left amygdala-anterior vmPFC (negative − neutral) × Reappraisal → PANAS log NA | $N$ = 126; β interaction = −0.09891<br>95% CI [−0.23768, 0.03986]<br>$p$ (analysis) .1604; $p$ (2-sided) .1604 | Not applicable |
| Exploratory connectivity moderation / PANAS / reappraisal | Left amygdala-posterior vmPFC (negative − neutral) × Reappraisal → PANAS PA | $N$ = 126; β interaction = 0.10625<br>95% CI [−0.23112, 0.44362]<br>$p$ (analysis) .5334; $p$ (2-sided) .5334 | Not applicable |
| Exploratory connectivity moderation / PANAS / reappraisal | Left amygdala-posterior vmPFC (negative − neutral) × Reappraisal → PANAS NA | $N$ = 126; β interaction = −0.17246<br>95% CI [−0.40741, 0.06248]<br>$p$ (analysis) .1484; $p$ (2-sided) .1484 | Not applicable |

| Analysis / specification | Association / interaction | OLS | Pearson correlation |
|---|---|---|---|
| Exploratory connectivity moderation / PANAS / reappraisal / log-NA robustness | Left amygdala-posterior vmPFC (negative − neutral) × Reappraisal → PANAS log NA | *N* = 126; β interaction = −0.08044<br>95% CI [−0.21639, 0.05550]<br>*p* (analysis) .2431; *p* (2-sided) .2431 | Not applicable |
| Exploratory connectivity moderation / PANAS / suppression | Left amygdala-anterior vmPFC (negative − neutral) × Suppression → PANAS PA | *N* = 126; β interaction = −0.05427<br>95% CI [−0.35383, 0.24529]<br>*p* (analysis) .7200; *p* (2-sided) .7200 | Not applicable |
| Exploratory connectivity moderation / PANAS / suppression | Left amygdala-anterior vmPFC (negative − neutral) × Suppression → PANAS NA | *N* = 126; β interaction = −0.08601<br>95% CI [−0.29301, 0.12100]<br>*p* (analysis) .4116; *p* (2-sided) .4116 | Not applicable |
| Exploratory connectivity moderation / PANAS / suppression / log-NA robustness | Left amygdala-anterior vmPFC (negative − neutral) × Suppression → PANAS log NA | *N* = 126; β interaction = −0.02279<br>95% CI [−0.14210, 0.09651]<br>*p* (analysis) .7054; *p* (2-sided) .7054 | Not applicable |
| Exploratory connectivity moderation / PANAS / suppression | Left amygdala-posterior vmPFC (negative − neutral) × Suppression → PANAS PA | *N* = 126; β interaction = 0.05344<br>95% CI [−0.25438, 0.36127]<br>*p* (analysis) .7312; *p* (2-sided) .7312 | Not applicable |
| Exploratory connectivity moderation / PANAS / suppression | Left amygdala-posterior vmPFC (negative − neutral) × Suppression → PANAS NA | *N* = 126; β interaction = −0.24552<br>95% CI [−0.45358, −0.03746]<br>*p* (analysis) .0212; *p* (2-sided) .0212 | Not applicable |
| Exploratory connectivity moderation / PANAS / suppression / log-NA robustness | Left amygdala-posterior vmPFC (negative − neutral) × Suppression → PANAS log NA | *N* = 126; β interaction = −0.12220<br>95% CI [−0.24300, −0.00140]<br>*p* (analysis) .0475; *p* (2-sided) .0475 | Not applicable |
| Exploratory connectivity moderation / Right amygdala / reappraisal | Right amygdala-anterior vmPFC (negative − neutral) × Reappraisal → Diary PA | *N* = 80; β interaction = 0.30525<br>95% CI [−0.13377, 0.74427]<br>*p* (analysis) .1695; *p* (2-sided) .1695 | Not applicable |
| Exploratory connectivity moderation / Right amygdala / reappraisal | Right amygdala-anterior vmPFC (negative − neutral) × Reappraisal → Diary NA | *N* = 80; β interaction = −0.03011<br>95% CI [−0.13423, 0.07402]<br>*p* (analysis) .5654; *p* (2-sided) .5654 | Not applicable |
| Exploratory connectivity moderation / Right amygdala / reappraisal / log-NA robustness | Right amygdala-anterior vmPFC (negative − neutral) × Reappraisal → Diary log NA | *N* = 80; β interaction = −0.40189<br>95% CI [−1.28812, 0.48435]<br>*p* (analysis) .3682; *p* (2-sided) .3682 | Not applicable |

| Analysis / specification | Association / interaction | OLS | Pearson correlation |
|---|---|---|---|
| Exploratory connectivity moderation / Right amygdala / reappraisal | Right amygdala-posterior vmPFC (negative − neutral) × Reappraisal → Diary PA | $N$ = 80; β interaction = 0.05042<br>95% CI [−0.62704, 0.72788]<br>$p$ (analysis) .8822; $p$ (2-sided) .8822 | Not applicable |
| Exploratory connectivity moderation / Right amygdala / reappraisal | Right amygdala-posterior vmPFC (negative − neutral) × Reappraisal → Diary NA | $N$ = 80; β interaction = 0.03693<br>95% CI [−0.12201, 0.19587]<br>$p$ (analysis) .6439; $p$ (2-sided) .6439 | Not applicable |
| Exploratory connectivity moderation / Right amygdala / reappraisal / log-NA robustness | Right amygdala-posterior vmPFC (negative − neutral) × Reappraisal → Diary log NA | $N$ = 80; β interaction = −0.44479<br>95% CI [−1.80052, 0.91095]<br>$p$ (analysis) .5144; $p$ (2-sided) .5144 | Not applicable |
| Exploratory connectivity moderation / Right amygdala / suppression | Right amygdala-anterior vmPFC (negative − neutral) × Suppression → Diary PA | $N$ = 80; β interaction = 0.38075<br>95% CI [−0.01786, 0.77936]<br>$p$ (analysis) .0608; $p$ (2-sided) .0608 | Not applicable |
| Exploratory connectivity moderation / Right amygdala / suppression | Right amygdala-anterior vmPFC (negative − neutral) × Suppression → Diary NA | $N$ = 80; β interaction = −0.06253<br>95% CI [−0.16001, 0.03494]<br>$p$ (analysis) .2045; $p$ (2-sided) .2045 | Not applicable |
| Exploratory connectivity moderation / Right amygdala / suppression / log-NA robustness | Right amygdala-anterior vmPFC (negative − neutral) × Suppression → Diary log NA | $N$ = 80; β interaction = −0.40736<br>95% CI [−1.25350, 0.43877]<br>$p$ (analysis) .3396; $p$ (2-sided) .3396 | Not applicable |
| Exploratory connectivity moderation / Right amygdala / suppression | Right amygdala-posterior vmPFC (negative − neutral) × Suppression → Diary PA | $N$ = 80; β interaction = −0.10148<br>95% CI [−0.52642, 0.32346]<br>$p$ (analysis) .6348; $p$ (2-sided) .6348 | Not applicable |
| Exploratory connectivity moderation / Right amygdala / suppression | Right amygdala-posterior vmPFC (negative − neutral) × Suppression → Diary NA | $N$ = 80; β interaction = 0.03526<br>95% CI [−0.06698, 0.13750]<br>$p$ (analysis) .4932; $p$ (2-sided) .4932 | Not applicable |
| Exploratory connectivity moderation / Right amygdala / suppression / log-NA robustness | Right amygdala-posterior vmPFC (negative − neutral) × Suppression → Diary log NA | $N$ = 80; β interaction = 0.13036<br>95% CI [−0.75336, 1.01408]<br>$p$ (analysis) .7691; $p$ (2-sided) .7691 | Not applicable |
| Exploratory connectivity moderation / suppression / log-NA robustness | Left amygdala-anterior vmPFC (negative − neutral) × Suppression → Diary log NA | $N$ = 80; β interaction = 0.16384<br>95% CI [−0.73554, 1.06323]<br>$p$ (analysis) .7170; $p$ (2-sided) .7170 | Not applicable |

| Analysis / specification | Association / interaction | OLS | Pearson correlation |
| --- | --- | --- | --- |
| Exploratory connectivity moderation / suppression / log-NA robustness | Left amygdala-posterior vmPFC (negative − neutral) × Suppression → Diary log NA | $N$ = 80; β interaction = 0.47688<br>95% CI [−0.50733, 1.46110]<br>$p$ (analysis) .3365; $p$ (2-sided) .3365 | Not applicable |
| Motion association / fd mean across runs | fd mean across runs → Left negative persistence | $N$ = 127; β = 0.06939<br>95% CI [−0.19817, 0.33695]<br>$p$ (analysis) .6081; $p$ (2-sided) .6081 | $N$ = 127; $r$ = −0.02531<br>95% CI [−0.19865, 0.14956]<br>$p$ (analysis) .7776; $p$ (2-sided) .7776 |
| Motion association / fd mean across runs | fd mean across runs → Left amygdala-anterior vmPFC (negative − neutral) | $N$ = 127; β = −0.05934<br>95% CI [−0.75665, 0.63797]<br>$p$ (analysis) .8663; $p$ (2-sided) .8663 | $N$ = 127; $r$ = 0.00533<br>95% CI [−0.16904, 0.17938]<br>$p$ (analysis) .9526; $p$ (2-sided) .9526 |
| Motion association / fd mean across runs | fd mean across runs → Left amygdala-posterior vmPFC (negative − neutral) | $N$ = 127; β = 0.15433<br>95% CI [−0.56247, 0.87114]<br>$p$ (analysis) .6702; $p$ (2-sided) .6702 | $N$ = 127; $r$ = 0.09617<br>95% CI [−0.07937, 0.26593]<br>$p$ (analysis) .2821; $p$ (2-sided) .2821 |
| Adjusted for fd mean across runs | Left negative persistence → Diary NA | $N$ = 81; β = 0.34841<br>95% CI [0.04557, 0.65126]<br>$p$ (analysis) .0124; $p$ (2-sided) .0248 | Not applicable |
| Adjusted for fd mean across runs | Age (neuroscience) → Left negative persistence | $N$ = 127; β = −0.00220<br>95% CI [−0.00516, 0.00076]<br>$p$ (analysis) .0718; $p$ (2-sided) .1437 | Not applicable |
| Adjusted for fd mean across runs | Left amygdala-anterior vmPFC (negative − neutral) → Diary NA | $N$ = 81; β = −0.07420<br>95% CI [−0.19499, 0.04659]<br>$p$ (analysis) .1121; $p$ (2-sided) .2242 | Not applicable |
| Adjusted for fd mean across runs / log-NA robustness | Left amygdala-anterior vmPFC (negative − neutral) → Diary log NA | $N$ = 81; β = −0.93282<br>95% CI [−1.97542, 0.10977]<br>$p$ (analysis) .0393; $p$ (2-sided) .0786 | Not applicable |
| Motion association / fd neg mean | fd neg mean → Left negative persistence | $N$ = 127; β = 0.04134<br>95% CI [−0.22070, 0.30338]<br>$p$ (analysis) .7550; $p$ (2-sided) .7550 | $N$ = 127; $r$ = −0.04033<br>95% CI [−0.21305, 0.13483]<br>$p$ (analysis) .6525; $p$ (2-sided) .6525 |
| Motion association / fd neg mean | fd neg mean → Left amygdala-anterior vmPFC (negative − neutral) | $N$ = 127; β = 0.12056<br>95% CI [−0.56147, 0.80259]<br>$p$ (analysis) .7266; $p$ (2-sided) .7266 | $N$ = 127; $r$ = 0.04409<br>95% CI [−0.13114, 0.21664]<br>$p$ (analysis) .6226; $p$ (2-sided) .6226 |

| Analysis / specification | Association / interaction | OLS | Pearson correlation |
|---|---|---|---|
| Motion association / fd neg mean | fd neg mean → Left amygdala-posterior vmPFC (negative − neutral) | $N = 127$; β = 0.26832<br>95% CI [−0.43174, 0.96838]<br>$p$ (analysis) .4488; $p$ (2-sided) .4488 | $N = 127$; $r = 0.13073$<br>95% CI [−0.04450, 0.29816]<br>$p$ (analysis) .1429; $p$ (2-sided) .1429 |
| Adjusted for fd neg mean | Left negative persistence → Diary NA | $N = 81$; β = 0.36827<br>95% CI [0.06745, 0.66910]<br>$p$ (analysis) .0086; $p$ (2-sided) .0172 | Not applicable |
| Adjusted for fd neg mean | Age (neuroscience) → Left negative persistence | $N = 127$; β = −0.00213<br>95% CI [−0.00509, 0.00083]<br>$p$ (analysis) .0783; $p$ (2-sided) .1565 | Not applicable |
| Adjusted for fd neg mean | Left amygdala-anterior vmPFC (negative − neutral) → Diary NA | $N = 81$; β = −0.08216<br>95% CI [−0.20297, 0.03865]<br>$p$ (analysis) .0895; $p$ (2-sided) .1791 | Not applicable |
| Adjusted for fd neg mean / log-NA robustness | Left amygdala-anterior vmPFC (negative − neutral) → Diary log NA | $N = 81$; β = −0.98640<br>95% CI [−2.02662, 0.05382]<br>$p$ (analysis) .0313; $p$ (2-sided) .0627 | Not applicable |

### Table S8. MR1 sensitivity results: OLS and correlation comparisons

| Analysis / specification | Association / interaction | OLS | Pearson correlation |
|---|---|---|---|
| Persistence and affect / log-NA robustness | Left negative persistence → Diary log NA | $N = 48$; β = 1.42413<br>95% CI [−1.10552, 3.95379]<br>*p* (analysis) .1311; *p* (2-sided) .2622 | $N = 48$; $r = 0.15899$<br>95% CI [−0.13107, 0.42397]<br>*p* (analysis) .1402; *p* (2-sided) .2804 |
| Persistence and affect / PANAS | Left negative persistence → PANAS PA | $N = 114$; β = 0.59713<br>95% CI [−0.03547, 1.22973]<br>*p* (analysis) .9680; *p* (2-sided) .0640 | $N = 114$; $r = 0.18019$<br>95% CI [−0.00385, 0.35243]<br>*p* (analysis) .9725; *p* (2-sided) .0551 |
| Persistence and affect / PANAS | Left negative persistence → PANAS NA | $N = 114$; β = 0.12091<br>95% CI [−0.38474, 0.62656]<br>*p* (analysis) .3182; *p* (2-sided) .6364 | $N = 114$; $r = 0.04183$<br>95% CI [−0.14319, 0.22402]<br>*p* (analysis) .3293; *p* (2-sided) .6586 |
| Persistence and affect / PANAS / log-NA robustness | Left negative persistence → PANAS log NA | $N = 114$; β = 0.08719<br>95% CI [−0.21954, 0.39393]<br>*p* (analysis) .2871; *p* (2-sided) .5742 | $N = 114$; $r = 0.05020$<br>95% CI [−0.13496, 0.23198]<br>*p* (analysis) .2979; *p* (2-sided) .5958 |
| Persistence and affect / Right hemisphere | Right negative persistence → Diary PA | $N = 48$; β = −1.28052<br>95% CI [−2.55468, −0.00635]<br>*p* (analysis) .0489; *p* (2-sided) .0489 | $N = 48$; $r = -0.25090$<br>95% CI [−0.49943, 0.03579]<br>*p* (analysis) .0854; *p* (2-sided) .0854 |
| Persistence and affect / Right hemisphere | Right negative persistence → Diary NA | $N = 48$; β = 0.38379<br>95% CI [−0.11593, 0.88352]<br>*p* (analysis) .1286; *p* (2-sided) .1286 | $N = 48$; $r = 0.19483$<br>95% CI [−0.09454, 0.45384]<br>*p* (analysis) .1845; *p* (2-sided) .1845 |
| Persistence and affect / Right hemisphere / log-NA robustness | Right negative persistence → Diary log NA | $N = 48$; β = 1.31623<br>95% CI [−1.58094, 4.21340]<br>*p* (analysis) .3642; *p* (2-sided) .3642 | $N = 48$; $r = 0.13724$<br>95% CI [−0.15286, 0.40556]<br>*p* (analysis) .3523; *p* (2-sided) .3523 |
| Age and persistence / Positive persistence | Age (neuroscience) → Left positive persistence | $N = 115$; β = −0.00034<br>95% CI [−0.00263, 0.00194]<br>*p* (analysis) .7660; *p* (2-sided) .7660 | $N = 115$; $r = -0.08413$<br>95% CI [−0.26319, 0.10053]<br>*p* (analysis) .3713; *p* (2-sided) .3713 |
| Age and persistence / Positive persistence | Age (neuroscience) → Right positive persistence | $N = 115$; β = −0.00027<br>95% CI [−0.00240, 0.00187]<br>*p* (analysis) .8060; *p* (2-sided) .8060 | $N = 115$; $r = -0.03059$<br>95% CI [−0.21251, 0.15338]<br>*p* (analysis) .7455; *p* (2-sided) .7455 |
| Age and persistence / Right hemisphere | Age (neuroscience) → Right negative persistence | $N = 115$; β = −0.00061<br>95% CI [−0.00284, 0.00161]<br>*p* (analysis) .5865; *p* (2-sided) .5865 | $N = 115$; $r = -0.06384$<br>95% CI [−0.24409, 0.12068]<br>*p* (analysis) .4979; *p* (2-sided) .4979 |

| Analysis / specification | Association / interaction | OLS | Pearson correlation |
|---|---|---|---|
| Connectivity and affect / log-NA robustness | Left amygdala-anterior vmPFC (negative − neutral) → Diary log NA | $N = 48$; β = −0.34007<br>95% CI [−1.49995, 0.81982]<br>$p$ (analysis) .2785; $p$ (2-sided) .5570 | $N = 48$; $r = −0.05225$<br>95% CI [−0.33147, 0.23538]<br>$p$ (analysis) .3622; $p$ (2-sided) .7243 |
| Connectivity and affect / log-NA robustness | Left amygdala-posterior vmPFC (negative − neutral) → Diary log NA | $N = 48$; β = 0.38046<br>95% CI [−0.93638, 1.69730]<br>$p$ (analysis) .7186; $p$ (2-sided) .5628 | $N = 48$; $r = 0.08439$<br>95% CI [−0.20465, 0.35990]<br>$p$ (analysis) .7158; $p$ (2-sided) .5685 |
| Connectivity and affect / Negative condition | Left amygdala-anterior vmPFC neg → Diary PA | $N = 48$; β = 0.31251<br>95% CI [−0.41350, 1.03853]<br>$p$ (analysis) .1949; $p$ (2-sided) .3897 | $N = 48$; $r = 0.10435$<br>95% CI [−0.18528, 0.37730]<br>$p$ (analysis) .2401; $p$ (2-sided) .4803 |
| Connectivity and affect / Negative condition | Left amygdala-anterior vmPFC neg → Diary NA | $N = 48$; β = −0.05934<br>95% CI [−0.34056, 0.22188]<br>$p$ (analysis) .3361; $p$ (2-sided) .6722 | $N = 48$; $r = −0.02647$<br>95% CI [−0.30829, 0.25962]<br>$p$ (analysis) .4291; $p$ (2-sided) .8583 |
| Connectivity and affect / Negative condition / log-NA robustness | Left amygdala-anterior vmPFC neg → Diary log NA | $N = 48$; β = −0.96107<br>95% CI [−2.53647, 0.61433]<br>$p$ (analysis) .1125; $p$ (2-sided) .2250 | $N = 48$; $r = −0.15691$<br>95% CI [−0.42222, 0.13316]<br>$p$ (analysis) .1434; $p$ (2-sided) .2869 |
| Connectivity and affect / Negative condition | Left amygdala-posterior vmPFC neg → Diary PA | $N = 48$; β = 0.24936<br>95% CI [−0.43862, 0.93735]<br>$p$ (analysis) .2342; $p$ (2-sided) .4683 | $N = 48$; $r = 0.15270$<br>95% CI [−0.13740, 0.41867]<br>$p$ (analysis) .1501; $p$ (2-sided) .3001 |
| Connectivity and affect / Negative condition | Left amygdala-posterior vmPFC neg → Diary NA | $N = 48$; β = −0.18924<br>95% CI [−0.44885, 0.07036]<br>$p$ (analysis) .0743; $p$ (2-sided) .1486 | $N = 48$; $r = −0.24568$<br>95% CI [−0.49524, 0.04134]<br>$p$ (analysis) .0462; $p$ (2-sided) .0923 |
| Connectivity and affect / Negative condition / log-NA robustness | Left amygdala-posterior vmPFC neg → Diary log NA | $N = 48$; β = −0.90237<br>95% CI [−2.39167, 0.58693]<br>$p$ (analysis) .1140; $p$ (2-sided) .2281 | $N = 48$; $r = −0.22206$<br>95% CI [−0.47615, 0.06626]<br>$p$ (analysis) .0646; $p$ (2-sided) .1293 |
| Connectivity and affect / PANAS | Left amygdala-anterior vmPFC (negative − neutral) → PANAS PA | $N = 114$; β = 0.18617<br>95% CI [−0.14272, 0.51505]<br>$p$ (analysis) .1321; $p$ (2-sided) .2643 | $N = 114$; $r = 0.13104$<br>95% CI [−0.05418, 0.30754]<br>$p$ (analysis) .0823; $p$ (2-sided) .1646 |
| Connectivity and affect / PANAS | Left amygdala-anterior vmPFC (negative − neutral) → PANAS NA | $N = 114$; β = −0.03829<br>95% CI [−0.29864, 0.22206]<br>$p$ (analysis) .3856; $p$ (2-sided) .7712 | $N = 114$; $r = −0.05204$<br>95% CI [−0.23372, 0.13315]<br>$p$ (analysis) .2912; $p$ (2-sided) .5824 |

| Analysis / specification | Association / interaction | OLS | Pearson correlation |
|---|---|---|---|
| Connectivity and affect / PANAS / log-NA robustness | Left amygdala-anterior vmPFC (negative − neutral) → PANAS log NA | $N = 114$; $\beta = -0.01087$<br>95% CI [−0.16892, 0.14718]<br>$p$ (analysis) .4459; $p$ (2-sided) .8918 | $N = 114$; $r = -0.04053$<br>95% CI [−0.22278, 0.14446]<br>$p$ (analysis) .3343; $p$ (2-sided) .6686 |
| Connectivity and affect / PANAS | Left amygdala-posterior vmPFC (negative − neutral) → PANAS PA | $N = 114$; $\beta = 0.22603$<br>95% CI [−0.11173, 0.56378]<br>$p$ (analysis) .0937; $p$ (2-sided) .1874 | $N = 114$; $r = 0.10495$<br>95% CI [−0.08052, 0.28340]<br>$p$ (analysis) .1332; $p$ (2-sided) .2664 |
| Connectivity and affect / PANAS | Left amygdala-posterior vmPFC (negative − neutral) → PANAS NA | $N = 114$; $\beta = -0.10144$<br>95% CI [−0.36883, 0.16595]<br>$p$ (analysis) .2268; $p$ (2-sided) .4536 | $N = 114$; $r = -0.07804$<br>95% CI [−0.25825, 0.10741]<br>$p$ (analysis) .2046; $p$ (2-sided) .4092 |
| Connectivity and affect / PANAS / log-NA robustness | Left amygdala-posterior vmPFC (negative − neutral) → PANAS log NA | $N = 114$; $\beta = -0.07360$<br>95% CI [−0.23568, 0.08849]<br>$p$ (analysis) .1850; $p$ (2-sided) .3701 | $N = 114$; $r = -0.09284$<br>95% CI [−0.27211, 0.09266]<br>$p$ (analysis) .1629; $p$ (2-sided) .3259 |
| Connectivity and affect / Positive minus neutral | Left amygdala-anterior vmPFC (positive − neutral) → Diary PA | $N = 48$; $\beta = 0.11567$<br>95% CI [−0.39276, 0.62410]<br>$p$ (analysis) .6483; $p$ (2-sided) .6483 | $N = 48$; $r = 0.06459$<br>95% CI [−0.22365, 0.34244]<br>$p$ (analysis) .6627; $p$ (2-sided) .6627 |
| Connectivity and affect / Positive minus neutral | Left amygdala-anterior vmPFC (positive − neutral) → Diary NA | $N = 48$; $\beta = -0.11839$<br>95% CI [−0.31089, 0.07410]<br>$p$ (analysis) .2213; $p$ (2-sided) .2213 | $N = 48$; $r = -0.15357$<br>95% CI [−0.41940, 0.13652]<br>$p$ (analysis) .2974; $p$ (2-sided) .2974 |
| Connectivity and affect / Positive minus neutral / log-NA robustness | Left amygdala-anterior vmPFC (positive − neutral) → Diary log NA | $N = 48$; $\beta = -0.17423$<br>95% CI [−1.28902, 0.94056]<br>$p$ (analysis) .7539; $p$ (2-sided) .7539 | $N = 48$; $r = -0.04206$<br>95% CI [−0.32234, 0.24501]<br>$p$ (analysis) .7766; $p$ (2-sided) .7766 |
| Connectivity and affect / Positive minus neutral | Left amygdala-posterior vmPFC (positive − neutral) → Diary PA | $N = 48$; $\beta = 0.00035$<br>95% CI [−0.55947, 0.56016]<br>$p$ (analysis) .9990; $p$ (2-sided) .9990 | $N = 48$; $r = -0.09436$<br>95% CI [−0.36862, 0.19500]<br>$p$ (analysis) .5235; $p$ (2-sided) .5235 |
| Connectivity and affect / Positive minus neutral | Left amygdala-posterior vmPFC (positive − neutral) → Diary NA | $N = 48$; $\beta = 0.06718$<br>95% CI [−0.14712, 0.28148]<br>$p$ (analysis) .5302; $p$ (2-sided) .5302 | $N = 48$; $r = 0.13172$<br>95% CI [−0.15834, 0.40085]<br>$p$ (analysis) .3722; $p$ (2-sided) .3722 |
| Connectivity and affect / Positive minus neutral / log-NA robustness | Left amygdala-posterior vmPFC (positive − neutral) → Diary log NA | $N = 48$; $\beta = 0.48219$<br>95% CI [−0.73414, 1.69851]<br>$p$ (analysis) .4280; $p$ (2-sided) .4280 | $N = 48$; $r = 0.17276$<br>95% CI [−0.11712, 0.43552]<br>$p$ (analysis) .2403; $p$ (2-sided) .2403 |

| Analysis / specification | Association / interaction | OLS | Pearson correlation |
|---|---|---|---|
| Connectivity and affect / Right amygdala | Right amygdala-anterior vmPFC (negative − neutral) → Diary PA | $N$ = 48; β = 0.01313<br>95% CI [−0.61730, 0.64355]<br>$p$ (analysis) .9667; $p$ (2-sided) .9667 | $N$ = 48; $r$ = −0.05534<br>95% CI [−0.33422, 0.23245]<br>$p$ (analysis) .7087; $p$ (2-sided) .7087 |
| Connectivity and affect / Right amygdala | Right amygdala-anterior vmPFC (negative − neutral) → Diary NA | $N$ = 48; β = 0.23351<br>95% CI [0.00245, 0.46457]<br>$p$ (analysis) .0477; $p$ (2-sided) .0477 | $N$ = 48; $r$ = 0.19855<br>95% CI [−0.09070, 0.45691]<br>$p$ (analysis) .1761; $p$ (2-sided) .1761 |
| Connectivity and affect / Right amygdala / log-NA robustness | Right amygdala-anterior vmPFC (negative − neutral) → Diary log NA | $N$ = 48; β = 0.95138<br>95% CI [−0.39603, 2.29880]<br>$p$ (analysis) .1615; $p$ (2-sided) .1615 | $N$ = 48; $r$ = 0.20962<br>95% CI [−0.07923, 0.46600]<br>$p$ (analysis) .1527; $p$ (2-sided) .1527 |
| Connectivity and affect / Right amygdala | Right amygdala-posterior vmPFC (negative − neutral) → Diary PA | $N$ = 48; β = −0.16503<br>95% CI [−0.86411, 0.53404]<br>$p$ (analysis) .6361; $p$ (2-sided) .6361 | $N$ = 48; $r$ = −0.13924<br>95% CI [−0.40726, 0.15087]<br>$p$ (analysis) .3453; $p$ (2-sided) .3453 |
| Connectivity and affect / Right amygdala | Right amygdala-posterior vmPFC (negative − neutral) → Diary NA | $N$ = 48; β = 0.14435<br>95% CI [−0.12144, 0.41014]<br>$p$ (analysis) .2791; $p$ (2-sided) .2791 | $N$ = 48; $r$ = 0.10269<br>95% CI [−0.18690, 0.37585]<br>$p$ (analysis) .4874; $p$ (2-sided) .4874 |
| Connectivity and affect / Right amygdala / log-NA robustness | Right amygdala-posterior vmPFC (negative − neutral) → Diary log NA | $N$ = 48; β = 0.78067<br>95% CI [−0.73441, 2.29575]<br>$p$ (analysis) .3042; $p$ (2-sided) .3042 | $N$ = 48; $r$ = 0.16405<br>95% CI [−0.12595, 0.42823]<br>$p$ (analysis) .2652; $p$ (2-sided) .2652 |

### Table S9. Contextual associations: MLM, OLS, and correlations

| Analysis | Association | MLM | OLS | Pearson correlation |
|---|---|---|---|---|
| Diary/PANAS convergence | Diary PA → PANAS PA | *N* = 136; β = 0.50626<br>95% CI [0.36727, 0.64524]<br>*p* (analysis) < .001; *p* (2-sided) < .001 | *N* = 136; β = 0.48611<br>95% CI [0.34092, 0.63130]<br>*p* (analysis) < .001; *p* (2-sided) < .001 | *N* = 136; *r* = 0.51993<br>95% CI [0.38532, 0.63287]<br>*p* (analysis) < .001; *p* (2-sided) < .001 |
| Diary/PANAS convergence | Diary NA → PANAS NA | *N* = 136; β = 0.59621<br>95% CI [0.35856, 0.83385]<br>*p* (analysis) < .001; *p* (2-sided) < .001 | *N* = 136; β = 0.56362<br>95% CI [0.30290, 0.82433]<br>*p* (analysis) < .001; *p* (2-sided) < .001 | *N* = 136; *r* = 0.42424<br>95% CI [0.27559, 0.55308]<br>*p* (analysis) < .001; *p* (2-sided) < .001 |
| Diary/PANAS convergence | Diary log NA → PANAS log NA | *N* = 136; β = 0.06072<br>95% CI [0.03416, 0.08728]<br>*p* (analysis) < .001; *p* (2-sided) < .001 | *N* = 136; β = 0.05722<br>95% CI [0.02713, 0.08731]<br>*p* (analysis) < .001; *p* (2-sided) < .001 | *N* = 136; *r* = 0.38599<br>95% CI [0.23278, 0.52050]<br>*p* (analysis) < .001; *p* (2-sided) < .001 |
| ERQ context / age erq | Age (neuroscience) → Reappraisal | *N* = 229; β = −0.00435<br>95% CI [−0.01916, 0.01046]<br>*p* (analysis) .5651; *p* (2-sided) .5651 | *N* = 229; β = −0.00381<br>95% CI [−0.02241, 0.01480]<br>*p* (analysis) .6869; *p* (2-sided) .6869 | *N* = 229; *r* = −0.04639<br>95% CI [−0.17498, 0.08376]<br>*p* (analysis) .4849; *p* (2-sided) .4849 |
| ERQ context / age erq | Age (neuroscience) → Suppression | *N* = 229; β = 0.01141<br>95% CI [−0.00619, 0.02901]<br>*p* (analysis) .2037; *p* (2-sided) .2037 | *N* = 229; β = 0.01923<br>95% CI [−0.00076, 0.03923]<br>*p* (analysis) .0593; *p* (2-sided) .0593 | *N* = 229; *r* = 0.06400<br>95% CI [−0.06619, 0.19205]<br>*p* (analysis) .3349; *p* (2-sided) .3349 |
| ERQ context / erq affect diary | Reappraisal → Diary PA | *N* = 135; β = 0.00931<br>95% CI [−0.09869, 0.11731]<br>*p* (analysis) .8659; *p* (2-sided) .8659 | *N* = 135; β = 0.00748<br>95% CI [−0.10715, 0.12212]<br>*p* (analysis) .8973; *p* (2-sided) .8973 | *N* = 135; *r* = 0.04091<br>95% CI [−0.12894, 0.20843]<br>*p* (analysis) .6375; *p* (2-sided) .6375 |
| ERQ context / erq affect diary | Reappraisal → Diary NA | *N* = 135; β = 0.03376<br>95% CI [−0.00374, 0.07126]<br>*p* (analysis) .0776; *p* (2-sided) .0776 | *N* = 135; β = 0.02359<br>95% CI [−0.02119, 0.06836]<br>*p* (analysis) .2988; *p* (2-sided) .2988 | *N* = 135; *r* = 0.04453<br>95% CI [−0.12537, 0.21189]<br>*p* (analysis) .6081; *p* (2-sided) .6081 |
| ERQ context / erq affect diary | Reappraisal → Diary log NA | *N* = 135; β = 0.22663<br>95% CI [−0.01412, 0.46738]<br>*p* (analysis) .0650; *p* (2-sided) .0650 | *N* = 135; β = 0.22773<br>95% CI [−0.02414, 0.47961]<br>*p* (analysis) .0759; *p* (2-sided) .0759 | *N* = 135; *r* = 0.14333<br>95% CI [−0.02627, 0.30490]<br>*p* (analysis) .0972; *p* (2-sided) .0972 |
| ERQ context / erq affect diary | Suppression → Diary PA | *N* = 135; β = −0.04761<br>95% CI [−0.15607, 0.06084]<br>*p* (analysis) .3895; *p* (2-sided) .3895 | *N* = 135; β = −0.05908<br>95% CI [−0.17629, 0.05813]<br>*p* (analysis) .3201; *p* (2-sided) .3201 | *N* = 135; *r* = −0.14083<br>95% CI [−0.30258, 0.02882]<br>*p* (analysis) .1033; *p* (2-sided) .1033 |
| ERQ context / erq affect diary | Suppression → Diary NA | *N* = 135; β = −0.00848<br>95% CI [−0.04601, 0.02906]<br>*p* (analysis) .6581; *p* (2-sided) .6581 | *N* = 135; β = −0.00710<br>95% CI [−0.05329, 0.03909]<br>*p* (analysis) .7613; *p* (2-sided) .7613 | *N* = 135; *r* = 0.10966<br>95% CI [−0.06041, 0.27355]<br>*p* (analysis) .2055; *p* (2-sided) .2055 |

| Analysis | Association | MLM | OLS | Pearson correlation |
|---|---|---|---|---|
| ERQ context / erq affect diary | Suppression → Diary log NA | *N* = 135; β = −0.04216<br>95% CI [−0.28481, 0.20048]<br>*p* (analysis) .7334; *p* (2-sided) .7334 | *N* = 135; β = −0.06771<br>95% CI [−0.32978, 0.19436]<br>*p* (analysis) .6097; *p* (2-sided) .6097 | *N* = 135; *r* = 0.02962<br>95% CI [−0.14004, 0.19759]<br>*p* (analysis) .7331; *p* (2-sided) .7331 |
| ERQ context / erq affect panas | Reappraisal → PANAS PA | *N* = 229; β = 0.17806<br>95% CI [0.09419, 0.26194]<br>*p* (analysis) < .001; *p* (2-sided) < .001 | *N* = 229; β = 0.16010<br>95% CI [0.06642, 0.25377]<br>*p* (analysis) < .001; *p* (2-sided) < .001 | *N* = 229; *r* = 0.26532<br>95% CI [0.14051, 0.38183]<br>*p* (analysis) < .001; *p* (2-sided) < .001 |
| ERQ context / erq affect panas | Reappraisal → PANAS NA | *N* = 229; β = −0.00621<br>95% CI [−0.05756, 0.04514]<br>*p* (analysis) .8126; *p* (2-sided) .8126 | *N* = 229; β = 0.00410<br>95% CI [−0.05611, 0.06431]<br>*p* (analysis) .8932; *p* (2-sided) .8932 | *N* = 229; *r* = −0.01492<br>95% CI [−0.14428, 0.11494]<br>*p* (analysis) .8223; *p* (2-sided) .8223 |
| ERQ context / erq affect panas | Reappraisal → PANAS log NA | *N* = 229; β = −0.00248<br>95% CI [−0.03431, 0.02934]<br>*p* (analysis) .8784; *p* (2-sided) .8784 | *N* = 229; β = 0.00517<br>95% CI [−0.03126, 0.04160]<br>*p* (analysis) .7797; *p* (2-sided) .7797 | *N* = 229; *r* = −0.01500<br>95% CI [−0.14436, 0.11487]<br>*p* (analysis) .8214; *p* (2-sided) .8214 |
| ERQ context / erq affect panas | Suppression → PANAS PA | *N* = 229; β = −0.10155<br>95% CI [−0.17734, −0.02575]<br>*p* (analysis) .0086; *p* (2-sided) .0086 | *N* = 229; β = −0.09951<br>95% CI [−0.18817, −0.01085]<br>*p* (analysis) .0280; *p* (2-sided) .0280 | *N* = 229; *r* = −0.16457<br>95% CI [−0.28806, −0.03569]<br>*p* (analysis) .0126; *p* (2-sided) .0126 |
| ERQ context / erq affect panas | Suppression → PANAS NA | *N* = 229; β = 0.05927<br>95% CI [0.01292, 0.10562]<br>*p* (analysis) .0122; *p* (2-sided) .0122 | *N* = 229; β = 0.05938<br>95% CI [0.00405, 0.11471]<br>*p* (analysis) .0356; *p* (2-sided) .0356 | *N* = 229; *r* = 0.16931<br>95% CI [0.04056, 0.29253]<br>*p* (analysis) .0103; *p* (2-sided) .0103 |
| ERQ context / erq affect panas | Suppression → PANAS log NA | *N* = 229; β = 0.03603<br>95% CI [0.00748, 0.06458]<br>*p* (analysis) .0134; *p* (2-sided) .0134 | *N* = 229; β = 0.03490<br>95% CI [0.00139, 0.06841]<br>*p* (analysis) .0413; *p* (2-sided) .0413 | *N* = 229; *r* = 0.16679<br>95% CI [0.03797, 0.29015]<br>*p* (analysis) .0115; *p* (2-sided) .0115 |
| ERQ context / reappraisal suppression | Reappraisal → Suppression | *N* = 229; β = 0.02537<br>95% CI [−0.12190, 0.17263]<br>*p* (analysis) .7357; *p* (2-sided) .7357 | *N* = 229; β = 0.06719<br>95% CI [−0.09060, 0.22497]<br>*p* (analysis) .4019; *p* (2-sided) .4019 | *N* = 229; *r* = −0.03124<br>95% CI [−0.16023, 0.09880]<br>*p* (analysis) .6381; *p* (2-sided) .6381 |

### Table S10. Task-condition means and paired differences

Activation estimates retain the original first-level beta units. Connectivity estimates are Fisher-*z* transformed beta-series correlations; contrasts subtract the second named condition from the first. Mean intervals concern the population mean (or paired mean difference), not the *t* statistic.

| Measure | Region/connection and condition | *N* | Mean / difference | 95% CI | *t* | *p* (2-sided) |
|---|---|---|---|---|---|---|
| Activation: condition mean | Left amygdala negative activation | 127 | 0.04917 | [0.03187, 0.06646] | 5.626 | < .001 |
| Activation: condition mean | Left amygdala neutral activation | 127 | 0.01477 | [−0.00125, 0.03079] | 1.825 | .0704 |
| Activation: condition mean | Left amygdala positive activation | 127 | 0.04008 | [0.02417, 0.05598] | 4.987 | < .001 |
| Activation: condition mean | Right amygdala negative activation | 127 | 0.06153 | [0.04640, 0.07665] | 8.049 | < .001 |
| Activation: condition mean | Right amygdala neutral activation | 127 | 0.03629 | [0.02275, 0.04984] | 5.302 | < .001 |
| Activation: condition mean | Right amygdala positive activation | 127 | 0.04846 | [0.03494, 0.06197] | 7.094 | < .001 |
| Activation: condition mean | Anterior vmPFC negative activation | 127 | −0.03967 | [−0.06183, −0.01751] | −3.543 | < .001 |
| Activation: condition mean | Anterior vmPFC neutral activation | 127 | −0.02889 | [−0.04799, −0.00979] | −2.994 | .0033 |
| Activation: condition mean | Anterior vmPFC positive activation | 127 | −0.01241 | [−0.03164, 0.00683] | −1.276 | .2041 |
| Activation: condition mean | Posterior vmPFC negative activation | 127 | −0.01202 | [−0.03127, 0.00723] | −1.236 | .2187 |
| Activation: condition mean | Posterior vmPFC neutral activation | 127 | −0.01540 | [−0.03454, 0.00374] | −1.592 | .1138 |
| Activation: condition mean | Posterior vmPFC positive activation | 127 | −0.01339 | [−0.03234, 0.00557] | −1.398 | .1647 |
| Activation: paired difference | Left amygdala (negative − neutral) activation | 127 | 0.03440 | [0.01635, 0.05244] | 3.772 | < .001 |
| Activation: paired difference | Left amygdala (negative − positive) activation | 127 | 0.00909 | [−0.00723, 0.02541] | 1.102 | .2726 |

| **Measure** | **Region/connection and condition** | ***N*** | **Mean / difference** | **95% CI** | ***t*** | ***p* (2-sided)** |
|---|---|---|---|---|---|---|
| Activation: paired difference | Right amygdala (negative − neutral) activation | 127 | 0.02523 | [0.00950, 0.04097] | 3.174 | .0019 |
| Activation: paired difference | Right amygdala (negative − positive) activation | 127 | 0.01307 | [−0.00128, 0.02742] | 1.802 | .0740 |
| Activation: paired difference | Anterior vmPFC (negative − neutral) activation | 127 | −0.01078 | [−0.03279, 0.01123] | −0.969 | .3344 |
| Activation: paired difference | Anterior vmPFC (negative − positive) activation | 127 | −0.02726 | [−0.04536, −0.00916] | −2.981 | .0035 |
| Activation: paired difference | Posterior vmPFC (negative − neutral) activation | 127 | 0.00338 | [−0.02204, 0.02879] | 0.263 | .7931 |
| Activation: paired difference | Posterior vmPFC (negative − positive) activation | 127 | 0.00136 | [−0.01809, 0.02082] | 0.139 | .8899 |
| Connectivity: condition mean | Left amygdala-Anterior vmPFC negative | 127 | 0.19153 | [0.13961, 0.24345] | 7.300 | < .001 |
| Connectivity: condition mean | Left amygdala-Anterior vmPFC neutral | 127 | 0.23535 | [0.17197, 0.29872] | 7.349 | < .001 |
| Connectivity: condition mean | Left amygdala-Anterior vmPFC positive | 127 | 0.24285 | [0.18807, 0.29762] | 8.774 | < .001 |
| Connectivity: condition mean | Left amygdala-Posterior vmPFC negative | 127 | 0.08578 | [0.03939, 0.13217] | 3.659 | < .001 |
| Connectivity: condition mean | Left amygdala-Posterior vmPFC neutral | 127 | 0.08092 | [0.02824, 0.13360] | 3.040 | .0029 |
| Connectivity: condition mean | Left amygdala-Posterior vmPFC positive | 127 | 0.12838 | [0.07674, 0.18003] | 4.920 | < .001 |
| Connectivity: condition mean | Right amygdala-Anterior vmPFC negative | 127 | 0.22592 | [0.17734, 0.27450] | 9.204 | < .001 |
| Connectivity: condition mean | Right amygdala-Anterior vmPFC neutral | 127 | 0.25041 | [0.19075, 0.31008] | 8.306 | < .001 |
| Connectivity: condition mean | Right amygdala-Anterior vmPFC positive | 127 | 0.26557 | [0.21107, 0.32007] | 9.643 | < .001 |
| Connectivity: condition mean | Right amygdala-Posterior vmPFC negative | 127 | 0.12065 | [0.06985, 0.17145] | 4.700 | < .001 |

| **Measure** | **Region/connection and condition** | ***N*** | **Mean / difference** | **95% CI** | ***t*** | ***p* (2-sided)** |
|---|---|---|---|---|---|---|
| Connectivity: condition mean | Right amygdala-Posterior vmPFC neutral | 127 | 0.13451 | [0.08148, 0.18754] | 5.019 | < .001 |
| Connectivity: condition mean | Right amygdala-Posterior vmPFC positive | 127 | 0.16912 | [0.11416, 0.22407] | 6.090 | < .001 |
| Connectivity: paired difference | Left amygdala-Anterior vmPFC (negative − neutral) | 127 | −0.04382 | [−0.10516, 0.01752] | −1.414 | .1599 |
| Connectivity: paired difference | Left amygdala-Anterior vmPFC (negative − positive) | 127 | −0.05132 | [−0.11410, 0.01147] | −1.617 | .1083 |
| Connectivity: paired difference | Left amygdala-Posterior vmPFC (negative − neutral) | 127 | 0.00486 | [−0.05581, 0.06553] | 0.159 | .8743 |
| Connectivity: paired difference | Left amygdala-Posterior vmPFC (negative − positive) | 127 | −0.04261 | [−0.10637, 0.02116] | −1.322 | .1885 |
| Connectivity: paired difference | Right amygdala-Anterior vmPFC (negative − neutral) | 127 | −0.02449 | [−0.08023, 0.03124] | −0.870 | .3861 |
| Connectivity: paired difference | Right amygdala-Anterior vmPFC (negative − positive) | 127 | −0.03965 | [−0.09869, 0.01939] | −1.329 | .1862 |
| Connectivity: paired difference | Right amygdala-Posterior vmPFC (negative − neutral) | 127 | −0.01386 | [−0.06844, 0.04072] | −0.502 | .6163 |
| Connectivity: paired difference | Right amygdala-Posterior vmPFC (negative − positive) | 127 | −0.04847 | [−0.10589, 0.00896] | −1.670 | .0974 |

### Table S11. Statistical indirect associations and component paths

a = age–persistence; b = persistence–affect adjusted for age; c′ = direct age–affect; c = total age–affect. Path tests are one-tailed: a < 0; b < 0 for PA and b > 0 for NA; c and c′ > 0 for PA and < 0 for NA. Reported CIs remain two-sided 95% intervals: normal-Wald for paths and percentile bootstrap for indirect effects, based on 2,000 successful family-level draws per specification. Indirect effects were tested in the predicted direction (positive for PA, negative for NA) using one-sided 95% percentile bounds. No indirect effect met the directional significance criterion. Dashes indicate that indirect inference uses bootstrap bounds rather than a path p value. Assessment order varied across participants, so these are statistical indirect associations and are not interpreted causally.

| Age specification | Path | Outcome | *N* | Estimate | 95% CI | *p (1-sided)* |
|---|---|---|---|---|---|---|
| Neuroscience age | a | Left negative persistence | 81 | −0.00132 | [−0.00441, 0.00176] | .1999 |
| Neuroscience age | *b* | Diary PA | 81 | −0.69957 | [−1.63263, 0.23349] | .0708 |
| Neuroscience age | c′ | Diary PA | 81 | 0.00995 | [−0.00572, 0.02561] | .1067 |
| Neuroscience age | c | Diary PA | 81 | 0.01066 | [−0.00548, 0.02680] | .0978 |
| Neuroscience age | *b* | Diary NA | 81 | 0.27793 | [0.01191, 0.54394] | .0203 |
| Neuroscience age | c′ | Diary NA | 81 | −0.00434 | [−0.00807, −0.00062] | .0112 |
| Neuroscience age | c | Diary NA | 81 | −0.00462 | [−0.00841, −0.00082] | .0085 |
| Neuroscience age | *b* | Diary log NA | 81 | 1.95009 | [−0.54517, 4.44534] | .0628 |
| Neuroscience age | c′ | Diary log NA | 81 | −0.05760 | [−0.09064, −0.02456] | < .001 |
| Neuroscience age | c | Diary log NA | 81 | −0.05854 | [−0.09180, −0.02528] | < .001 |
| Diary age | a | Left negative persistence | 81 | −0.00115 | [−0.00423, 0.00194] | .2327 |
| Diary age | *b* | Diary PA | 81 | −0.70978 | [−1.64320, 0.22364] | .0681 |
| Diary age | c′ | Diary PA | 81 | 0.00897 | [−0.00644, 0.02437] | .1270 |
| Diary age | c | Diary PA | 81 | 0.00935 | [−0.00655, 0.02525] | .1246 |
| Diary age | *b* | Diary NA | 81 | 0.28265 | [0.01635, 0.54894] | .0187 |
| Diary age | c′ | Diary NA | 81 | −0.00413 | [−0.00780, −0.00046] | .0137 |
| Diary age | c | Diary NA | 81 | −0.00434 | [−0.00808, −0.00061] | .0113 |
| Diary age | *b* | Diary log NA | 81 | 2.00981 | [−0.48700, 4.50663] | .0573 |
| Diary age | c′ | Diary log NA | 81 | −0.05603 | [−0.08851, −0.02354] | < .001 |
| Diary age | c | Diary log NA | 81 | −0.05665 | [−0.08938, −0.02391] | < .001 |
| Neuroscience age | a × *b* | Diary PA | 81 | 0.00093 | [−0.00134, 0.00516] | — |
| Neuroscience age | a × *b* | Diary NA | 81 | −0.00037 | [−0.00109, 0.00037] | — |
| Neuroscience age | a × *b* | Diary log NA | 81 | −0.00258 | [−0.01041, 0.00237] | — |

| Age specification | Path | Outcome | *N* | Estimate | 95% CI | *p (1-sided)* |
|---|---|---|---|---|---|---|
| Diary age | a × *b* | Diary PA | 81 | 0.00082 | [−0.00149, 0.00483] | — |
| Diary age | a × *b* | Diary NA | 81 | −0.00032 | [−0.00105, 0.00038] | — |
| Diary age | a × *b* | Diary log NA | 81 | −0.00231 | [−0.01037, 0.00284] | — |

**Supplementary Figure S1. Exploratory moderation of primary diary associations**

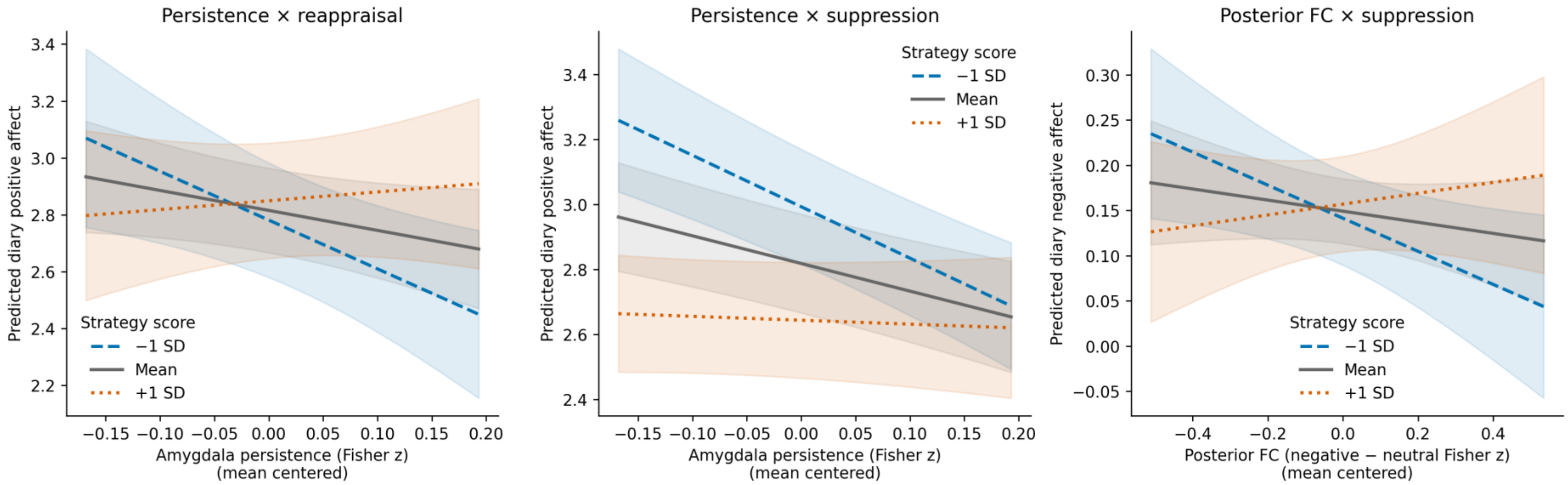


Left: left-amygdala negative persistence × reappraisal predicting positive affect. Middle: persistence × suppression predicting positive affect. Right: left-amygdala–posterior vmPFC connectivity × suppression predicting raw negative affect. All models use the primary QC-retained diary sample ($N = 80$), family random intercepts, and the covariates specified in the manuscript. Lines show fixed-effect predicted means averaged over the model sample's covariate distribution at strategy scores one *SD* below the mean, at the mean, and one *SD* above it; these are evaluation points, not participant subgroups. Shading indicates pointwise two-sided 95% confidence intervals for predicted means, not individual prediction intervals. Predictors are mean centered and displayed over their 5th–95th percentile range. Panels show the three interactions meeting nominal $p < .05$ among 18 primary diary interaction tests, without multiplicity adjustment; all interaction estimates are reported in Supplementary Tables S1 and S3. Attenuation of a slope does not establish beneficial or causal compensation.
